# Dietary macronutrients modulate the proteome of brown adipose tissue in males and their female offspring

**DOI:** 10.1101/2025.02.01.636065

**Authors:** Erin L Macartney, Alistair M Senior, Angela J Crean, Lewin Small, Tamara J Pulpitel, Marcelo A Nobrega, Romain Barrès, Stephen J Simpson

## Abstract

Brown adipose tissue (BAT) dissipates energy as heat, not only under cold exposure but also in the dissipation of excess ingested energy. Therefore, enhancing BAT activity is a potential avenue to combat weight gain. Dietary macronutrient composition influences BAT size and has recently been shown to influence BAT size of daughters through the patriline of C57BL/6J mice. However, the effects of macronutrient composition and any paternal effects on BAT function have yet to be characterised. Using the Geometric Framework for Nutrition, we investigated the effects of macronutrient composition on the BAT proteome in male mice and intergenerational effects in their offspring. In fathers, >50% of the proteome was affected by macronutrient composition, with distinct clusters of proteins that responded in similar ways. We identified two clusters with inverse patterns that correlated with BAT mass. Notably, UCP1 was reduced on low fat diets that promoted increased BAT mass, while there were increased levels of proteins involved in protein turnover on those same diets. The same diets also led to a reduction in proteins involved in purine biosynthesis (often UCP1 inhibitors). We did not find any effects of paternal diet on the BAT proteome in sons, but paternal protein intake negatively affected basigin expression in daughters - a protein that regulates UCP1 transcription. Our results highlight that dietary macronutrient composition in males remodels the protein expression landscape of BAT, and pre-conceptionally reprograms BAT expression profiles of female offspring.

## Introduction

Diet and metabolic health are profoundly linked, yet understanding this link is complex due to the intricate relationship between many tissues, organs, and physiological processes, along with the dynamic and multidimensional nature of nutrition ^1^.

Metabolic phenotypes show heritability through genomic determination ^2,3^, but there is evidence showing that metabolic function can be influenced by ‘non-genetic’ mechanisms reflecting parental nutritional environment and metabolic health ^4–6^.

To overcome some of the challenges associated with the complex and multidimensional nature of nutrition, the Geometric Framework for Nutrition (GFN) provides a powerful tool to unravel the effects of different macronutrients on metabolic outcomes ^1,7^. It does so by investigating the effects of nutrients in multidimensional nutritional space where each macronutrient component represents a dimension and can be manipulated systematically. Recently, two studies on male mice fed one of ten isocaloric diets differing systematically in protein, carbohydrate, and fat ratios reported changes in reproductive and metabolic traits, including some effects of paternal diet on offspring metabolic phenotypes ^8,9^. Fathers exhibited both additive and interactive effects on a range of traits related to metabolism, including liver triglycerides, glucose tolerance, subcutaneous white adipose tissue, and brown adipose tissue. There were also effects of paternal diet on aspects of offspring metabolism in a sex-specific manner. While paternal diet had minimal effects on the metabolism of sons, daughters showed differences in fasting glucose and body fat composition, including weight of brown adipose tissue, a highly metabolically active tissue involved in dissipating energy as heat ^9^. Fathers on low fat diets had daughters with less subcutaneous white fat and larger brown adipose tissue depots than other dietary ratios. This suggests that effects are sex-specific, and that the diet of the fathers influences the metabolic health of their daughters, potentially through brown adipose tissue activity.

Brown adipose tissue (BAT) is a specialised type of fat tissue in mammals that is important for regulating non-shivering thermogenesis and energy expenditure and is implicated in regulating other metabolic phenotypes such as glucose homeostasis and insulin sensitivity ^10^. In mammals, uncoupling protein 1 (UCP1) is thought to be exclusively expressed in BAT where it is involved in dissipating energy as heat^11,12^. BAT is thought to play a crucial role in cold tolerance ^13,14^ but is also influenced by diet^15–17^. For example, high-fat and high-calorie diets in mice have been shown to upregulate UCP1 ^18^ and increase recruitment of BAT tissue ^19^. Such effects are often considered ‘adaptive’ and may act to attenuate weight gain^12,18,20^. Similarly, smaller BAT and loss of BAT function can be associated with increased obesity ^20–23^. Male mice reared on low protein to carbohydrate ratio diets had larger BAT tissues and higher body temperatures ^24^, however, the gene for UCP1 was downregulated in these diets, suggesting that larger BAT size does not necessarily correspond to an increase in UCP1, and that other mechanisms of thermogenesis may be occurring^25^.

Building on the work of Crean et al., (2023)^8^ and Crean et al., (2024)^9^, we conducted total proteomics of whole BAT to examine diet-induced changes in BAT functioning in both F0 males and their F1 offspring. We first conducted a series of hierarchical statistical models in both the F0 males (fathers) and their male and female offspring to generate a ‘set’ of proteins significantly affected by diet. Next, we clustered our significant set of proteins based on their response to the three nutritional dimensions of the diet (protein, carbohydrate, fat) and conducted functional enrichment analyses on the proteins in these clusters. We were then able to explore potentially important proteins found in these clusters and enriched biological processes. These included proteins known to be important for the functioning of BAT, including Ucp1, and ‘hub’ proteins that were highly connected to other proteins in our enriched processes. This study represents the first comprehensive analysis of BAT functioning through total proteomics (avoiding confounding issues of post-transcriptional regulation^26^) across a range of biologically feasible dietary macronutrient compositions, including testing for non-genetic inheritance of BAT function due to paternal diet. Our results have implications for understanding the dietary determinants of metabolic health within and across generations.

## Materials and methods

### Experimental setup

Male (F0) C57BL/6J mice were reared on one of ten isocaloric diets that varied systematically in protein, fat, and carbohydrate concentrations within physiologically feasible ranges (Supp. Mat. 1 Fig. S1). Dietary protein was exome matched ^27^ to the *Mus musculus* genome and the diets were made to be standardised for calories (3.4 ± 0.1 kcal/g; Specialty Feeds, Glen Forrest, Australia) by adjusting the amount of dietary cellulose.

Four-week-old males (F0 males/fathers) were housed in groups of 3 and allowed to acclimate for four days. Cages were randomly allocated to dietary treatments (N = 60 males, n = 6 males/diet). After 12 weeks with *ad libitum* access to their allocated diet, males were allowed to mate with a standardised chow-fed female. Males were kept overnight with females for a maximum of four-nights per ovulation cycle and kept in their dietary treatment cages during the day. Once mating occurred, females remained undisturbed until a litter was born, and fathers were returned to the dietary treatments. Food intake for fathers was calculated by weighing the food provided and the food remaining corrected for spillage over two 24-h periods at weeks 15 and 20.

Fathers were then euthanised by sodium pentobarbital (100 mg/kg) and exsanguination at 22 weeks old (18 weeks total of dietary exposure). BAT was removed and weighed, snap frozen in liquid nitrogen, then stored at -80 degrees Celsius until used for proteomics.

Offspring were raised with their mothers until weaning and then chow fed until they were euthanised at 20 weeks old. Where possible, four offspring per male (two of each sex) were sacrificed.

All mice were housed in standard laboratory temperatures at 22 °C. All procedures were reviewed and approved by the University of Sydney Animal Ethics Committee (project number 2019/1610)

### Proteomics of BAT

1ml of 4% sodium deoxycholate (SDC), 100mM Tris-HCL buffer (pH = 8) and a stainless- steel bead were added to frozen whole BAT samples. Tissues were boiled for 10 min at 95 °C at 1000 rpm in a ThermoMixer (Eppendorf) and homogenised by steel bead disruption for 2 minutes in a bead mill (Retsch). Samples were then boiled for another 10 minutes at 95 °C and 1000 rpm in a ThermoMixer and then centrifuged (Eppendorf) for 10 min at 20,000 g to clarify. The supernatant was then transferred to a new 1.5 ml Eppendorf and stored at -80 °C until used for protein quantification.

Next, blocks of 80 randomly selected samples were defrosted for 1 min at 95 °C and 1000 rpm in a ThermoMixer, then centrifuged for 5 minutes at 20,000 g. We then performed a bicinchoninic protein assay (BCA) (Sigma-Adrich) of the supernatant to quantify the total protein per sample. Based on the BCA, a concentration of 30 µg of protein in 40 µL of 4% SDC buffer was added to each well of a 500ul 96 deep-well plate and then frozen at -30 °C until used for protein digest and cleanup.

96-well plates were defrosted at room temperature, then proteins were reduced and alkylated by adding TCEP (Thermo Fisher) and 2-chloroacetamide (Sigma-Adrich) with a final concentration of 10 mM and 40 mM respectively. Plates were vortexed for 10 minutes at 95 °C at 1000 rpm. The SDC buffer was diluted to 1% by adding 140 µL of MS- grade water. Samples were cooled to 37 °C and Trypsin (Sigma-Adrich) and LysC (Novachem) were added in a 1:50 protease: protein concentration. Samples were then left to digest at 37 °C at1000 rpm for 16 hours in a ThermoMixer.

Digested peptides were diluted 1:1 with 99% ethyl-acetate and 1% trifluoracetic acid (TFA) and centrifuged for 1 minute at 2000 g. 100 µl of the bottom phase was then added to 200 µl equilibrated StageTips with punched SDB-RPS discs and mounted into a 3D-printed holder with a polypropylene waste plate. StageTips were washed twice with 100 µl of 99% ethyl-acetate and 1% TFA, then once with 100 µl of 0.2% TFA. StageTips in the 3D-printed holder were then mounted onto a clean 96-well non-skirted PCR plate

(Thermo Fisher Scientific) and 100 µl of 1.25% ammonium hydroxide and 80% acetonitrile was added. Peptides were then dried in a vacuum concentrator (Genevac) set at a maximum temperature of 45 °C and NH3-H_2_O mode for 2 hours. Peptides were resuspended in 60 µl of 5% formic acid and stored at 5 °C overnight. Peptides were then transferred to 96-well V-bottom plate (Greiner) and covered with a clear silicone micromat (Thermo Fisher scientific) and stored at 4 °C until used for mass spectrometry.

### Mass spectrometry

Peptides in the 96-well plate were placed in the autosampler of a Shimadzu Nexera LC- 40 and were injected (20 μL) onto a 2.1 x 50mm, 1.9 μm, InfinityLab Poroshell 120 EC- C18 analytical column (Agilent). Peptides were resolved over a gradient from 9% acetonitrile to 29% acetonitrile for 13 minutes with a flow rate of 0.8 ml/min and then 80% acetonitrile for 3 minutes at 40 °C. The ZenoTOF 7600 system (Sciex) was operated using the TurbulonSpray ion source in SWATH mode with positive polarity and a spray voltage of 4500 V. The Zeno SWATH DIA method consisted of 60 variable-width SWATH DIA windows that spanned the Q1 mass range 400-900 Da with MS/MS accumulation times of 13 ms.

Zeno SWATH DIA data were processed using DIA-NN (version 1.8.1)^28^ with an in silico generated spectral library, standard settings with “high accuracy”, “robust LC” and “MBR” activated. Peptides were searched against a reference mouse proteome (UP000000589) downloaded from www.uniprot.org on 08/07/2024.

### Data analysis

All analyses were conducted in the R environment (version 4.3.3) using RStudio (version 2023.12.1+402). All data and code can be found at https://osf.io/j9dv4/.

Briefly, our analysis of proteomic data proceeded in three-steps; 1) normalisation and cleaning, 2) a hierarchical statistical screen designed to identify proteins significantly affected by the diet, 3) clustering of significant proteins by their functional response to the GFN dietary design.

First, we filtered out proteins that were missing in more than 30% of the samples. The remaining proteins were then log transformed and normalised from the median (i.e., the median was set to zero).

In the second part of the analysis, F0 males and F1 female and male offspring were separated into different datasets and analysed separately. To test for effects of diet on protein expression in all three datasets to generate our significant ‘set’ of proteins, we used the R package limma ^29^. We fitted a series of statistical models hierarchically. We began by fitting three interaction models between all pairwise combinations of macronutrients (i.e., between percent protein and percent carbohydrate, percent protein and percent fat, and percent carbohydrate and percent fat). This interaction term is equivalent to a quadratic term in a mixture model ^29,30^. We also ran three additive/linear models with all combinations of macronutrients mentioned above.

Adjusted p-values were calculated using the Benjamini-Hochberg false discovery rate.

These six models per dataset then gave us a set of proteins that were significantly influenced by diet after correcting for multiple comparisons and were used in further analyses. If there were no proteins that were significantly affected by diet after correcting for multiple comparisons in any of these six models (e.g., in the male and female offspring analyses), we then reduced the models to three univariate models testing for main effects of individual nutrients (i.e., a model with logged percent protein, percent carbohydrate, or percent fat alone).

Next, we took all the proteins identified as being significantly affected by diet and clustered them based on their response to dietary macronutrients. Each significant protein was scaled using z-transformation before fitting a quadratic mixture model that included the interaction and main effects of all three nutrients (percent) using lme4 (Bates et al., 2015) (∼*0 + protein + carbohydrate + fat + protein:carbohydrate + protein:fat + carbohyrdate:fat*). Using the e1071 package ^32^, proteins were then assigned into a cluster based on their model coefficients and highest membership to a cluster using fuzzy c-means clustering. Our proteins were clustered into five clusters which was determined by finding a balance between the number of clusters identified using the NbClust package ^33^ (note that this uses k-means clustering, not fuzzy c-means), PCA plots, and visual inspection of how well the response of individual proteins to diet fit within the average response of each cluster. The average response of proteins within a cluster was then mapped onto macronutrient space as a right-angled surface with percent protein on the x-axis, percent carbohydrate on the y-axis, and percent fat as the distance from the hypotenuse to the origin ^34^.

To investigate whether the proteins within these clusters fell into shared functions/pathways, we conducted functional enrichment analyses, limited to biological processes, using the package clusterProfiler ^35^. The enriched pathways were simplified to account for redundancy between highly similar pathways using the “wang” method. To investigate the enriched pathways further, we constructed networks of protein-protein interactions between proteins that were involved in significantly enriched biological processes using the STRINGdb package ^36^. For this, we filtered for proteins that had a medium to high score threshold, and proteins that are likely to be ‘hub’ proteins based on having node degrees to five or more other proteins. We then calculated the most highly connected proteins based on the number of node degrees.

All analyses were also run with dietary intake as a co-variate in the model (see Supp. Mat. 2). This is because including dietary intake in the models accounts for differences in how much total food was consumed between the diets, thus removing a potential driver of dietary effects.

Our final sample size of individuals was N = 57 F0 males (one replicate from diets 6, 7, and 8 did not generate usable data from the mass spectrometry output). N = 92 F1 females, 97 = F1 males. Some males failed to produce offspring: diet 5 n = 3, diet 6 n = 1, diet 9 n=3, diet 10 n=1 (Supp. Mat.1; Fig. S1).

## Results

### Body composition and phenotypes

Full phenotypic data of F0 males and offspring can be found in Crean et al., (2024)^9^.

Briefly, F0 males had the highest weight gain on low fat diets, especially on intermediate protein and carbohydrate (note the diets were standardised for calories) (Fig. 1). This pattern was similar to the amount (grams) of subcutaneous white adipose tissue (WAT) ^9^ (Fig. 1). Interscapular BAT, was also highest on low fat diets, especially on low protein, high carbohydrate diets ^9^ (Fig. 1). Additionally, males showed some degree of protein leverage as males had higher food intake on the low protein diet ^9^ (Fig. 1).

**Fig. 1.**
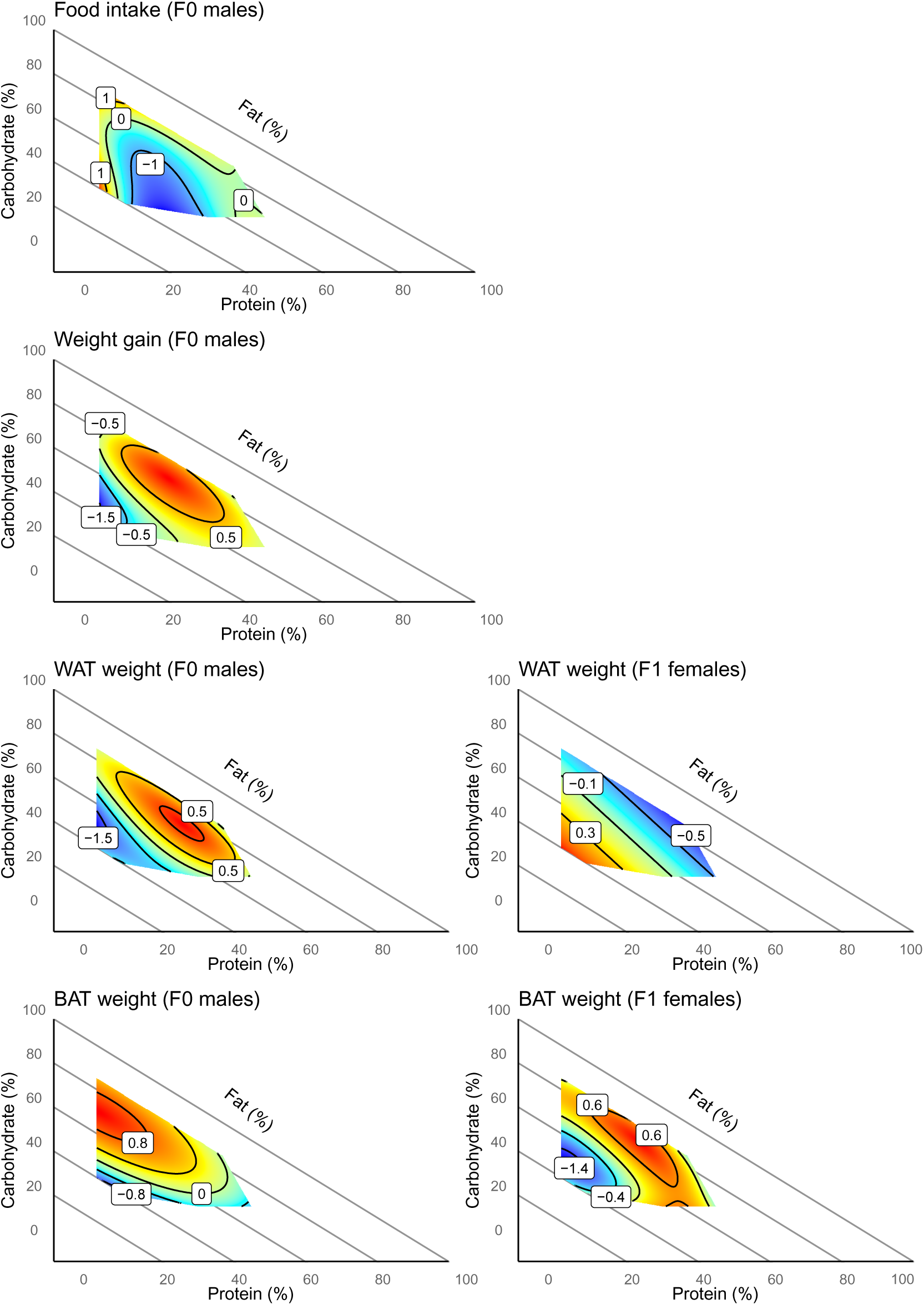
Surface plots of F0 male food intake, F0 male weight gain, F0 male WAT weight, F0 male BAT weight, as well as F1 female offspring WAT weight and F1 female offspring BAT weight across the 10 diets. Figures replotted using the data and models from Crean et al., (2024)^9^. All responses are scaled to z-scores.

In the F1 females, along with differences in WAT weight that showed a strong negative relationship with paternal dietary fat (Fig. 1), and other metabolic phenotypes^9^, paternal diet influenced the interscapular BAT weight of daughters ^9^ (Fig. 1). Specifically, BAT was largest when their fathers were reared on the low-fat diet, but this was especially pronounced on intermediate carbohydrate and protein (i.e., the response was shifted along the protein and carbohydrate axes compared to F0 males/fathers) (Fig. 1).

However, there was no relationship between paternal diet on F1 male offspring BAT weight, or other aspects of adiposity and metabolism^9^.

### Dietary effects on F0 males (fathers)

After filtering for proteins that were missing in >30% of samples, our dataset included 2582 individual proteins in BAT. We detected 1385 proteins that were significantly affected by diet which were used for further analyses (Supp Mat. 1, Fig. S2). These proteins were clustered into five response types (i.e., average response surface topologies), all with relatively equal numbers of proteins (Fig. 2; min = 239 proteins in cluster 3, max = 362 proteins in cluster 4).

**Fig. 2.**
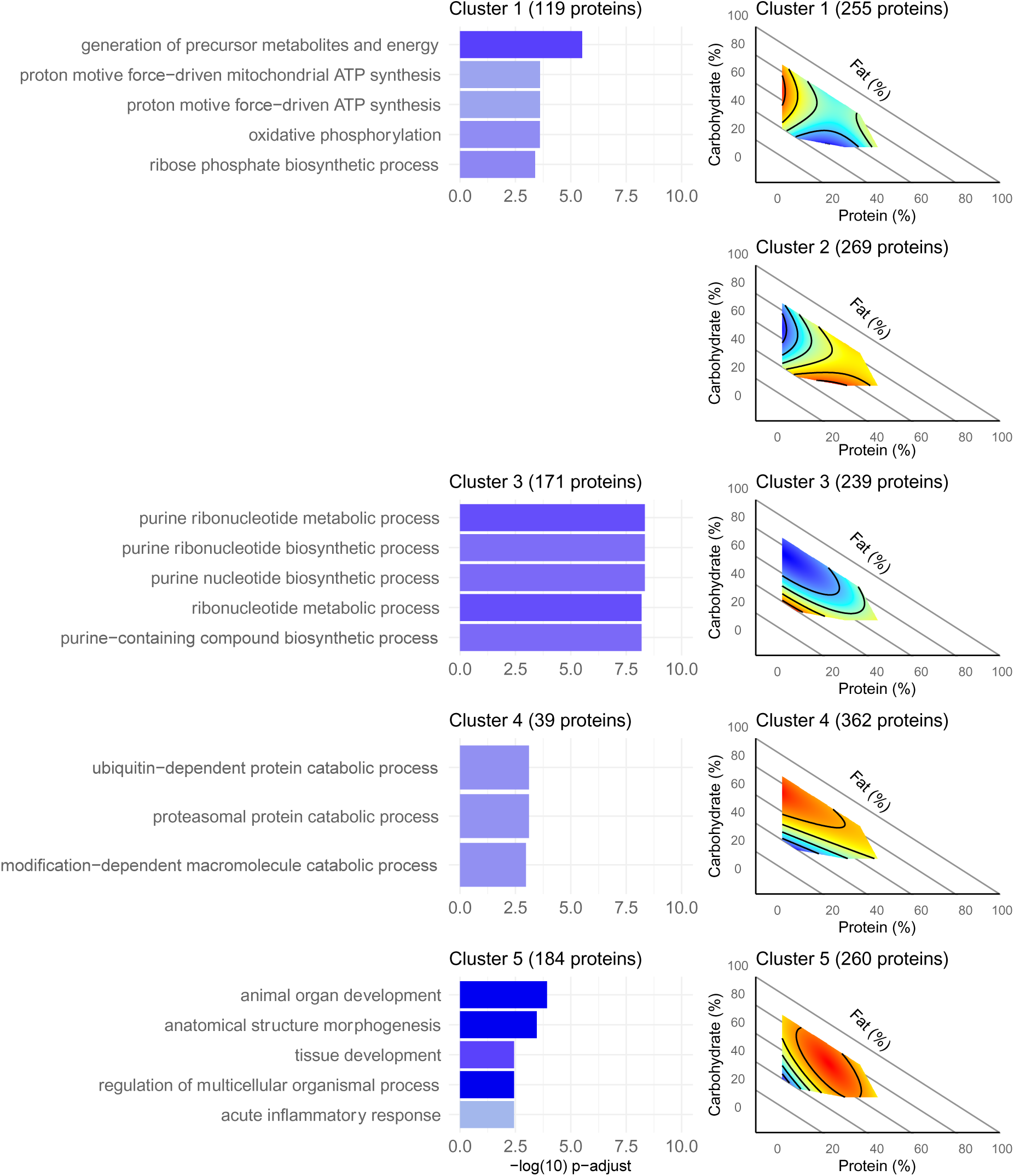
Surface plots of average protein expression across nutritional space, and bar plots of the top five (where possible) enriched biological processes for the five clusters of proteins. For the surface plots, red corresponds to the highest abundance of proteins and blue corresponds to the lowest abundance of proteins. For the bar plots, darker blue corresponds to a higher gene ratio. There were no functionally enriched biological processes for cluster 2.

Clusters 1 and 2 appear inverse to each other (Fig. 2). In cluster 1, proteins had increased abundance on low protein diets and decreased abundance on low carbohydrate diets. In cluster 2, proteins were less abundant on low protein diets and more abundant on low carbohydrate diets (Fig. 2). Functional enrichment analyses of these clusters showed that processes involved with energy production (i.e., ATP synthesis) were significantly enriched in cluster 1 (Fig. 2), and the most connected ‘hub’ proteins in this cluster were cytochrome C1 (CYC1) and ubiquinol-cytochrome C reductase core protein 1 (UQCRC1) (both found in the mitochondria and involved in the electron transport chain) (Fig. 3; Supp. Mat. 1, Fig. S3). We did not detect any significantly enriched pathways in cluster 2, though we note that glucose transporter 4 (GLUT4), which controls insulin-stimulated glucose uptake, belonged to this cluster (Supp. Mat. 1, Fig. S4).

**Fig. 3.**
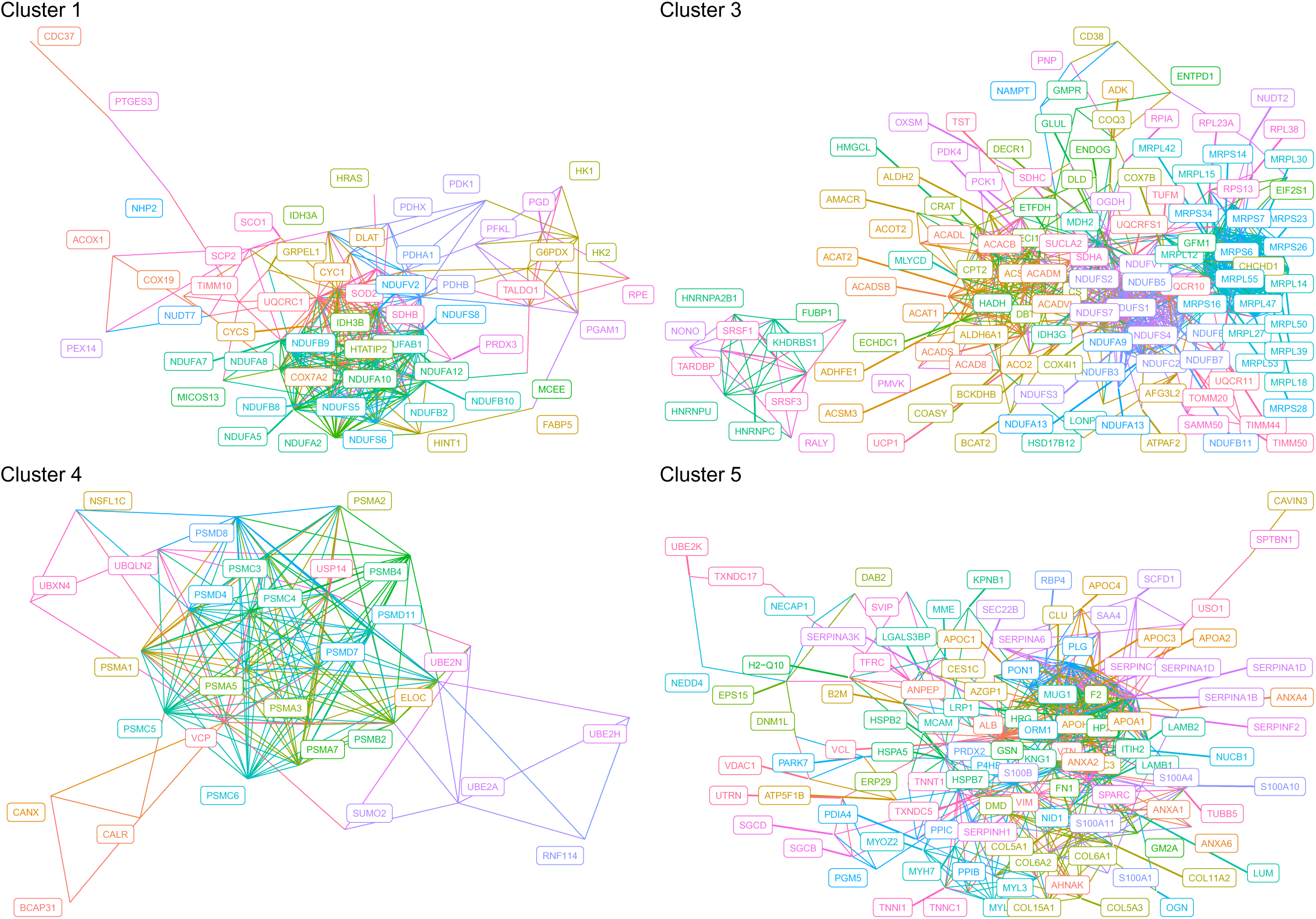
Schematic networks of the significant proteins that were involved in enriched biological processes. Colours represent individual proteins (sorted alphabetically). There were no proteins involved in functionally enriched biological processes for cluster 2.

Clusters 3 and 4 were also inverse to each other and responses were driven by dietary fat content (Fig. 2). Proteins in cluster 3 were mostly in low abundance across the nutritional space but were highly abundant on high-fat diets (i.e., low carbohydrate and low protein diets). In contrast, proteins in cluster 4 were mostly in high abundance across nutritional space but were lowly abundant on high fat diets (Fig. 2). The most significantly enriched pathways in cluster 3 were involved in purine biosynthesis and metabolism (Fig. 2), and the most highly connected hub proteins in this cluster were citrate synthase (CS) and NADH oxidoreductase core subunit S2 (NDUFS2) (Fig. 3; Supp. Mat. 1, Fig S5); involved in the citric acid cycle and the electron transport chain, respectively (note that these two proteins were no longer in our significant ‘set’ of proteins after accounting for dietary intake; Supp. Mat 2). UCP1 also belonged to cluster 3 (Fig. 4) but was not involved in any of our top five enriched biological processes. The most significantly enriched pathways in cluster 4 were related to catabolism of ubiquitinated proteins (Fig. 2), and the most connected hub proteins in this cluster were proteasome proteins PSMB2, PSMC5, PSMD4 (all subunits of the proteasome), and Valosin-containing protein (VCP) (Fig. 3; Fig. 5); all of which are involved in the identification and breakdown on ubiquitinated proteins. Hormone-sensitive lipase (HSL), which facilitates the breakdown of triglycerides into free fatty acids also belonged to cluster 4 but was not significantly involved in any enriched pathways (Fig. 5).

**Fig. 4.**
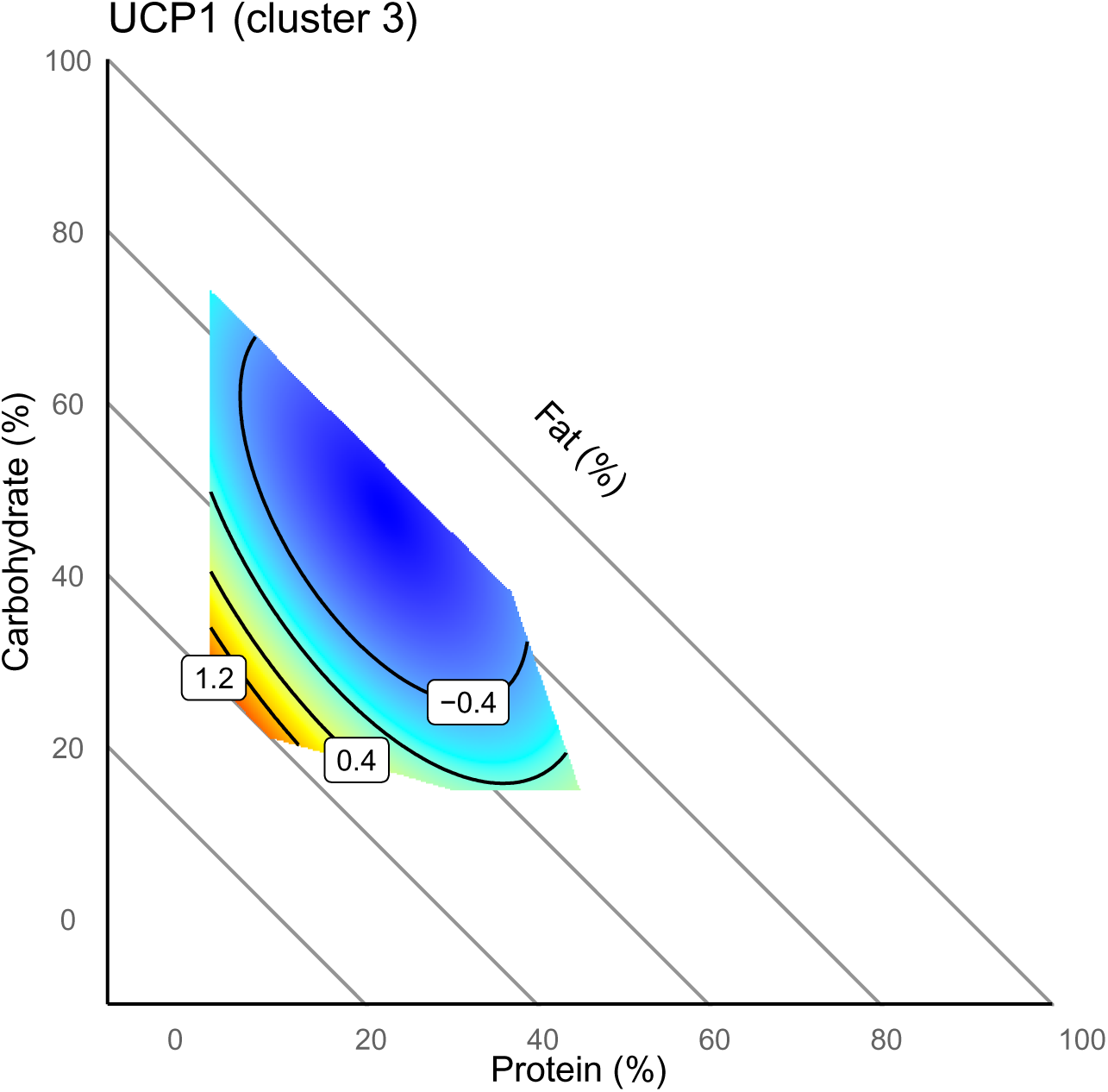
Surface plot of UCP1 (uncoupling protein 1) from cluster 3. Red corresponds to a high abundance of proteins and blue corresponds to a low abundance of proteins.

**Fig. 5.**
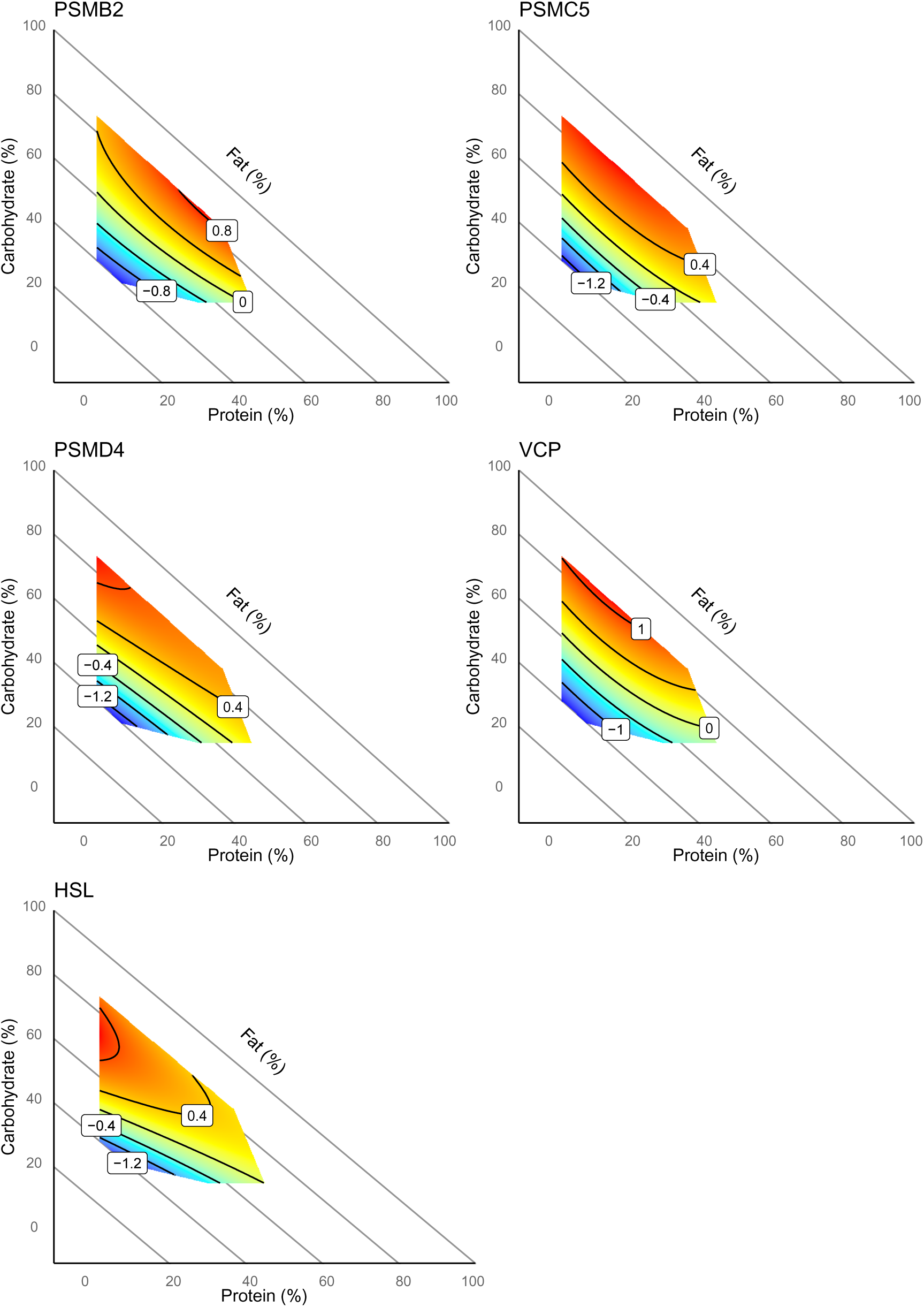
Surface plot of ‘hub’ proteins involved in protein turnover (PSMB2, PSCM5, PSMD4, VCP), as well as HSL, all from cluster 4. Red corresponds to a high abundance of proteins and blue corresponds to a low abundance of proteins. All coefficients are z- transformed.

Of note is that the response surface of cluster 4 closely resembles that of the F0 male BAT weight, while that for cluster 3 is the inverse, suggesting the effect of dietary macronutrients on BAT mass is correlated with the effects of diet on these elements of the BAT proteome. Indeed, when calculating spearman correlations between the individual proteins in cluster 4 mentioned above and BAT weight, we detected moderate positive correlations: Psmb2 *r_s_* = 0.41, Psmc5 *r_s_* = 0.44, Psmd4 *r_s_* = 0.69, Vcp *r_s_* = 0.68, and Hsl *r_s_* = 0.62. Cluster 3, which resembles the inverse of BAT weight also showed moderate negative correlations between the proteins and BAT weight: Ucp1 *r_s_* = -0.51, Cs *r_s_* = -0.61, Ndufs2 *r_s_* = -0.63.

Lastly, cluster 5 was dominated by proteins that display decreased abundance on high fat, low carbohydrate, low protein diets (Fig. 2). The most significantly enriched pathways in this cluster were to do with morphogenesis and development (Fig. 2). The most connected ‘hub’ proteins in this cluster were fibronectin 1 (FN1) and apolipoprotein AI (APOAI) (Fig. 3; Supp. Mat 1, Fig. S6), which are involved in structural organisation and integrity, and supporting lipid metabolism and cholesterol regulation, respectively.

Our significant ‘set’ of proteins was reduced by 61 when dietary intake was included in the models, indicating that variations in these proteins were likely due to differences in food quantity consumed across diets, rather than differences driven by macronutrient composition itself. Our clusters and functional enrichment analyses were qualitatively comparable (see Supp. Mat. 2). However, Cluster 2 had significantly enriched biological processes when accounting for dietary intake (see Supp. Mat. 2).

### Paternal dietary effects on offspring

To gain insight into the effects of pre-conceptional diet with different macronutrient composition on developmental programming of BAT, we next explored paternal effects on the male and female F1 offspring BAT proteome. In female offspring, we found a single protein, basigin (BSG), that was negatively related to the proportion of protein in the paternal diet, after accounting for dietary intake, when only including dietary protein as a main effect (Fig. 6) (F = 13.009, adjusted p-value = 0.027). Indeed, the surface plot of BSG in female offspring closely resembles that of their fathers, but slightly attenuated (Fig. 6). The Spearman correlation between paternal BSG expression and female offspring BSG expression = 0.27.

**Fig. 6.**
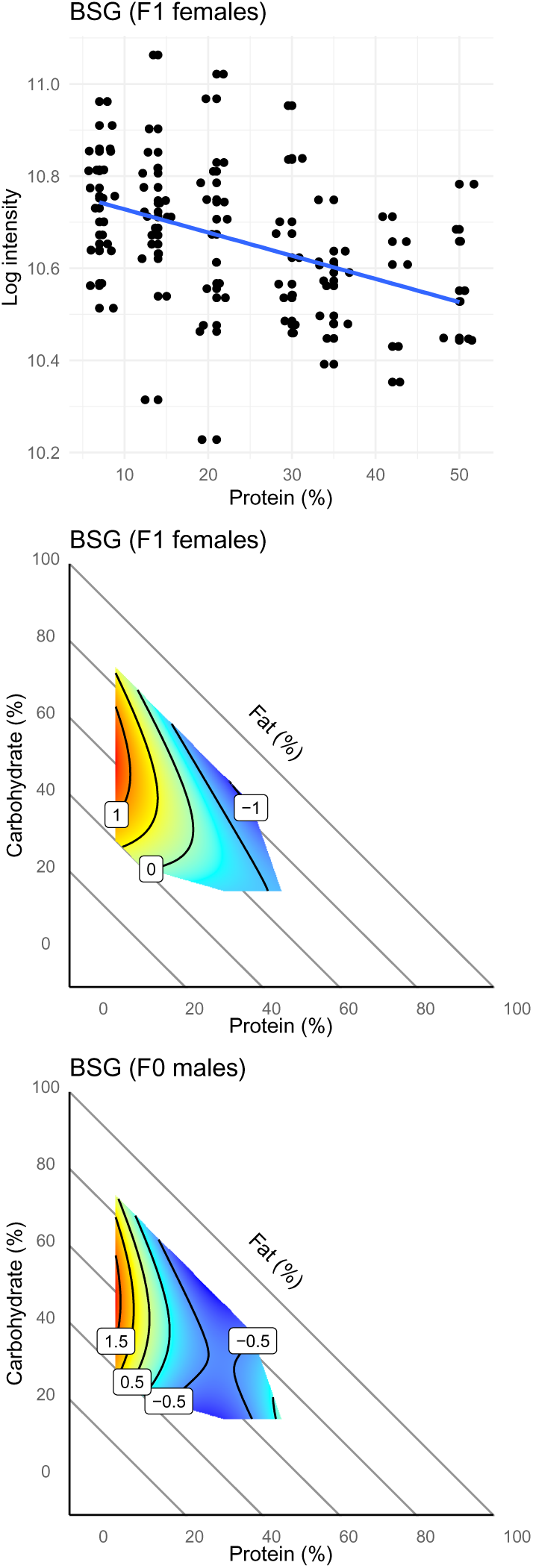
The relationship between paternal diet and basigin (BSG) abundance in F1 females. The scatter plot shows the log intensity of BSG abundance on the y-axis and percent protein in the paternal diet on the x-axis. The surface plots show F1 female and F0 male BSG abundance across paternal nutritional space (coefficients are z-transformed). Red corresponds to a high abundance of proteins and blue corresponds to a low abundance of proteins. See Supp. Mat. 1, Fig. S7 for scatter of log BSG expression and dietary protein in kjs.

We did not detect any effects of paternal diet on BAT protein expression in male offspring. This effect remained consistent in all models (see Supp. Mat. 2). When only examining BSG abundance, we did not find any differences in BSG abundance between the sexes regardless of paternal diet (t = 0.28, p-value = 0.78).

## Discussion

This study is the first to examine how macronutrient balance of dietary protein, carbohydrate, and fat, when accounting for calories, affect the total proteome of brown adipose tissue (BAT) in F0 male C57BL/6J mice and their F1 offspring. We found that in F0 males, over 50% of detected proteins were significantly influenced by diet after correcting for multiple comparisons. Clustering these proteins based on their response to diet and performing functional enrichment analyses revealed patterns with implications for BAT function and non-shivering thermogenesis. Notably, we detected two inversely responding clusters that were correlated with BAT size whereby low-fat diets that resulted in larger interscapular BAT also resulted in lower abundance of UCP1 and proteins involved in purine biosynthesis, but higher abundance of proteins involved in protein turnover. Despite the substantial protein response to diet in the BAT of fathers, we did not detect any effect of paternal macronutrient balance on the BAT proteome in sons, and only detected one protein in daughters, basigin (BSG), which was significantly affected by paternal dietary protein intake. Interestingly, BSG has recently been described to play a role in normal BAT functioning, including UCP1 transcription ^37^.

In fathers, clusters 3 and 4 showed inverse responses to macronutrient balance and were correlated with BAT size. UCP1, part of cluster 3, was negatively correlated with BAT size and was generally under expressed, except on a high-fat diet (low in carbohydrates and protein). While previous studies have shown that dietary fat can upregulate UCP1, most did not control for caloric intake, making it unclear if fat alone drives this effect ^38–40^. Here, all diet formulations were standardised for calories through adjusting non-digestible cellulose, meaning that effects could be attributed to nutrient composition rather than caloric differences (although low-protein males did have higher food intake, results remained similar when intake was accounted for in models). These findings also align with another study ^24^, reporting a negative relationship between BAT size and UCP1 transcription on low protein-to-carbohydrate ratios (fat was fixed in that study). Body temperature increased on these diets, with larger BAT but lower UCP1 expression ^24^, suggesting that alternative, UCP1-independent thermogenic pathways may be active. Indeed, non-shivering thermogenesis can occur independently of UCP125.

We measured an upregulation of proteins involved in protein turnover in cluster 4, including proteasome subunit proteins (19S and 20S), which were overexpressed on low-fat diets that led to larger BAT depots and lower UCP1. The proteasome, a key protein quality-control mechanism, is essential for managing cellular stress under high thermogenic demand ^41,42^. In addition, proteasome inhibition in BAT reduces non-shivering thermogenesis independently of UCP1^41^. Notably, loss of PSMB4 (belonging to cluster 4) disrupts proteostasis and adipogenesis ^43^. UCP1 is also ubiquitinated and degraded by the proteasome ^44^. While we are not able to determine which proteins were being ubiquitinated and degraded in this study, these findings suggest that low-fat diets may not only promote BAT recruitment but may also enhance protein turnover/quality control, despite lower total levels of UCP1. This relationship could be an artefact of increased adiposity in these males, however, as body temperatures can be increased through diet despite lower UCP1 expression ^24^, ubiquitination may play a role in maintaining cellular health under intense thermogenic conditions.

Although UCP1 abundance is generally low except for on high-fat, low-carbohydrate, low-protein diets, we are unable to distinguish between its active and inactive states.

UCP1 activity is regulated by activators such as free fatty acids and is inhibited by purines ^45–47^. Interestingly, together with UCP1, abundance of proteins involved in purine metabolism and biosynthesis were elevated under high-fat diets, while low-fat diets were marked by increased levels of HSL, an enzyme responsible for breaking down triglycerides into free fatty acids ^48^. These findings suggest that macronutrient balance may not only affect UCP1 protein levels but may also play a role in modulating its activation and inhibition. Distinguishing how macronutrient intake mediates active and inactive forms of UCP1 would be a promising future direction to further understand dietary mediated thermogenesis.

There were no detectable effects of paternal diet on the BAT proteome of sons, and only one detectable protein, basigin (BSG), that was negatively affected by paternal dietary protein in daughters. The lack of an intergenerational effect in male offspring may appear surprising given the ever-growing literature showing effects mediated by paternal environment, including dietary effects on offspring metabolic phenotypes^49,50^. However, we feel comfortable concluding that in this experiment paternal diet did not induce a change in BAT proteome amongst male offspring for three reasons. First, we used a dietary design that comprehensively sampled the dietary macronutrient space, and which induced and detected very substantial direct effects of macronutrients on paternal BAT. Second, while it is impossible to statistically prove a null effect, the distribution of p-values for effects in male offspring follows the uniform distribution expected under a null hypothesis as per statistical theory (Supp. Mat. 2). Finally, our results align with previous work on metabolic phenotypes in this study system^9^.

Notably, Crean et al., (2024) ^9^ did not find any effects of paternal dietary macronutrients on the metabolic phenotypes of sons but reported multiple effects of paternal diet on metabolism in daughters. In particular, fathers on diets that promoted increased adiposity produced leaner daughters with enlarged BAT. Albeit that it was the only protein affected, the negative effect of paternal dietary protein on BSG expression in daughters indicates that paternal macronutrient balance can mediate the function of BAT and potentially metabolic health of daughters. Indeed, Rupar et al., (2023)^37^ recently demonstrated that BSG is necessary for normal UCP1 transcription, and other aspects of normal BAT function, but these effects were only evident under stimulation with norepinephrine. Given that our offspring were reared under standard laboratory conditions and were chow-fed, effects of paternal diet, including any effects due to altered BSG expression, may become more pronounced under increased demand for non-shivering thermogenesis (i.e., through diet or colder temperatures).

As mentioned above, our diets were standardised for calories during manufacture. To achieve this, non-digestible cellulose was used, especially to dilute the energy content of the high-fat diets. Consequently, we cannot eliminate the possibility that some effects may be influenced by cellulose. However, given that cellulose has negligible nutritional value, any dietary effects attributable to cellulose are unlikely to be biologically meaningful. Because of this, we believe that the advantages of using isocaloric diets through cellulose manipulation outweigh any potential confounding effects of non-digestible cellulose. Additionally, while the diets were designed to be standardised for calories, mice had *ad libitum* access to their assigned diets. We also observed differences in food intake based on dietary protein. However, our results remained qualitatively comparable when dietary intake was included as a co-variate in our models. Lastly, we used fuzzy-c clustering to identify patterns in protein response surfaces and conduct functional enrichment analyses. This clustering method assigns proteins to multiple clusters based on a membership score, which allows for overlapping responses. However, clustering analyses are sensitive to the number of clusters selected, and membership ambiguity can arise when protein responses overlap across multiple clusters. As a result, protein assignment to a cluster may vary depending on the chosen number of clusters and membership thresholds. To mitigate this, we determined the optimal number of clusters using a combination of clustering indices and visual inspection of individual protein response surfaces within clusters.

While protein assignments may vary slightly between clustering approaches, pilot analyses using different clustering divisions showed that our results remained robust.

Overall, our results add nuance to our understanding of adaptive dietary effects on BAT size and function by showing that low-fat diets (especially when paired with high carbohydrate and low protein) results in larger BAT and lower UCP1 protein, but higher total adiposity and poorer metabolic outcomes compared to high-fat diets if caloric intake remains the same. Nevertheless, while total UCP1 may be lower, larger BAT and an over expression of other biological processes known to be important for non-shivering thermogenesis (i.e., protein turnover and potentially differential activation and inhibition of UCP1 through free fatty acids and purines), may be acting to ameliorate any further detrimental effects of chronic low-fat exposure. It also advances our understanding of how paternal macronutrient balance may affect offspring BAT function, particularly in daughters. New avenues for future research include investigating targets of protein turnover and differences in UCP1 activation and inhibition across nutritional space. Additionally, deeper research into how paternal macronutrient balance may affect the BAT proteome of offspring when under high thermogenic demand would be interesting. Our study has implications not only for elucidating potentially adaptive dietary responses in BAT and thus, metabolism, but also for therapeutics and public health.

## Data Availability

Data Independent Acquisition (DIA) proteomics data will be deposited to the ProteomeXchange Consortium via the PRIDE partner repository. The R code and processed protein intensity data used for data analysis can be found at https://osf.io/j9dv4/ (will be made public upon acceptance).

## Supporting information

Supp Mat 1

Supp Mat 2

## Acknowledgements

We would like to acknowledge all members of the Gametic Epigenetics Consortium against Obesity (GECKO) for their feedback and input, as well as members of the Simpson research group. We acknowledge the Sydney Mass Spectrometry facility for access and support. We also acknowledge the Charles Perkins Centre at the University of Sydney for the infrastructure and facilities including staff from Laboratory Animal Services for assistance with animal care. This work was supported by a Challenge Programme Grant from the Novo Nordisk Foundation (NNF18OC0033754) to the Gametic Epigenetics Consortium against Obesity. The Novo Nordisk Foundation Center for Basic Metabolic Research is an independent research center at the University of Copenhagen, partially funded by an unrestricted donation from the Novo Nordisk Foundation (NNF18CC0034900). This work was supported by the French Government (National Research Agency, ANR) through the “Investments for the Future” programs LABEX SIGNALIFE ANR-11-LABX-0028-01 and IDEX UCAJedi ANR-15-IDEX-01.

## Author Contributions

S.J.S., R.B., M.A.N., and A.S. have participated in the study design. A.J.C. and T.J.P. conducted the animal work, and E.L.M. and L.S. conducted the proteomics and mass spectrometry work. E.L.M. analysed the data with the help of L.S. and A.S. E.L.M. drafted the manuscript. All authors provided input and approved the final version of the manuscript.

## Competing interests

The authors have no competing interests to declare.

## Notes

https://osf.io/j9dv4/

### Competing Interest Statement

The authors have declared no competing interest.

