## Supplementary material for "Dietary macronutrients modulate the proteome of brown adipose tissue in males and their female offspring": Supp Mat 1

**Supplementary material 1: Figures**

| Diet | % Prot | % Carb | % Fat |
|---|---|---|---|
| 1 | 7 | 33 | 60 |
| 2 | 7 | 78 | 15 |
| 3 | 14 | 56 | 30 |
| 4 | 14 | 26 | 60 |
| 5 | 21 | 64 | 15 |
| 6 | 21 | 34 | 45 |
| 7 | 30 | 40 | 30 |
| 8 | 35 | 20 | 45 |
| 9 | 42 | 43 | 15 |
| 10 | 50 | 20 | 30 |

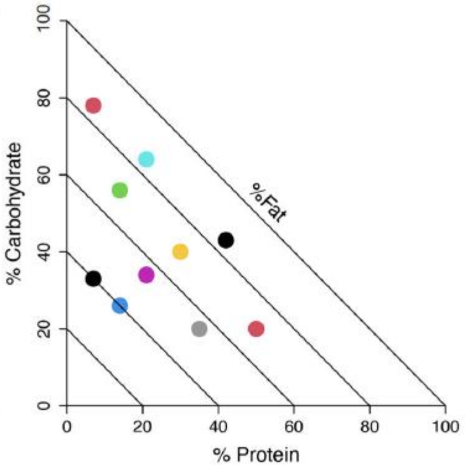


**Fig. S1**. Figure from Crean et al (2024). The macronutrient composition of the 10 isocaloric diets that fathers were reared on. Each diet is plotted on a right-angled mixture triangle where movement along the x-axis represents an increase in protein (%), movement up the y-axis represents an increase in carbohydrate (%) and movement towards the origin from the diagonal edge represents an increase in fat (%).

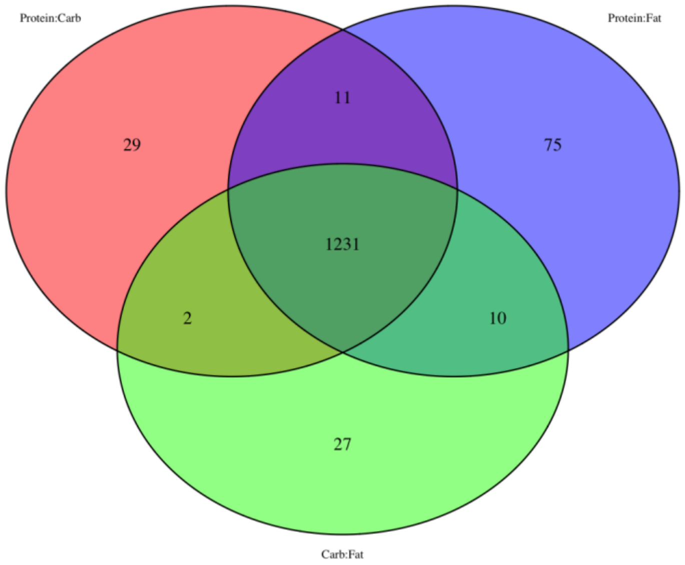


**Fig. S2.** Venn diagram showing the number of proteins that were influenced by the different combinations of nutrients in the diet of F1 males to be used for further analyses. Each circle encompasses linear/additive and quadratic/interactive effects of the nutritional components.

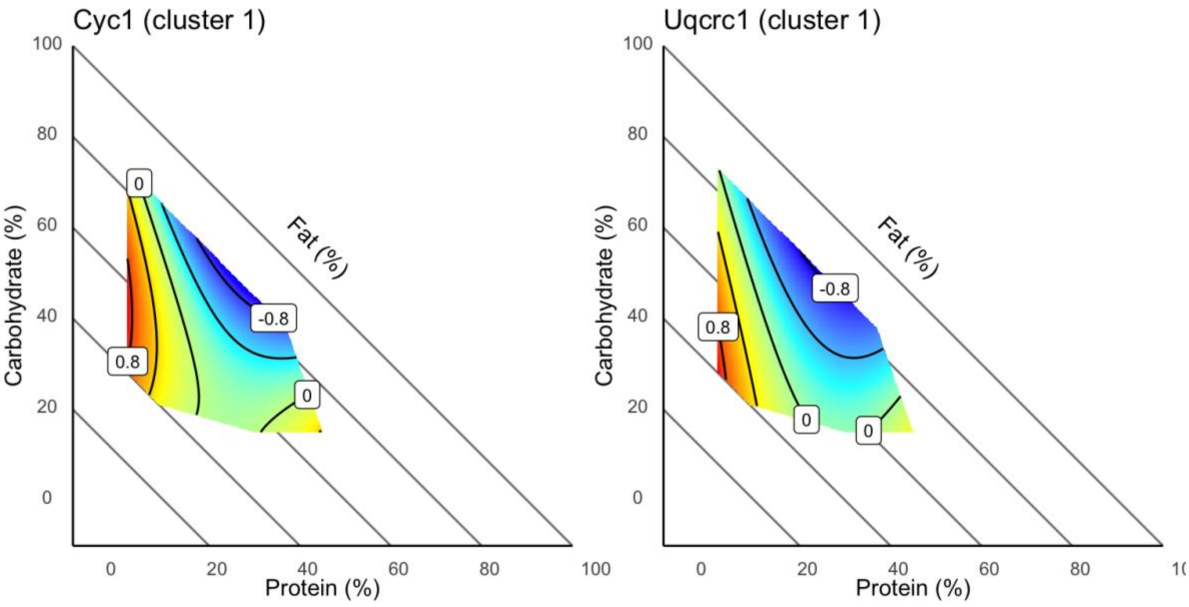


**Fig. S3.** Surface plot for the two 'hub' proteins in cluster 1: CYC1 (cytochrome c) and UQCRC1 (ubiquinol-cytochrome c reductase core protein 1). Red corresponds to a high abundance of proteins and blue corresponds to a low abundance of proteins.

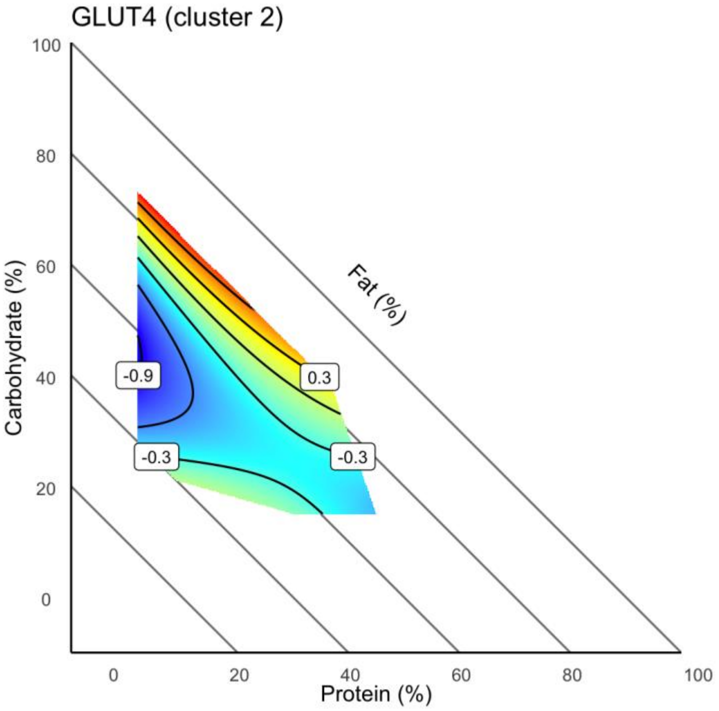


**Fig. S4.** Surface plot for GLUT4 (glucose transporter type 4) in cluster 2. Red corresponds to a high abundance of proteins and blue corresponds to a low abundance of proteins.

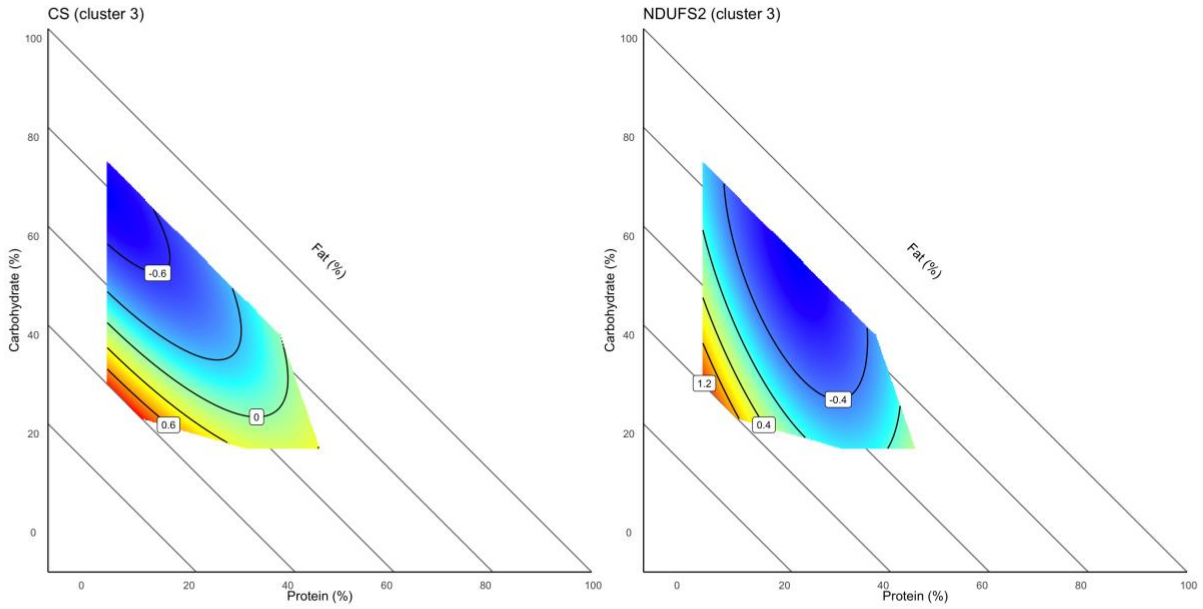


**Fig. S5.** Surface plot for the two ‘hub’ proteins in cluster 3: CS cCitrate synthase) and NDUFS2 (NADH oxidoreductase core subunit S2). Red corresponds to a high abundance of proteins and blue corresponds to a low abundance of proteins.

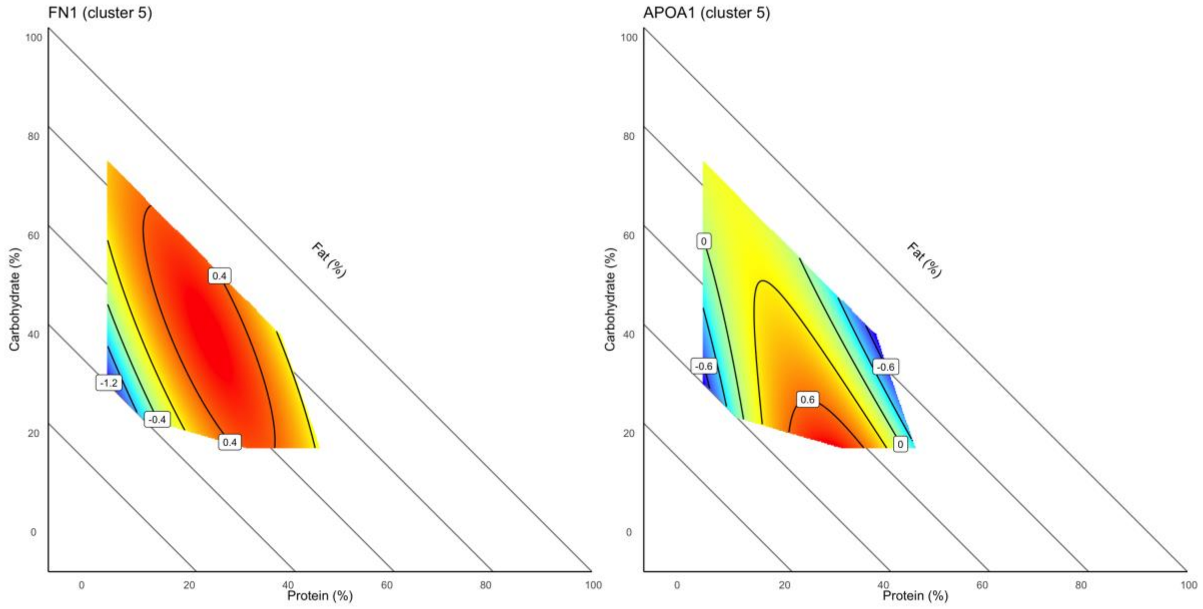


**Fig. S6.** Surface plot for the two ‘hub’ proteins in cluster 5: FN1 (fibronectin 1) and APOA1(apolipoprotein A-I). Red corresponds to a high abundance of proteins and blue corresponds to a low abundance of proteins.

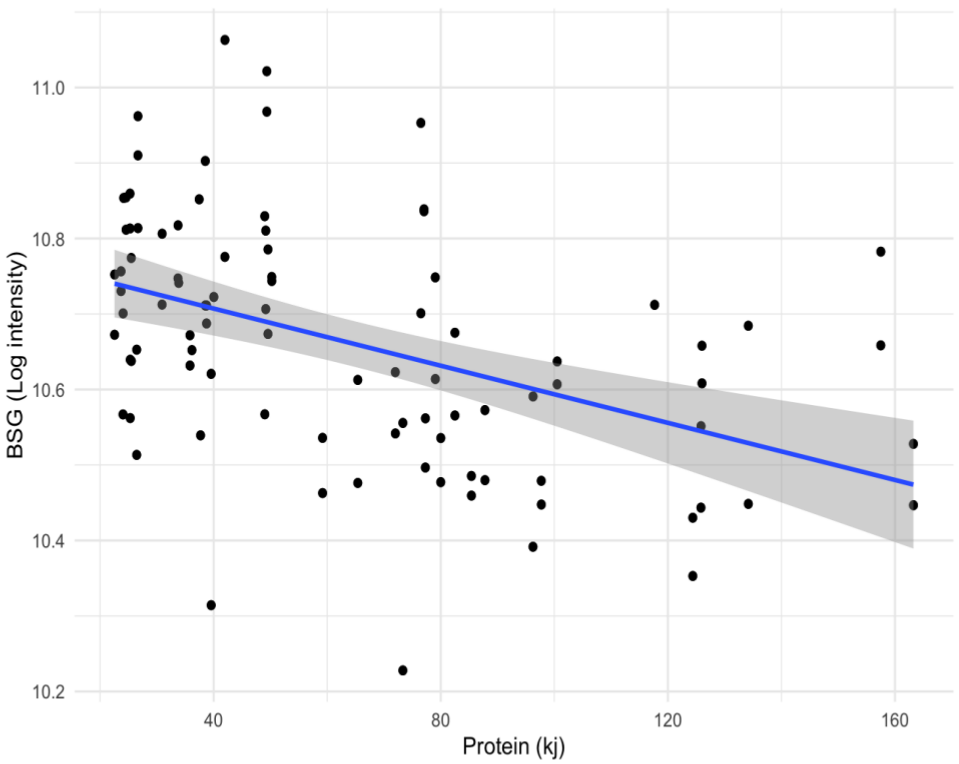


**Fig. S7** Scatter plot of the relationship between paternal dietary protein intake (kjs) and log intensity of BSG expression.
