## Supplementary material for "Dietary macronutrients modulate the proteome of brown adipose tissue in males and their female offspring": Supp Mat 2

### Supplementary Material 2

This file contains all the supplementary analyses:

- Including food intake as a covariate
- All models run on male and female offspring
- Surface plots of 'hub' proteins

### 1 Loading packages

```
# if (!requireNamespace('BiocManager', quietly = TRUE)) {
# install.packages('BiocManager') }
# BiocManager::install('SummarizedExperiment')

# if (!require('BiocManager', quietly = TRUE)) install.packages('BiocManager')
# BiocManager::install('QFeatures')

# if (!requireNamespace('BiocManager', quietly = TRUE))
# install.packages('BiocManager') BiocManager::install('limma')

# if (!require('BiocManager', quietly = TRUE)) install.packages('BiocManager')
# BiocManager::install('preprocessCore')

# if (!requireNamespace('BiocManager', quietly = TRUE))
# install.packages('BiocManager') BiocManager::install('clusterProfiler')

# BiocManager::install('org.Mm.eg.db') # Use the appropriate organism database

library(pacman)

pacman::p_load(tidyverse, here, readxl, readr, Rmisc, devtools, str2str, plotly,
    data.table, statmod, patchwork, pheatmap, Rfast, janitor, SummarizedExperiment,
    QFeatures, limma, mixexp, sp, geometry, gridExtra, mgcv, metR, preprocessCore,
    mgcv, VennDiagram, lme4, tidyheatmaps, NbClust, clusterProfiler, org.Mm.eg.db,
    ggfun, e1071, scatterplot3d, plotly, mixexp, STRINGdb, ggraph, igraph, tidygraph)
```

### 2 Load data

```
raw_dat <- read_tsv("Data/unique_genes_matrix8.tsv")
metadata <- read_csv("Data/Metadata.csv")
intake <- read_csv("Data/Intake.csv")  #includes paternal BAT and growth
offspring_weights <- read_csv("Data/Offspring_weights.csv")
average_females <- read_csv("Data/Average_female_BATweight.csv")
Cluster1_imported <- read.csv("Data/Cluster1.csv")  #used for surfaces
# #Loading Summarised Experiment of log transformed data with MAR and >30%
# missing proteins removed
SummarizedExperiment <- readRDS("Data/SummarizedExperiment.rds")

# identical(SummarizedExperiment@colData$Sample_name,colnames(assay(SummarizedExperiment)))
```

### 3 MDS plot

```
mds <- plotMDS(assay(SummarizedExperiment, 1), plot = FALSE, top = 1000)

mds_df <- data.frame(Sample_name = colnames(mds$distance.matrix.squared), X = mds$x,
    Y = mds$y, Dim1 = mds$eigen.vectors[, 1], Dim2 = mds$eigen.vectors[, 2], Dim3 = mds$eigen.vectors[,
        3], diet = SummarizedExperiment$Diet, sex = SummarizedExperiment$Sex, batch = SummarizedExperim
    generation = SummarizedExperiment$Offspring_vs_dad)

mds_df$sex <- as.factor(mds_df$sex)
mds_df$diet <- as.factor(mds_df$diet)
mds_df$batch <- as.factor(mds_df$batch)
mds_df$generation <- as.factor(mds_df$generation)

mds_offspring <- mds_df %>%
    filter(generation == "O")
mds_dads <- mds_df %>%
    filter(generation == "P")

MDS_plotter <- function(assay_data, colored_by) {
    library(ggplot2)
    library(patchwork)

    # df1 <- na.omit(assay_data)
    df1 <- assay_data
    graph1 <- ggplot(df1, aes_string(x = "Dim1", y = "Dim2", color = colored_by,
        text = "Sample_name")) + geom_point(size = 3)

    graph2 <- ggplot(df1, aes_string(x = "Dim1", y = "Dim3", color = colored_by,
        text = "Sample_name")) + geom_point(size = 3)

    graph3 <- ggplot(df1, aes_string(x = "Dim2", y = "Dim3", color = colored_by,
        text = "Sample_name")) + geom_point(size = 3)

    combined_graph <- graph1/graph2/graph3 + plot_layout(guides = "collect")

    return(combined_graph)
}

generation_MDS <- MDS_plotter(mds_df, "generation")
batch_MDS <- MDS_plotter(mds_df, "batch")

sex_MDS <- MDS_plotter(mds_offspring, "sex")
diet_MDS_dads <- MDS_plotter(mds_dads, "diet")
diet_MDS_offspring <- MDS_plotter(mds_offspring, "diet")
```

##### 3.1 Generation

```r
print(generation_MDS)
```

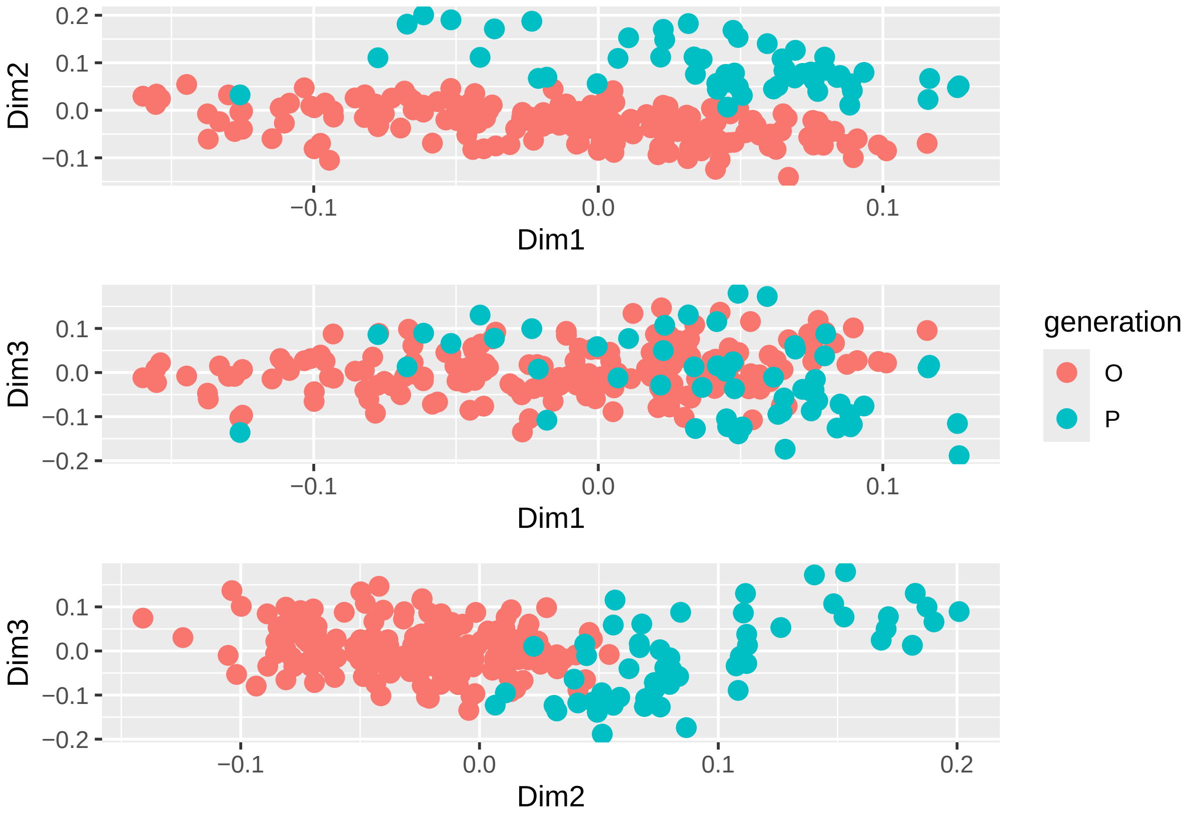

##### 3.2 Offspring sex

```r
print(sex_MDS)
```

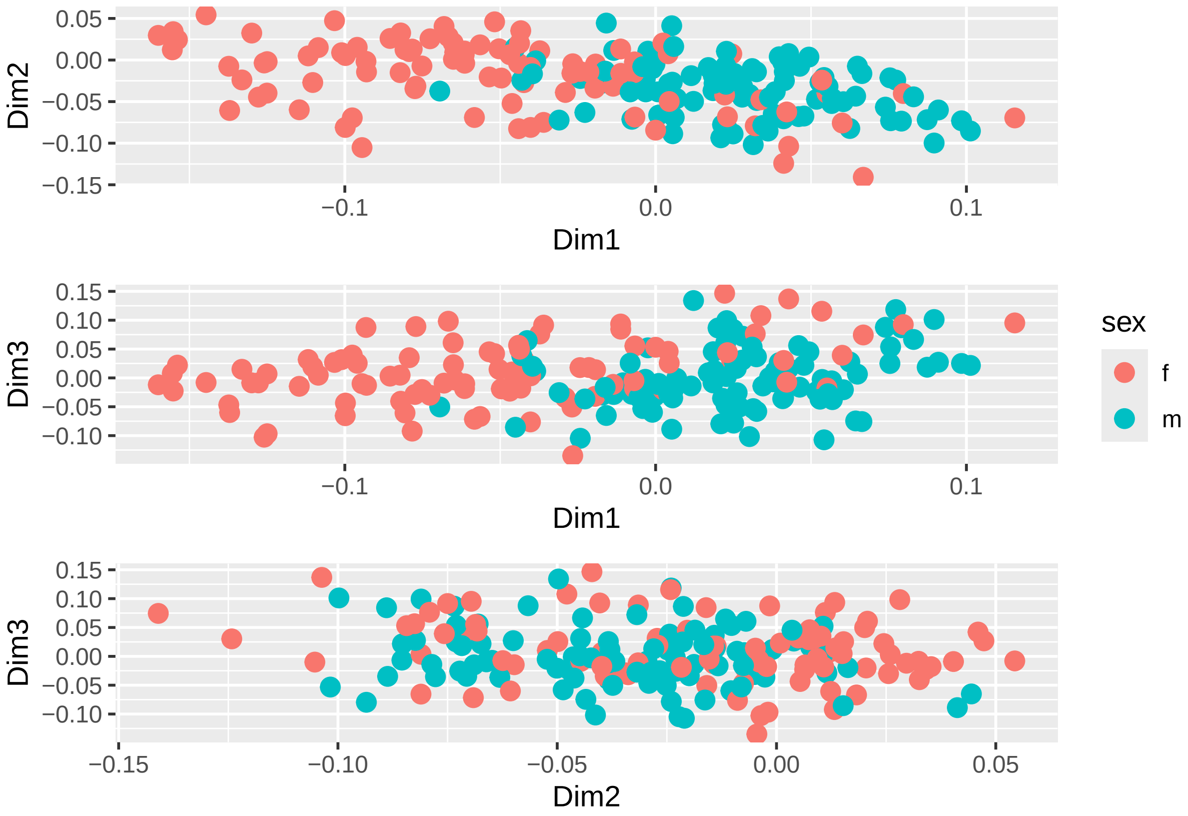

##### 3.3 Diet on F0 males

```
print(diet_MDS_dads)
```

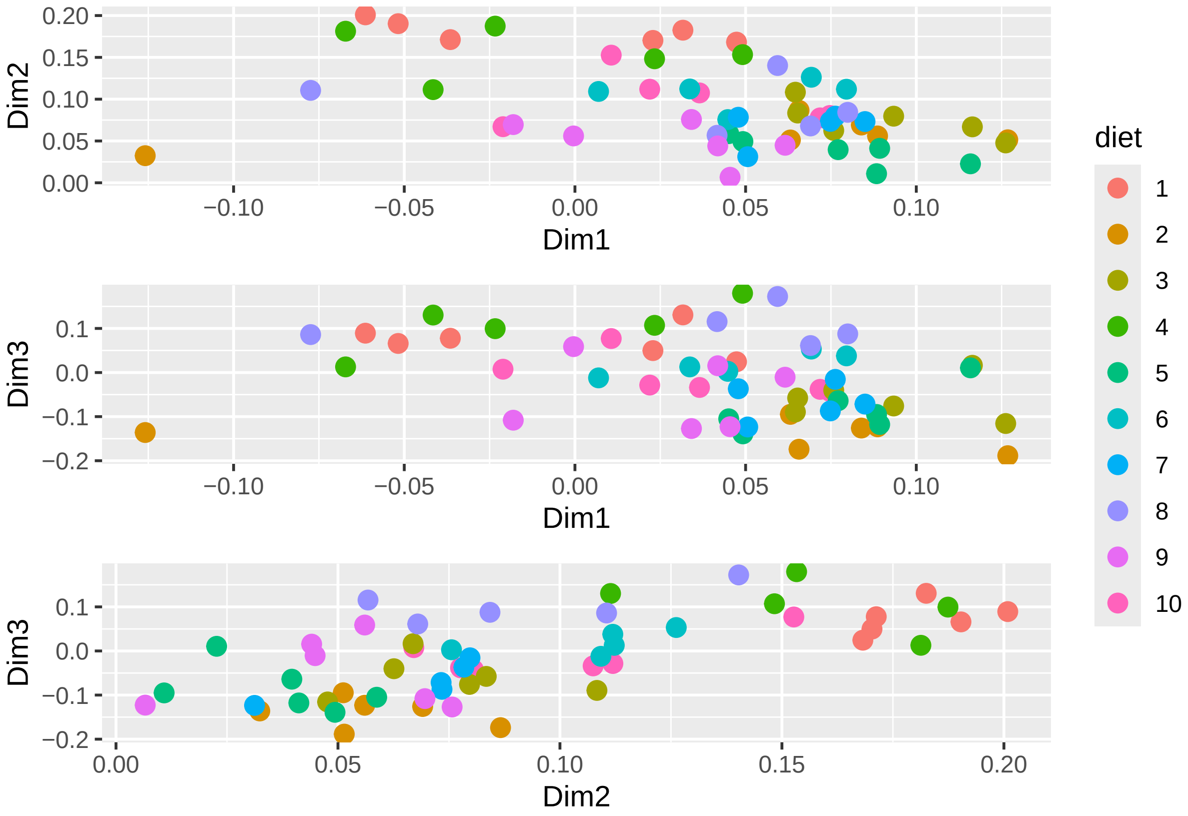

##### 3.4 Diet on F1 ofspring

```
print(diet_MDS_offspring)
```

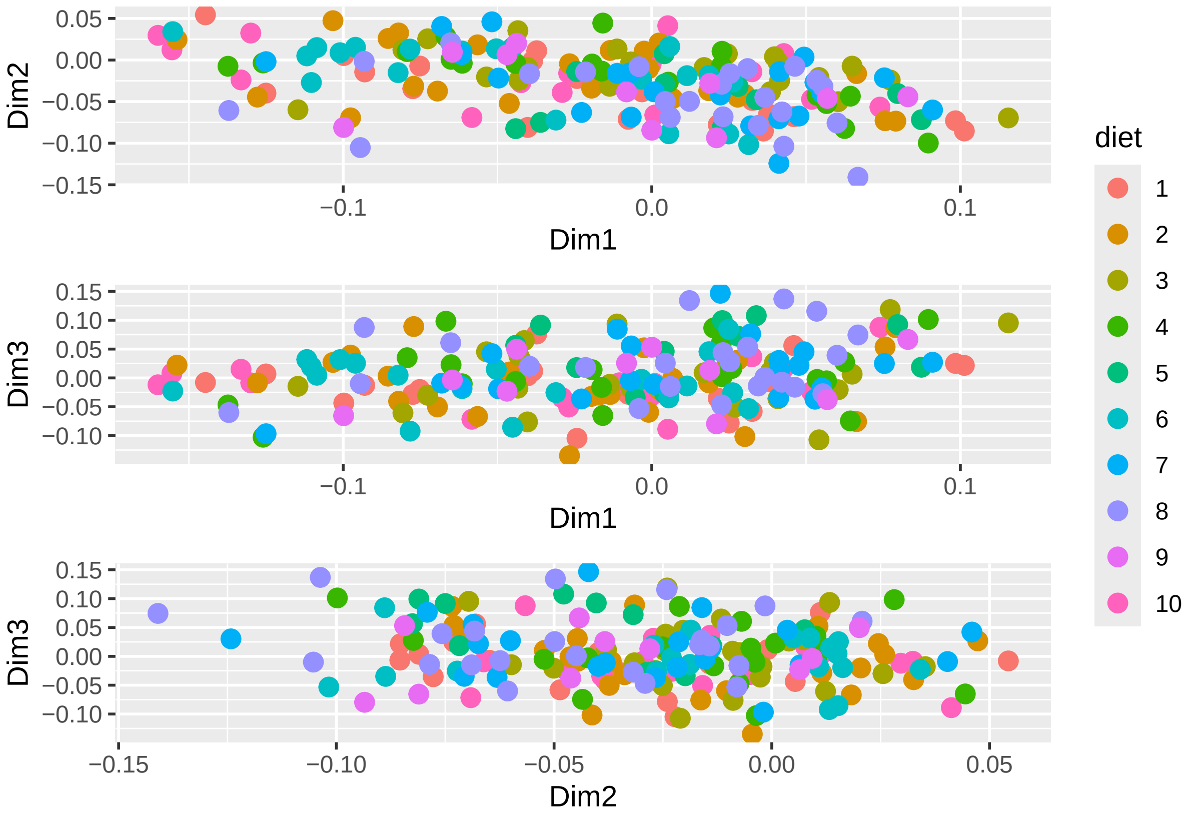

##### 3.5 Diet and sex, F1 offspring

```
mds1v2 <- ggplot(mds_offspring, aes(Dim1, Dim2)) + geom_text(aes(label = sex, color = diet),
    size = 3)
mds1v3 <- ggplot(mds_offspring, aes(Dim1, Dim3)) + geom_text(aes(label = sex, color = diet),
    size = 3)
mds2v3 <- ggplot(mds_offspring, aes(Dim2, Dim3)) + geom_text(aes(label = sex, color = diet),
    size = 3)

mds1v2/mds1v3/mds2v3
```

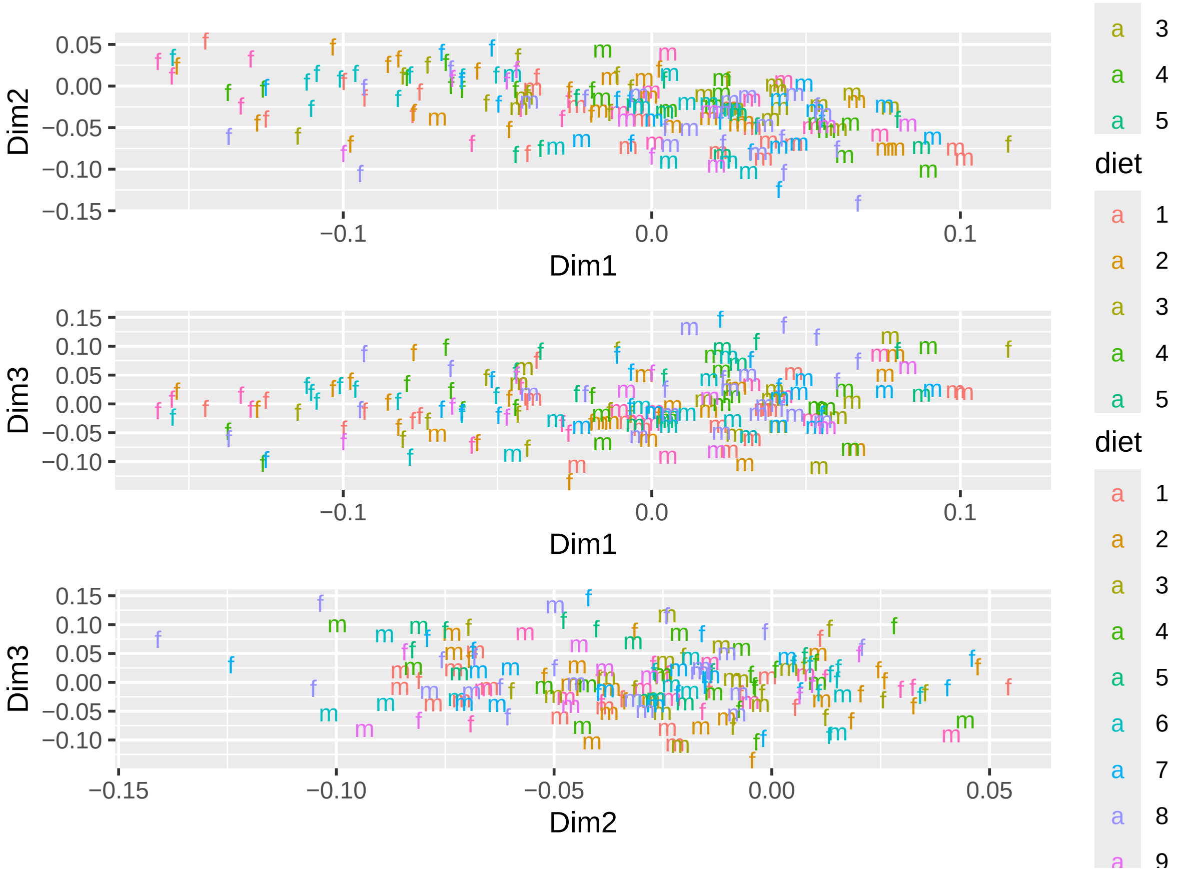

##### 3.6 Batch

```
print(batch_MDS)  # No batch/plate separation so don't need to do a batch correction
```

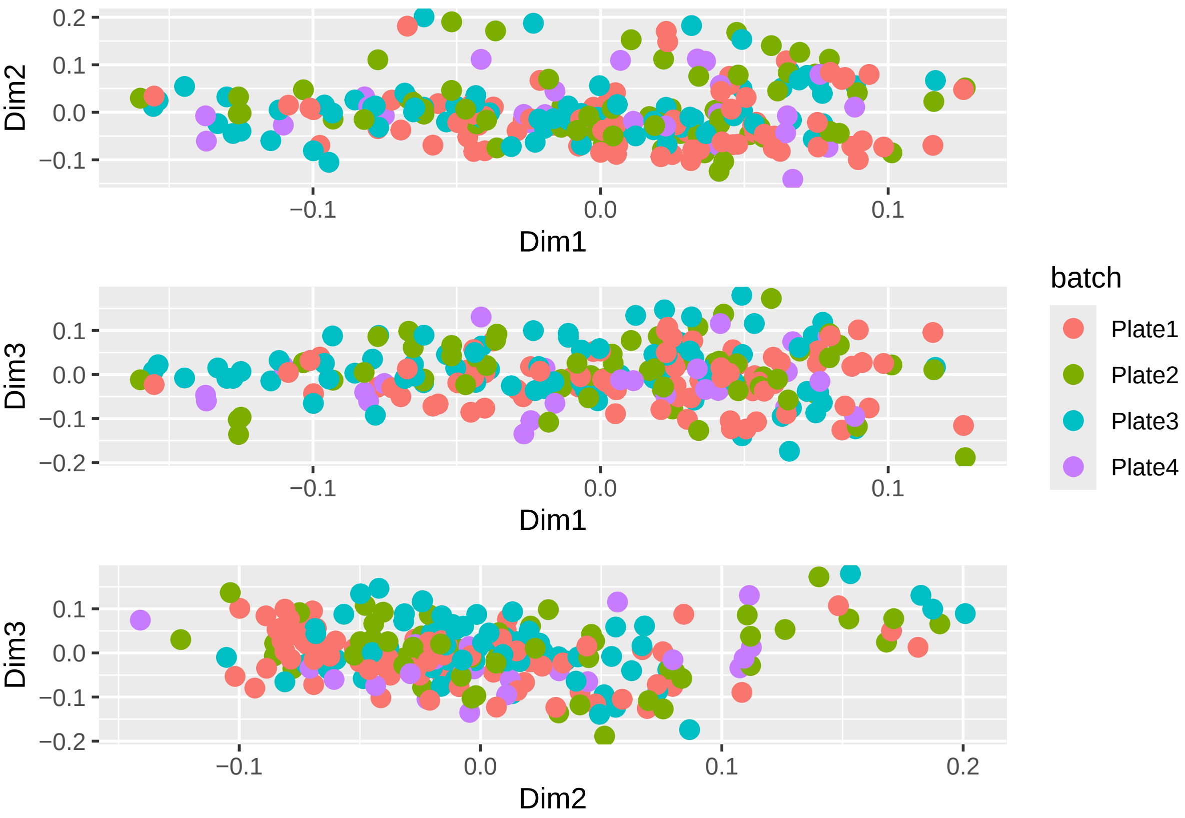

### 4 F0 Males (fathers) analysis with dietary intake as co-var

This analysis includes dietary intake as a co-variate for all our additive/linear and interaction/quadratic models to see if our 'set' of significant proteins change.

Numbers of significant proteins only differed slightly

```
# separating sumamrised experiment object

SummarizedExperiment_offspring <- SummarizedExperiment[, SummarizedExperiment$Offspring_vs_dad ==
    "O"]

# identical(SummarizedExperiment_offspring@colData$Sample_name,colnames(assay(SummarizedExperiment_offs

SummarizedExperiment_fathers <- SummarizedExperiment[, SummarizedExperiment$Offspring_vs_dad ==
    "P"]

# identical(SummarizedExperiment_fathers@colData$Sample_name,colnames(assay(SummarizedExperiment_father

SummarizedExperiment_females <- SummarizedExperiment_offspring[, SummarizedExperiment_offspring$Sex ==
    "f"]

# identical(SummarizedExperiment_females@colData$Sample_name,colnames(assay(SummarizedExperiment_female

SummarizedExperiment_males <- SummarizedExperiment_offspring[, SummarizedExperiment_offspring$Sex ==
```

```r
    "m"]

# identical(SummarizedExperiment_males@colData$Sample_name,colnames(assay(SummarizedExperiment_males)))
```

##### 4.0.1 Protein and carbs (%)

```r
# create design matrix
Percent_prot = as.numeric(SummarizedExperiment_fathers$Percent_prot)
Percent_carb = as.numeric(SummarizedExperiment_fathers$Percent_carb)
dietary_intake = as.numeric(SummarizedExperiment_fathers$Food_intake)

design1 <- model.matrix(~Percent_prot + Percent_carb + dietary_intake)

data_sample_names <- colnames(SummarizedExperiment_fathers)
design_sample_names <- rownames(design1)

rownames(design1) <- data_sample_names
```

```r
fit1 <- lmFit(assay(SummarizedExperiment_fathers), design1)
fit1 <- eBayes(fit1)
table1 <- topTable(fit1, adjust = "fdr", sort.by = "F")
```

###### 4.0.1.1 Additive model

```r
table1b <- topTable(fit1, sort = "none", number = Inf, adjust = "fdr")

count(table1b$adj.P.Val < 0.05)
```

###### 4.0.1.1.1 Number of significant proteins

```
## [1] 1198
```

```r
hist(table1b$P.Value)
```

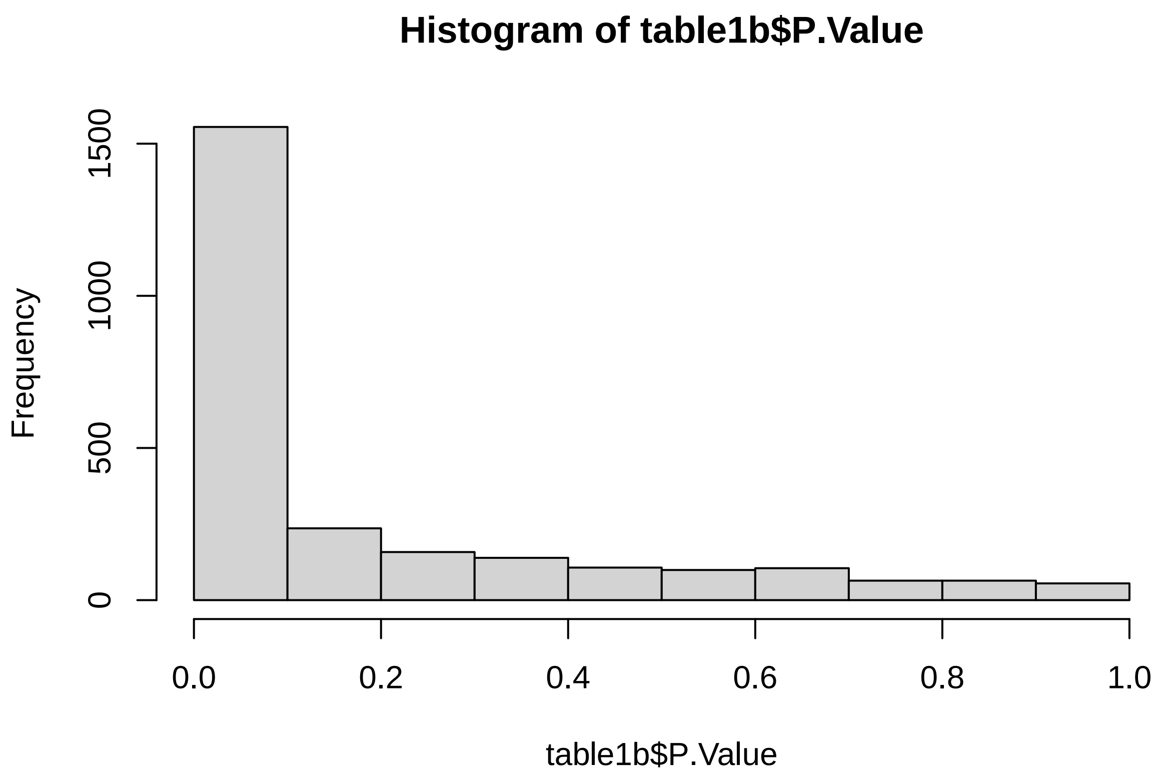

```r
# create design matrix
design2 <- model.matrix(~Percent_prot * Percent_carb + dietary_intake)

data_sample_names <- colnames(SummarizedExperiment_fathers)
design_sample_names <- rownames(design2)

rownames(design2) <- data_sample_names
```

###### 4.0.1.2 Interaction/quadratic model

```r
fit2 <- lmFit(assay(SummarizedExperiment_fathers), design2)
fit2 <- eBayes(fit2)
```

```r
table2_interaction <- topTable(fit2, sort = "none", number = Inf, adjust = "fdr")
count(table2_interaction$adj.P.Val < 0.05)
```

###### 4.0.1.2.1 Number of significant proteins

```
## [1] 1124
```

```r
hist(table2_interaction$P.Value)
```

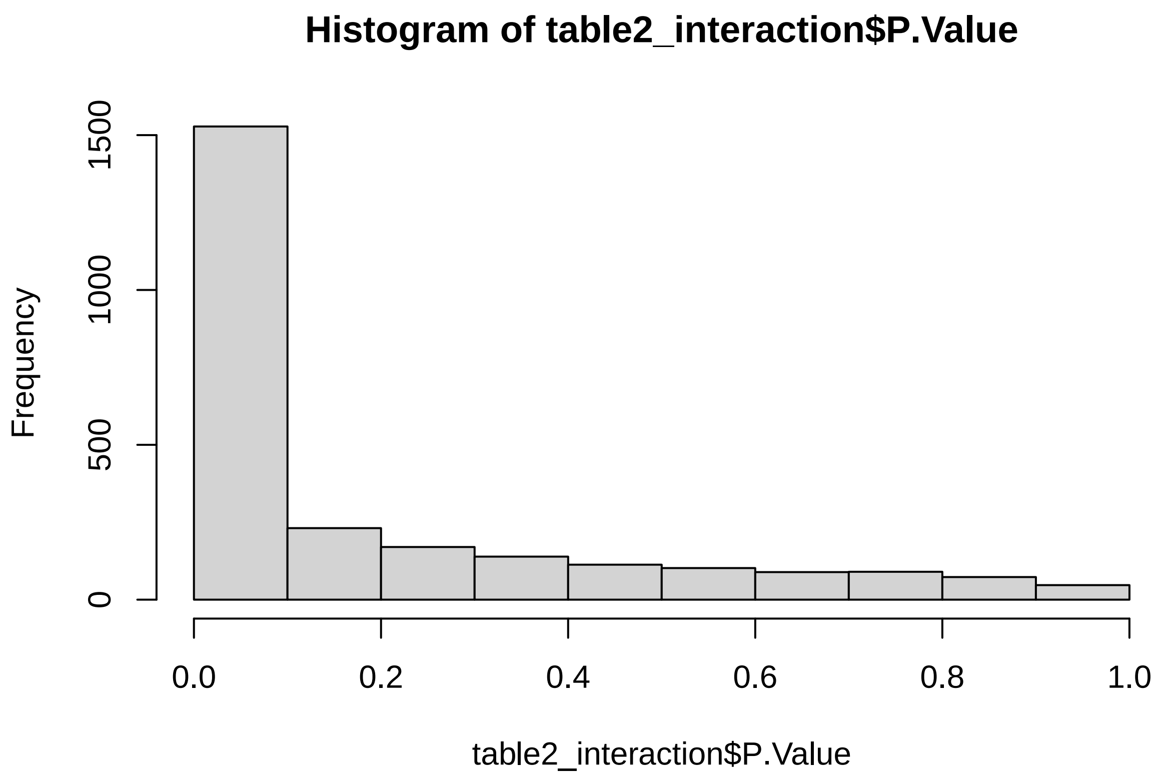

**4.0.1.3 Comparing models** 1237 proteins were significantly affected by diet in the main text 1234 proteins were significantly affected by diet when dietary intake was included

```r
# This is only taking the rownames
Sig_additive1 <- rownames(table1b[table1b$adj.P.Val < 0.05, ])
Sig_interaction1 <- rownames(table2_interaction[table2_interaction$adj.P.Val < 0.05,
    ])

# removing interaction column so that both tables are comparable
# Sig_interaction <- subset(Sig_interaction, select =
# -Percent_prot.Percent_carb)

# find common and different protein

common_proteins1 <- intersect(Sig_additive1, Sig_interaction1)
unique_to_additive1 <- setdiff(Sig_additive1, Sig_interaction1)
unique_to_interaction1 <- setdiff(Sig_interaction1, Sig_additive1)

common1 <- length(common_proteins1)
additive1 <- length(unique_to_additive1)
interaction1 <- length(unique_to_interaction1)
```

```r
v1 <- draw.pairwise.venn(area1 = additive1 + common1, area2 = interaction1 + common1,
    cross.area = common1, category = c("Additive Model", "Interaction Model"), fill = c("blue",
        "red"), alpha = 0.5, cex = 1.5, cat.cex = 1.5, cat.pos = c(-20, 20), cat.dist = 0.05,
    0.05, ind = FALSE)

grid.newpage()
grid.draw(v1)
```

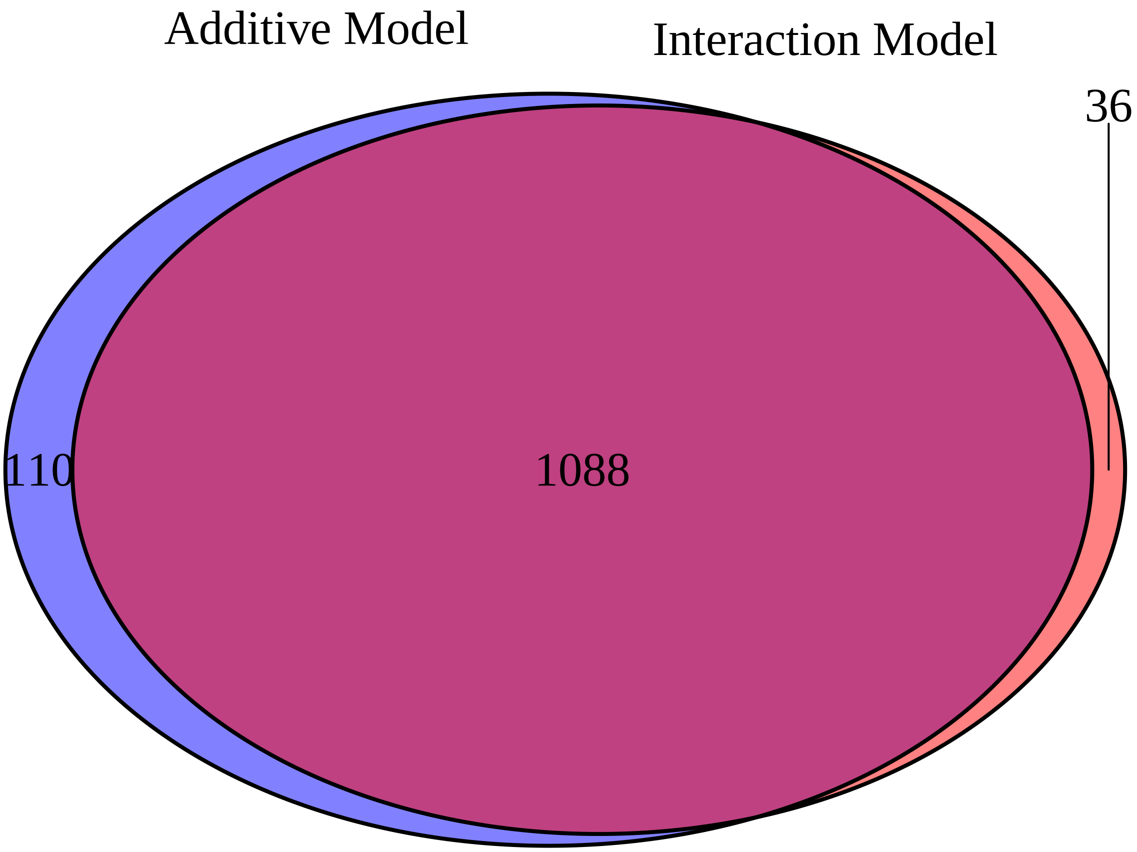

###### 4.0.2 Protein and fat (%)

```r
# create design matrix
Percent_prot = as.numeric(SummarizedExperiment_fathers$Percent_prot)
Percent_fat = as.numeric(SummarizedExperiment_fathers$Percent_fat)
dietary_intake = as.numeric(SummarizedExperiment_fathers$Food_intake)

design3 <- model.matrix(~Percent_prot + Percent_fat + dietary_intake)

data_sample_names <- colnames(SummarizedExperiment_fathers)
design_sample_names <- rownames(design3)

rownames(design3) <- data_sample_names
```

```r
fit3 <- lmFit(assay(SummarizedExperiment_fathers), design3)
fit3 <- eBayes(fit3)
table3 <- topTable(fit3, adjust = "fdr", sort.by = "F")
# table6
```

###### 4.0.2.1 Additive model

```r
table3b <- topTable(fit3, sort = "none", number = Inf, adjust = "fdr")

count(table3b$adj.P.Val < 0.05)
```

###### 4.0.2.1.1 Number of significant proteins

```
## [1] 1202
```

```r
hist(table3b$P.Value)
```

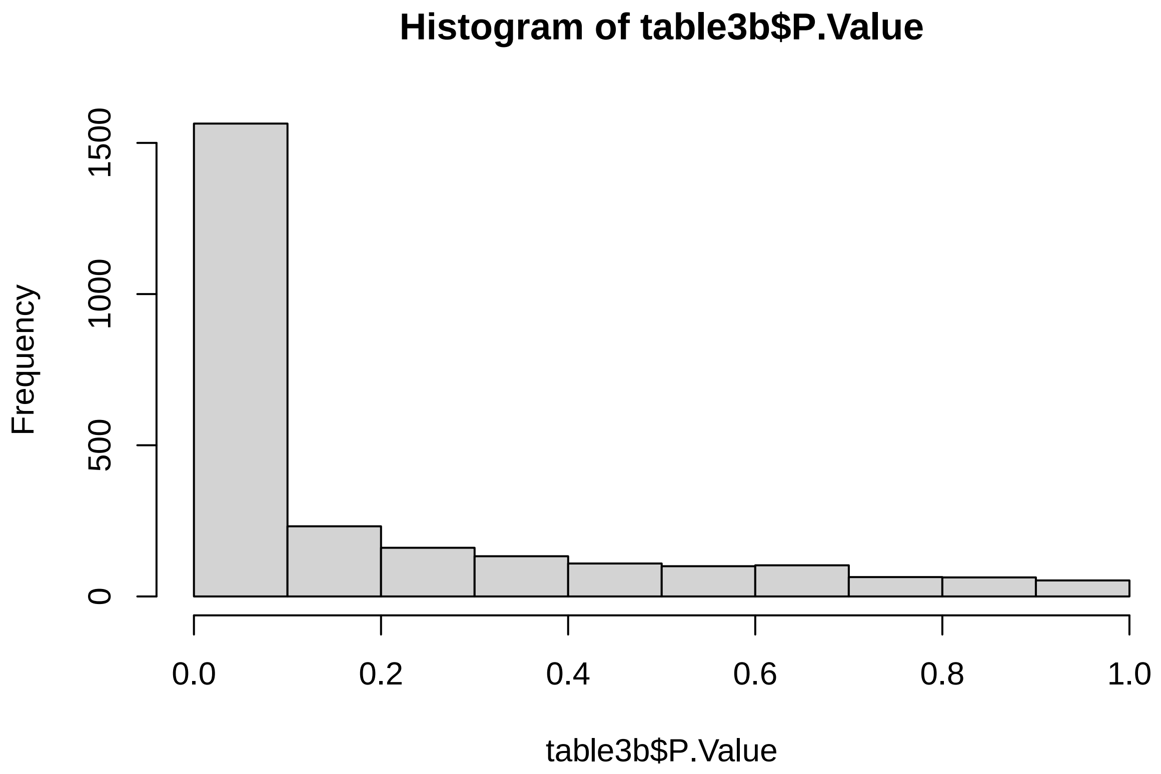

```r
# create design matrix Percent_prot =
# as.numeric(SummarizedExperiment_fathers$Percent_prot) Percent_carb =
# as.numeric(SummarizedExperiment_fathers$Percent_fat)

design4 <- model.matrix(~Percent_prot * Percent_fat + dietary_intake)

data_sample_names <- colnames(SummarizedExperiment_fathers)
design_sample_names <- rownames(design4)

rownames(design4) <- data_sample_names
```

###### 4.0.2.2 Interaction/quadratic model

```r
fit4 <- lmFit(assay(SummarizedExperiment_fathers), design4)
fit4 <- eBayes(fit4)
```

```r
table4_interaction <- topTable(fit4, sort = "none", number = Inf, adjust = "fdr")
count(table4_interaction$adj.P.Val < 0.05)
```

###### 4.0.2.2.1 Number of significant proteins

```
## [1] 1167
```

```r
hist(table4_interaction$P.Value)
```

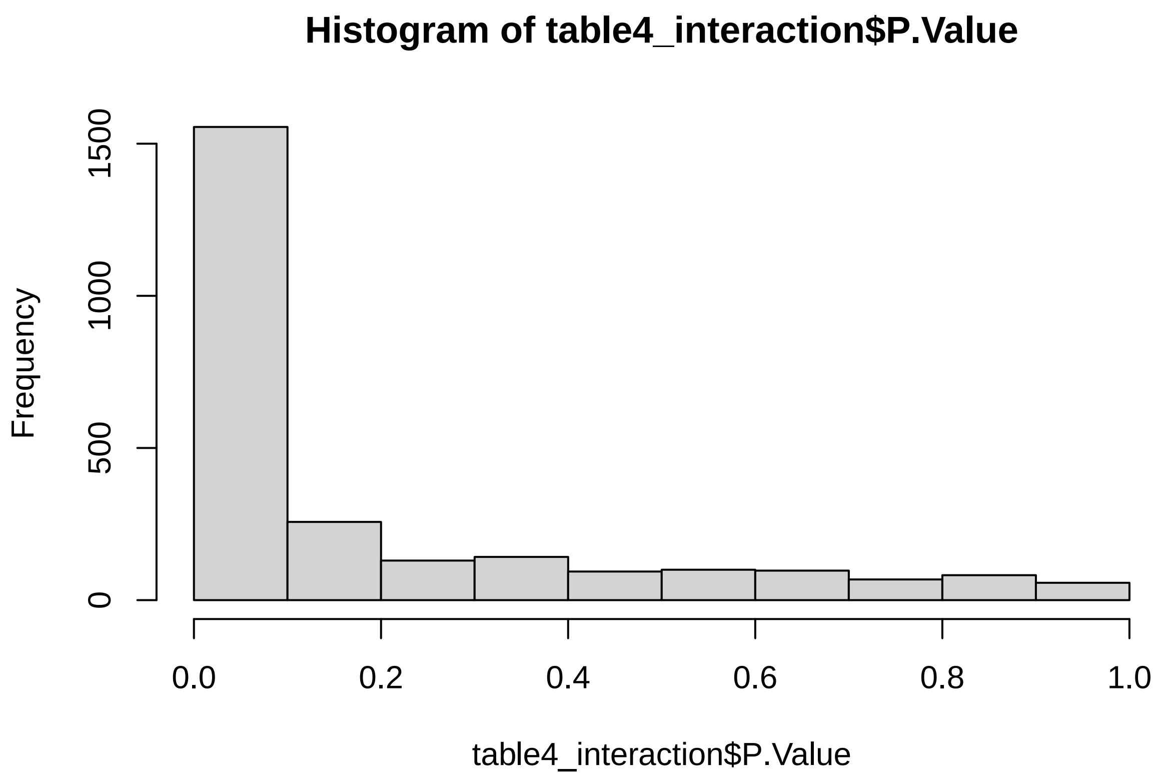

###### 4.0.2.3 Comparing models

1327 proteins were significantly affected by diet in the main text 1262 proteins were signficantly affected by diet with dietary intake as included

```
# This is only taking the rownames
Sig_additive2 <- rownames(table3b[table3b$adj.P.Val < 0.05, ])
Sig_interaction2 <- rownames(table4_interaction[table4_interaction$adj.P.Val < 0.05,
    ])

# find common and different protein

common_proteins2 <- intersect(Sig_additive2, Sig_interaction2)
unique_to_additive2 <- setdiff(Sig_additive2, Sig_interaction2)
unique_to_interaction2 <- setdiff(Sig_interaction2, Sig_additive2)

common2 <- length(common_proteins2)
additive2 <- length(unique_to_additive2)
interaction2 <- length(unique_to_interaction2)
```

```
v2 <- draw.pairwise.venn(area1 = additive2 + common2, area2 = interaction2 + common2,
    cross.area = common2, category = c("Additive Model", "Interaction Model"), fill = c("blue",
        "red"), alpha = 0.5, cex = 1.5, cat.cex = 1.5, cat.pos = c(-20, 20), cat.dist = 0.05,
    0.05, ind = FALSE)

grid.newpage()
grid.draw(v2)
```

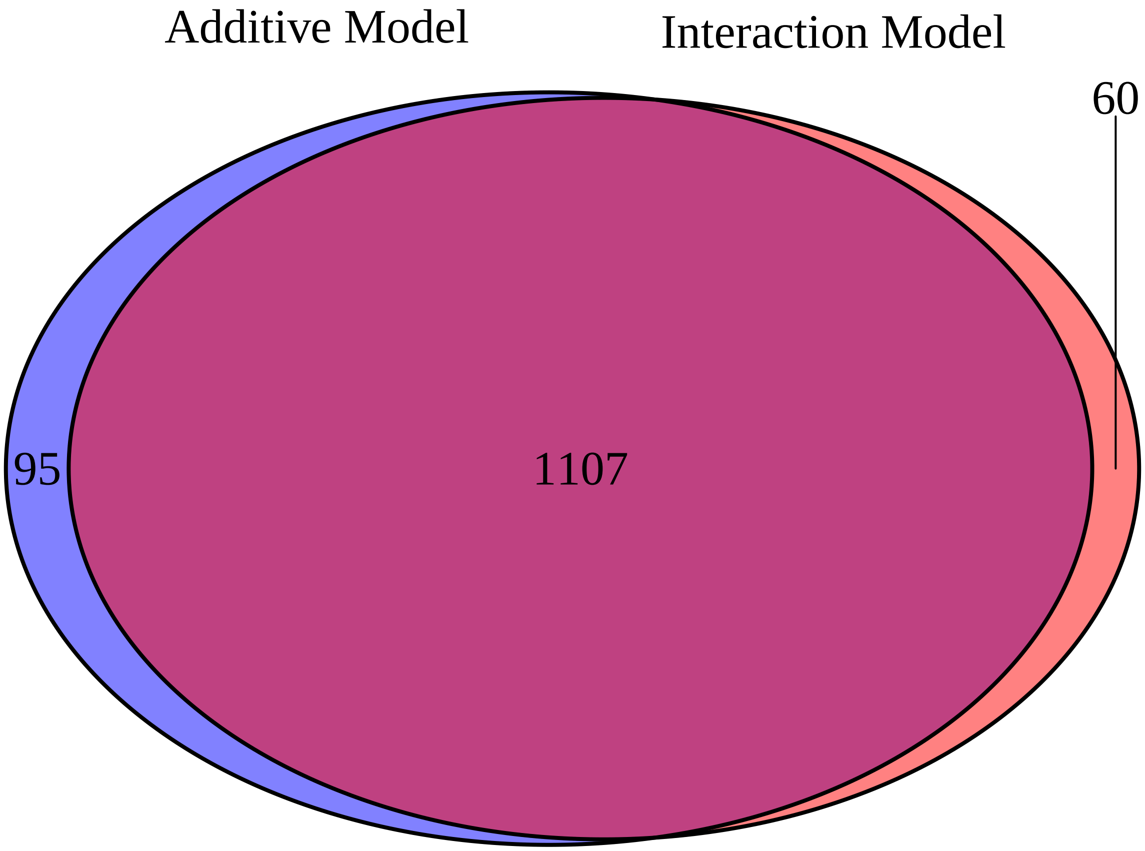

###### 4.0.3 Carbs and fat (%)

```
# create design matrix
Percent_carb = as.numeric(SummarizedExperiment_fathers$Percent_carb)
Percent_fat = as.numeric(SummarizedExperiment_fathers$Percent_fat)
dietary_intake = as.numeric(SummarizedExperiment_fathers$Food_intake)

design5 <- model.matrix(~Percent_carb + Percent_fat + dietary_intake)

data_sample_names <- colnames(SummarizedExperiment_fathers)
design_sample_names <- rownames(design5)

rownames(design5) <- data_sample_names
```

```
fit5 <- lmFit(assay(SummarizedExperiment_fathers), design5)
fit5 <- eBayes(fit5)
table5 <- topTable(fit5, adjust = "fdr", sort.by = "F")
# table8
```

###### 4.0.3.1 Additive model

```r
table5b <- topTable(fit5, sort = "none", number = Inf, adjust = "fdr")

count(table5b$adj.P.Val < 0.05)
```

###### 4.0.3.1.1 Number of significant proteins

```
## [1] 1208
```

```r
hist(table5b$P.Value)
```

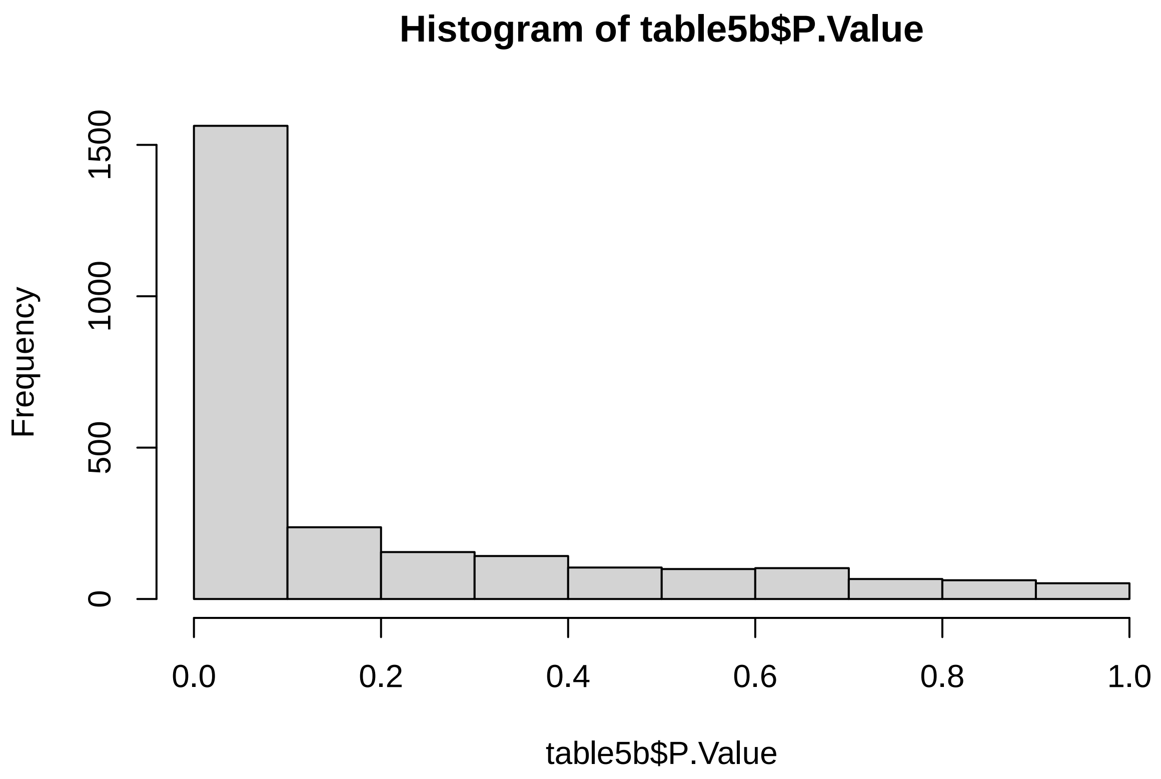

```r
# 1238 DE
```

```r
# #create design matrix Percent_carb =
# as.numeric(SummarizedExperiment_fathers$Percent_carb) Percent_fat =
# as.numeric(SummarizedExperiment_fathers$Percent_fat)

design6 <- model.matrix(~Percent_carb * Percent_fat + dietary_intake)

data_sample_names <- colnames(SummarizedExperiment_fathers)
design_sample_names <- rownames(design6)
```

```r
rownames(design6) <- data_sample_names
```

###### 4.0.3.2 Interaction/quadratic model

```r
fit6 <- lmFit(assay(SummarizedExperiment_fathers), design6)
fit6 <- eBayes(fit6)
```

```r
table6_interaction <- topTable(fit6, sort = "none", number = Inf, adjust = "fdr")
count(table6_interaction$adj.P.Val < 0.05)
```

###### 4.0.3.2.1 Number of significant proteins

```
## [1] 1148
```

```r
hist(table6_interaction$P.Value)
```

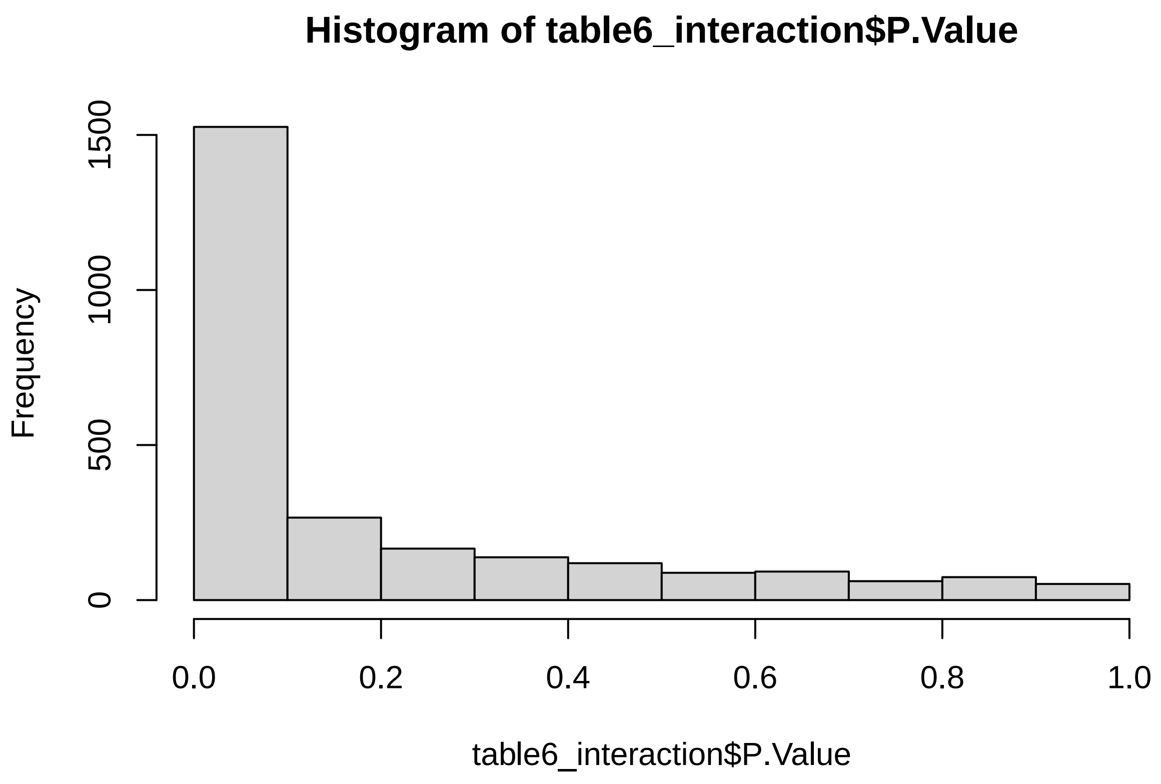

```r
# 1144 DE
```

###### 4.0.3.3 Comparing models

1270 proteins were significantly affected by diet in the main text 1247 proteins were significantly affected by diet when dietary intake was included

```
# This is only taking the rownames
Sig_additive3 <- rownames(table5b[table5b$adj.P.Val < 0.05, ])
Sig_interaction3 <- rownames(table6_interaction[table6_interaction$adj.P.Val < 0.05,
    ])

# find common and different protein

common_proteins3 <- intersect(Sig_additive3, Sig_interaction3)
unique_to_additive3 <- setdiff(Sig_additive3, Sig_interaction3)
unique_to_interaction3 <- setdiff(Sig_interaction3, Sig_additive3)

common3 <- length(common_proteins3)
additive3 <- length(unique_to_additive3)
interaction3 <- length(unique_to_interaction3)
```

```
v3 <- draw.pairwise.venn(area1 = additive3 + common3, area2 = interaction3 + common3,
    cross.area = common3, category = c("Additive Model", "Interaction Model"), fill = c("blue",
        "red"), alpha = 0.5, cex = 1.5, cat.cex = 1.5, cat.pos = c(-20, 20), cat.dist = 0.05,
    0.05, ind = FALSE)

grid.newpage()
grid.draw(v3)
```

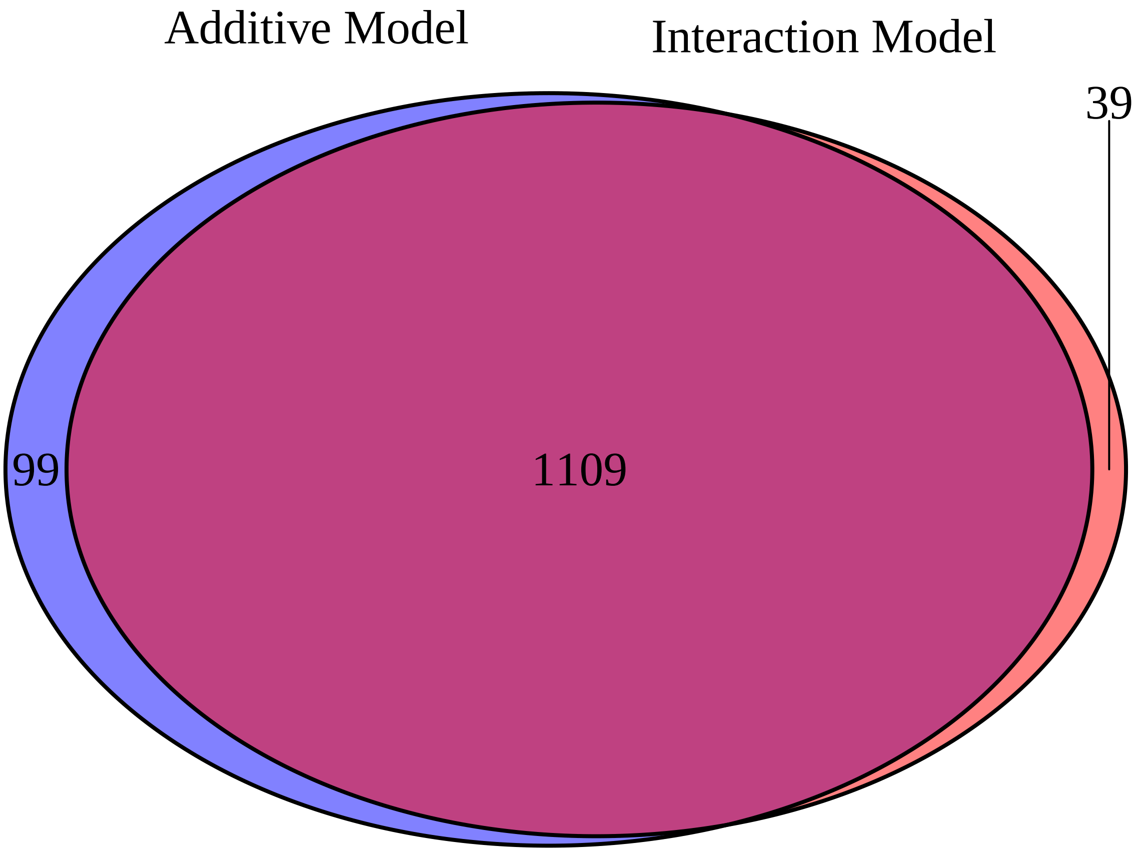

#### 4.1 Clustering analysis

```r
# combing all significant proteins from all models

pPpC_additive <- rownames((table1b[table1b$adj.P.Val < 0.05, ]))
pPpC_interaction <- rownames((table2_interaction[table2_interaction$adj.P.Val < 0.05,
    ]))
pPpF_additive <- rownames((table3b[table3b$adj.P.Val < 0.05, ]))
pPpF_interaction <- rownames((table4_interaction[table4_interaction$adj.P.Val < 0.05,
    ]))
pCpF_additive <- rownames((table5b[table5b$adj.P.Val < 0.05, ]))
pCpF_interaction <- rownames((table6_interaction[table6_interaction$adj.P.Val < 0.05,
    ]))

combined_proteins <- unique(c(pPpC_additive, pPpC_interaction, pPpF_additive, pPpF_interaction,
    pCpF_additive, pCpF_interaction))
```

```r
protein_tibble_fathers <- as_tibble(assay(SummarizedExperiment_fathers), rownames = "proteins")
metadata_fathers <- as.data.frame(colData(SummarizedExperiment_fathers))
log_intensity_data_fathers <- pivot_longer(protein_tibble_fathers, cols = -proteins,
    names_to = "Sample_name", values_to = "LogIntensity")
tidy_protein_table_fathers <- log_intensity_data_fathers %>%
    left_join(metadata_fathers, by = "Sample_name")

Sig_proteins_joined <- tidy_protein_table_fathers[tidy_protein_table_fathers$proteins %in%
    combined_proteins, ]

# converting proportions on a 0 - 1 scale
Sig_proteins_joined$pP <- (Sig_proteins_joined$Percent_prot/100)
Sig_proteins_joined$pC <- (Sig_proteins_joined$Percent_carb/100)
Sig_proteins_joined$pF <- (Sig_proteins_joined$Percent_fat/100)
```

```r
# Loop to run the full model (below) on all proteins (z-transformed) log
# intensity. Clustering is then done on model coefficients

results <- list()

unique_proteins <- unique(Sig_proteins_joined$proteins)

# setting dataframe layout
output_df <- data.frame(Protein = character(length(unique_proteins)), pP = numeric(length(unique_protei[illegible]
    pC = numeric(length(unique_proteins)), pF = numeric(length(unique_proteins)),
    `pP:pC` = numeric(length(unique_proteins)), `pP:pF` = numeric(length(unique_proteins)),
    `pC:pF` = numeric(length(unique_proteins)), stringsAsFactors = FALSE)

for (i in 1:length(unique_proteins)) {
    protein <- unique_proteins[i]

    # making sure that it subsets the data for each protein
    subset_data <- Sig_proteins_joined[Sig_proteins_joined$proteins == protein, ]

    subset_data$LogIntensity_z <- scale(subset_data$LogIntensity)
```

```r
    model <- lm(LogIntensity_z ~ 0 + pP + pC + pF + pP:pC + pP:pF + pC:pF, data = subset_data)

    # extracting the coeficients for each model
    coef_values <- coef(model)

    # data wrangling
    output_df[i, "Protein"] <- protein
    output_df[i, "pP"] <- coef_values["pP"]
    output_df[i, "pC"] <- coef_values["pC"]
    output_df[i, "pF"] <- coef_values["pF"]
    output_df[i, "pP.pC"] <- coef_values["pP:pC"]
    output_df[i, "pP.pF"] <- coef_values["pP:pF"]
    output_df[i, "pC.pF"] <- coef_values["pC:pF"]
}

clustering_data_notz <- output_df[, -1]
```

```r
# fuzzy-c clustering with 5 clusters (based on Farris et al and visual
# inspection of clusters through PCA). Also based on nbClust and kmeans.

clustering_data_notz <- as.matrix(clustering_data_notz)

pca_result <- prcomp(clustering_data_notz, center = TRUE, scale. = TRUE)
pca_scores <- pca_result$x

set.seed(123)  # For reproducibility
fcm_result <- cmeans(clustering_data_notz, centers = 5, iter.max = 100, m = 2)
cluster_assignments <- fcm_result$cluster
# head(cluster_assignments)

# scatterplot3d(pca_scores[, 1], pca_scores[, 2], pca_scores[, 3], color =
# cluster_assignments, pch = 19, xlab = 'PC1', ylab = 'PC2', zlab = 'PC3', main
# = '3D PCA of Protein Data with Fuzzy Clustering') # 3D interactive PCA plot
# with transparent points using plotly plot_ly(x = pca_scores[, 1], y =
# pca_scores[, 2], z = pca_scores[, 3], type = 'scatter3d', mode = 'markers',
# color = as.factor(cluster_assignments), marker = list(size = 3, opacity =
# 0.7)) %>% layout(scene = list(xaxis = list(title = 'PC1'), yaxis = list(title
# = 'PC2'), zaxis = list(title = 'PC3')))
```

```r
# assigning clustering back to data
output_df <- as.data.frame(output_df)

output_df$Cluster <- as.factor(cluster_assignments)
```

```r
protein_tibble_fathers <- as_tibble(assay(SummarizedExperiment_fathers), rownames = "proteins")
metadata_fathers <- as.data.frame(colData(SummarizedExperiment_fathers))
log_intensity_data_fathers <- pivot_longer(protein_tibble_fathers, cols = -proteins,
    names_to = "Sample_name", values_to = "LogIntensity")
tidy_protein_table_fathers <- log_intensity_data_fathers %>%
    left_join(metadata_fathers, by = "Sample_name")

Sig_proteins_joined <- tidy_protein_table_fathers[tidy_protein_table_fathers$proteins %in%
    combined_proteins, ]
```

```r
# converting proportions on a 0 - 1 scale
Sig_proteins_joined$pP <- (Sig_proteins_joined$Percent_prot/100)
Sig_proteins_joined$pC <- (Sig_proteins_joined$Percent_carb/100)
Sig_proteins_joined$pF <- (Sig_proteins_joined$Percent_fat/100)
```

```r
# Loop to run the full model (below) on all proteins (z-transformed) log
# intensity. Clustering is then done on model coefficients

results <- list()

unique_proteins <- unique(Sig_proteins_joined$proteins)

# setting dataframe layout
output_df <- data.frame(Protein = character(length(unique_proteins)), pP = numeric(length(unique_protei
    pC = numeric(length(unique_proteins)), pF = numeric(length(unique_proteins)),
    `pP:pC` = numeric(length(unique_proteins)), `pP:pF` = numeric(length(unique_proteins)),
    `pC:pF` = numeric(length(unique_proteins)), stringsAsFactors = FALSE)

for (i in 1:length(unique_proteins)) {
    protein <- unique_proteins[i]

    # making sure that it subsets the data for each protein
    subset_data <- Sig_proteins_joined[Sig_proteins_joined$proteins == protein, ]

    subset_data$LogIntensity_z <- scale(subset_data$LogIntensity)
    model <- lm(LogIntensity_z ~ 0 + pP + pC + pF + pP:pC + pP:pF + pC:pF, data = subset_data)

    # extracting the coeficients for each model
    coef_values <- coef(model)

    # data wrangling
    output_df[i, "Protein"] <- protein
    output_df[i, "pP"] <- coef_values["pP"]
    output_df[i, "pC"] <- coef_values["pC"]
    output_df[i, "pF"] <- coef_values["pF"]
    output_df[i, "pP.pC"] <- coef_values["pP:pC"]
    output_df[i, "pP.pF"] <- coef_values["pP:pF"]
    output_df[i, "pC.pF"] <- coef_values["pC:pF"]
}

clustering_data_notz <- output_df[, -1]
```

```r
# fuzzy-c clustering with 5 clusters (based on Farris et al and visual
# inspection of clusters through PCA). Also based on nbClust and kmeans.

clustering_data_notz <- as.matrix(clustering_data_notz)

pca_result <- prcomp(clustering_data_notz, center = TRUE, scale. = TRUE)
pca_scores <- pca_result$x

set.seed(123)  # For reproducibility
fcm_result <- cmeans(clustering_data_notz, centers = 5, iter.max = 100, m = 2)
cluster_assignments <- fcm_result$cluster
```

```r
# head(cluster_assignments)

# scatterplot3d(pca_scores[, 1], pca_scores[, 2], pca_scores[, 3], color =
# cluster_assignments, pch = 19, xlab = 'PC1', ylab = 'PC2', zlab = 'PC3', main
# = '3D PCA of Protein Data with Fuzzy Clustering') # 3D interactive PCA plot
# with transparent points using plotly plot_ly(x = pca_scores[, 1], y =
# pca_scores[, 2], z = pca_scores[, 3], type = 'scatter3d', mode = 'markers',
# color = as.factor(cluster_assignments), marker = list(size = 3, opacity =
# 0.7)) %>% layout(scene = list(xaxis = list(title = 'PC1'), yaxis = list(title
# = 'PC2'), zaxis = list(title = 'PC3')))
```

```r
# assigning clustering back to data
output_df <- as.data.frame(output_df)

output_df$Cluster <- as.factor(cluster_assignments)
```

NOTE that is it here that we rename clusters to match patterns in surfaces and the surfaces in the main text - original cluster 1 now = cluster 2 - original cluster 2 now = cluster 1 - original cluster 3 now = cluster 4 - original cluster 4 now = cluster 3

```r
Cluster1 <- output_df[output_df$Cluster == "2", ]
Cluster2 <- output_df[output_df$Cluster == "1", ]
Cluster3 <- output_df[output_df$Cluster == "4", ]
Cluster4 <- output_df[output_df$Cluster == "3", ]
Cluster5 <- output_df[output_df$Cluster == "5", ]

# averaging the coefficients for each term across all proteins

pP_C1_mean <- mean(Cluster1$pP)
pC_C1_mean <- mean(Cluster1$pC)
pF_C1_mean <- mean(Cluster1$pF)
pPpC_C1_mean <- mean(Cluster1$pP.pC)
pPpF_C1_mean <- mean(Cluster1$pP.pF)
pCpF_C1_mean <- mean(Cluster1$pC.pF)

C1_means <- c(pP_C1_mean, pC_C1_mean, pF_C1_mean, pPpC_C1_mean, pPpF_C1_mean, pCpF_C1_mean)

pP_C2_mean <- mean(Cluster2$pP)
pC_C2_mean <- mean(Cluster2$pC)
pF_C2_mean <- mean(Cluster2$pF)
pPpC_C2_mean <- mean(Cluster2$pP.pC)
pPpF_C2_mean <- mean(Cluster2$pP.pF)
pCpF_C2_mean <- mean(Cluster2$pC.pF)

C2_means <- c(pP_C2_mean, pC_C2_mean, pF_C2_mean, pPpC_C2_mean, pPpF_C2_mean, pCpF_C2_mean)

pP_C3_mean <- mean(Cluster3$pP)
pC_C3_mean <- mean(Cluster3$pC)
pF_C3_mean <- mean(Cluster3$pF)
pPpC_C3_mean <- mean(Cluster3$pP.pC)
pPpF_C3_mean <- mean(Cluster3$pP.pF)
pCpF_C3_mean <- mean(Cluster3$pC.pF)
```

```
C3_means <- c(pP_C3_mean, pC_C3_mean, pF_C3_mean, pPpC_C3_mean, pPpF_C3_mean, pCpF_C3_mean)

pP_C4_mean <- mean(Cluster4$pP)
pC_C4_mean <- mean(Cluster4$pC)
pF_C4_mean <- mean(Cluster4$pF)
pPpC_C4_mean <- mean(Cluster4$pP.pC)
pPpF_C4_mean <- mean(Cluster4$pP.pF)
pCpF_C4_mean <- mean(Cluster4$pC.pF)

C4_means <- c(pP_C4_mean, pC_C4_mean, pF_C4_mean, pPpC_C4_mean, pPpF_C4_mean, pCpF_C4_mean)

pP_C5_mean <- mean(Cluster5$pP)
pC_C5_mean <- mean(Cluster5$pC)
pF_C5_mean <- mean(Cluster5$pF)
pPpC_C5_mean <- mean(Cluster5$pP.pC)
pPpF_C5_mean <- mean(Cluster5$pP.pF)
pCpF_C5_mean <- mean(Cluster5$pC.pF)

C5_means <- c(pP_C5_mean, pC_C5_mean, pF_C5_mean, pPpC_C5_mean, pPpF_C5_mean, pCpF_C5_mean)

terms <- c("pP", "pC", "pF", "pPpC", "pPpF", "pCpF")

C1_descriptiveStats <- data.frame(Factor = terms, Mean = C1_means)

C2_descriptiveStats <- data.frame(Factor = terms, Mean = C2_means)

C3_descriptiveStats <- data.frame(Factor = terms, Mean = C3_means)

C4_descriptiveStats <- data.frame(Factor = terms, Mean = C4_means)

C5_descriptiveStats <- data.frame(Factor = terms, Mean = C5_means)
```

```
findConvex.prop <- function(x, y, rgnames, res = 101) {
    hull <- cbind(x, y)[chull(cbind(x, y)), ]
    x.new <- seq(0, 1, len = res)
    y.new <- seq(0, 1, len = res)
    ingrid <- as.data.frame(expand.grid(x.new, y.new))
    Fgrid <- ingrid
    Fgrid[(point.in.polygon(ingrid[, 1], ingrid[, 2], hull[, 1], hull[, 2]) == 0),
        ] <- NA
    names(Fgrid) <- rgnames
    return(Fgrid)
}
```

###### 4.1.1 Cluster 1

```
# need this to create surface
data = metadata
p_P = "Percent_prot"
p_C = "Percent_carb"
```

```r
p_F = "Percent_fat"

coeffs = C1_descriptiveStats$Mean

mixture.surface <- function(data, coeffs, p_P = "p_P", p_C = "p_C", p_F = "p_F") {
}

# Set the resolution of the surface
surface.resolution <- 501

# How many values to round surface
round.surf <- 3

# This specifies the color scheme for surface - it is actually a function that
# returns a function
rgb.palette <- colorRampPalette(c("blue", "cyan", "yellow", "red"), space = "Lab",
    interpolate = "linear")

# How many different colours should we use on the plot
no.cols <- 256

# Get the colors to use from the palette specified above
map <- rgb.palette(no.cols)

# How many levels should there be on the surface
nlev <- 5

# Labels for each
labels <- c("Protein (%)", "Carbohydrate (%)", "Fat (%)")

## plot the RMT surface
iso.lines <- seq(1, 0, -0.2)

# Set the layout
par(mfrow = c(1, 1), mar = c(5, 5, 5, 1))

# Make sure the proportions are closed off to 1
total <- (data[, p_P] + data[, p_C] + data[, p_F])
data$p_P <- data[, p_P]/total
data$p_C <- data[, p_C]/total
data$p_F <- data[, p_F]/total

## estimate convex hull and predict
mdff2 <- findConvex.prop(data$p_P, data$p_C, c("p_P", "p_C"), surface.resolution)
mdff2$p_F <- with(mdff2, 1 - p_P - p_C)

# Get the predicted surface
X <- model.matrix(~0 + p_P + p_C + p_F + p_P:p_C + p_P:p_F + p_C:p_F, data = mdff2)
Y <- (X %*% coeffs)
plot_data <- as.data.frame(X)
plot_data$Y <- Y
```

```r
plot_data$p_P <- plot_data$p_P * 100
plot_data$p_C <- plot_data$p_C * 100
plot_data$p_F <- plot_data$p_F * 100
contour_use <- signif((max(Y) - min(Y))/5, 1)
```

```r
  C1_surface <- ggplot(plot_data) +
  # Add diagonal lines with solid lines, ensuring they cover the whole plot area
  geom_abline(intercept = 100, slope = -1, color = "grey50") +
  geom_abline(intercept = 80, slope = -1, color = "grey50") +
  geom_abline(intercept = 60, slope = -1, color = "grey50") +
  geom_abline(intercept = 40, slope = -1, color = "grey50") +
  geom_abline(intercept = 20, slope = -1, color = "grey50") +

  # Add other plot layers on top of the lines
  geom_tile(aes(x = p_P, y = p_C, fill = Y)) +
  scale_fill_gradientn(colors = map) +
  geom_contour(data = plot_data, aes(x = p_P, y = p_C, z = Y), na.rm = TRUE, color = "black", binwidth =
  # geom_label_contour(data = plot_data, aes(x = p_P, y = p_C, z = Y), size = 3, binwidth = contour_use

  # Set theme and labels, remove grid lines and specific axis lines
  theme_minimal() +
  theme(
    panel.grid = element_blank(),       # Remove grid lines
    panel.border = element_blank(),     # Remove the border around the plot panel
    axis.line = element_line(color = "black"),       # Add back the x and y axis lines
    legend.position = "none",            # Remove legend
    axis.text.x = element_text(hjust = -1, vjust = -0.5),  # Adjust horizontal and vertical position fo
    axis.text.y = element_text(hjust = 0.5, vjust = -1.5)    # Adjust horizontal and vertical position
  ) +
  scale_x_continuous(limits = c(0, 100), breaks = seq(0, 100, by = 20)) +  # Set x-axis limits and labe
  scale_y_continuous(limits = c(0, 100), breaks = seq(0, 100, by = 20)) +  # Set y-axis limits and labe
  # coord_fixed(ratio = 1) +   # Fix the aspect ratio to ensure alignment of diagonals and axes
  ggtitle("Cluster 1 (255 proteins)") +
  labs(x = "Protein (%)", y = "Carbohydrate (%)") +

  # Adjust the "Fat" label position
  annotate("text", x = 55, y = 55, label = "Fat (%)", color = "black", angle = -45, hjust = 1)

# Display the plot
C1_surface
```

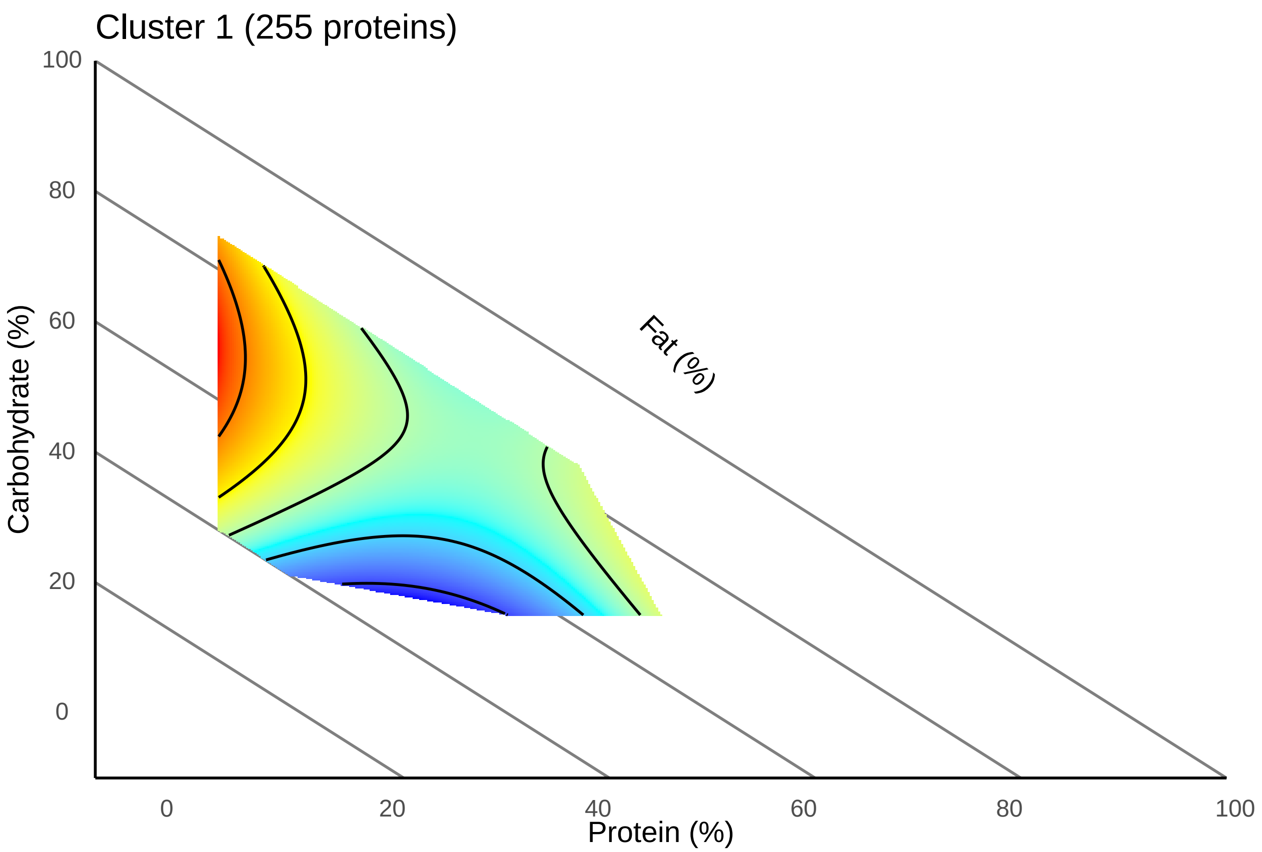

###### 4.1.1.1 Average surface

```
valid_ids <- keys(org.Mm.eg.db, keytype = "ENTREZID")

protein_list <- Cluster1$Protein
background <- protein_tibble_fathers$proteins

# Convert Uniprot IDs to Entrez IDs
entrez_ids_c1 <- bitr(protein_list, fromType = "SYMBOL", toType = "ENTREZID", OrgDb = "org.Mm.eg.db")
# 1.69% of genes failed to map

# Extract the Entrez IDs
entrez_list <- entrez_ids_c1$ENTREZID

# removing missing values
entrez_list <- entrez_list[!is.na(entrez_list) & entrez_list != ""]
entrez_list <- entrez_list[entrez_list %in% valid_ids]

entrez_ids_background <- bitr(background, fromType = "SYMBOL", toType = "ENTREZID",
    OrgDb = "org.Mm.eg.db")

# Extract the Entrez IDs
entrez_list_background <- entrez_ids_background$ENTREZID

# removing missing values
```

```r
entrez_list_background <- entrez_list_background[!is.na(entrez_list_background) &
    entrez_list_background != ""]
entrez_list_background <- entrez_list_background[entrez_list_background %in% valid_ids]

# GO analysis go_cluster1_all <- enrichGO(gene = entrez_list, OrgDb =
# org.Mm.eg.db, keyType = 'ENTREZID', ont = 'all', universe =
# entrez_list_background, pAdjustMethod = 'BH', pvalueCutoff = 0.05) #
# go_cluster1_all_simplified <- simplify(go_cluster1_all, cutoff = 0.7, by =
# 'p.adjust', select_fun = min, measure = 'Wang')

go_cluster1_BP <- enrichGO(gene = entrez_list, OrgDb = org.Mm.eg.db, keyType = "ENTREZID",
    ont = "BP", universe = entrez_list_background, pAdjustMethod = "BH", pvalueCutoff = 0.05)
#
go_cluster1_BP_simplified <- clusterProfiler::simplify(go_cluster1_BP, cutoff = 0.7,
    by = "p.adjust", select_fun = min, measure = "Wang")
```

**4.1.1.2 Enrichment analysis** Lower number of proteins involved in enriched pathways (lower gene ratios and p-values)

```r
# GO_cluster1 <-clusterProfiler::dotplot(merge_result(list( 'Cellular
# Component' = go_cluster1_all_simplified)), showCategory = 5) + xlab(NULL)
# GO_cluster1 <- clusterProfiler::dotplot(go_cluster1_all_simplified,
# showCategory = 5)

# GO_cluster1

bar_data_c1 <- go_cluster1_BP_simplified@result
top_bar_data_c1 <- bar_data_c1[order(bar_data_c1$p.adjust), ][1:5, ]
top_bar_data_c1$log10_p_adjust <- -log10(top_bar_data_c1$p.adjust)
top_bar_data_c1$percent_proteins <- ((top_bar_data_c1$Count/211) * 100)

# top_bar_data <- bar_data[order(bar_data$log10_p_adjust), ][1:10]

# enrichedgenes_cluster1 <-
# unique(unlist(strsplit(go_cluster1_BP_simplified$geneID, '/')))
# length(enrichedgenes_cluster1)

C1_bar <- ggplot(top_bar_data_c1, aes(x = log10_p_adjust, y = reorder(Description,
    log10_p_adjust), fill = percent_proteins)) + geom_bar(stat = "identity") + xlim(0,
    10) + scale_fill_gradientn(colors = c("lightblue", "blue", "darkblue"), limits = c(0,
    50), breaks = seq(0, 60, by = 20)) + labs(x = NULL, y = NULL, fill = NULL) +
    theme_minimal() + ggtitle("Cluster 1 (61 proteins)") + theme(axis.text.y = element_text(size = 12),
    axis.text.x = element_text(size = 12), legend.position = "none")
C1_bar
```

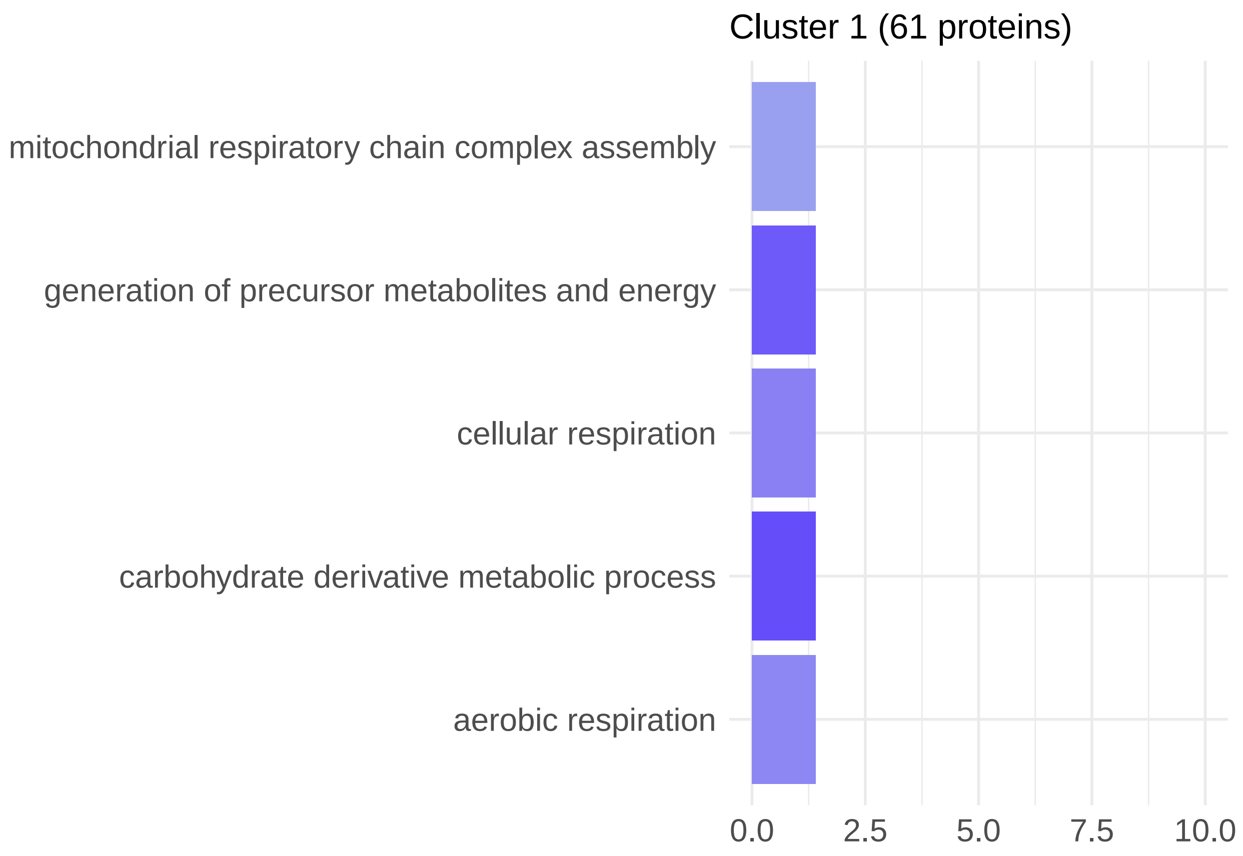

##### 4.1.2 Cluster 2

```
# need this to create surface
data = metadata
p_P = "Percent_prot"
p_C = "Percent_carb"
p_F = "Percent_fat"

coeffs = C2_descriptiveStats$Mean

mixture.surface <- function(data, coeffs, p_P = "p_P", p_C = "p_C", p_F = "p_F") {
}

# Set the resolution of the surface
surface.resolution <- 501

# How many values to round surface
round.surf <- 3

# This specifies the color scheme for surface - it is actually a function that
# returns a function
rgb.palette <- colorRampPalette(c("blue", "cyan", "yellow", "red"), space = "Lab",
    interpolate = "linear")
```

```r
# How many different colours should we use on the plot
no.cols <- 256

# Get the colors to use from the palette specified above
map <- rgb.palette(no.cols)

# How many levels should there be on the surface
nlev <- 5

# Labels for each
labels <- c("Protein (%)", "Carbohydrate (%)", "Fat (%)")

## plot the RMT surface
iso.lines <- seq(1, 0, -0.2)

# Set the layout
par(mfrow = c(1, 1), mar = c(5, 5, 5, 1))

# Make sure the proportions are closed off to 1
total <- (data[, p_P] + data[, p_C] + data[, p_F])
data$p_P <- data[, p_P]/total
data$p_C <- data[, p_C]/total
data$p_F <- data[, p_F]/total

## estimate convex hull and predict
mdff2 <- findConvex.prop(data$p_P, data$p_C, c("p_P", "p_C"), surface.resolution)
mdff2$p_F <- with(mdff2, 1 - p_P - p_C)

# Get the predicted surface
X <- model.matrix(~0 + p_P + p_C + p_F + p_P:p_C + p_P:p_F + p_C:p_F, data = mdff2)
Y <- (X %*% coeffs)
plot_data <- as.data.frame(X)
plot_data$Y <- Y
plot_data$p_P <- plot_data$p_P * 100
plot_data$p_C <- plot_data$p_C * 100
plot_data$p_F <- plot_data$p_F * 100
contour_use <- signif((max(Y) - min(Y))/5, 1)
```

```r
  C2_surface <- ggplot(plot_data) +
  # Add diagonal lines with solid lines, ensuring they cover the whole plot area
  geom_abline(intercept = 100, slope = -1, color = "grey50") +
  geom_abline(intercept = 80, slope = -1, color = "grey50") +
  geom_abline(intercept = 60, slope = -1, color = "grey50") +
  geom_abline(intercept = 40, slope = -1, color = "grey50") +
  geom_abline(intercept = 20, slope = -1, color = "grey50") +

  # Add other plot layers on top of the lines
  geom_tile(aes(x = p_P, y = p_C, fill = Y)) +
  scale_fill_gradientn(colors = map) +
```

```
  geom_contour(data = plot_data, aes(x = p_P, y = p_C, z = Y), na.rm = TRUE, color = "black", binwidth =
  # geom_label_contour(data = plot_data, aes(x = p_P, y = p_C, z = Y), size = 3, binwidth = contour_use

  # Set theme and labels, remove grid lines and specific axis lines
  theme_minimal() +
  theme(
    panel.grid = element_blank(),        # Remove grid lines
    panel.border = element_blank(),      # Remove the border around the plot panel
    axis.line = element_line(color = "black"),        # Add back the x and y axis lines
    legend.position = "none",              # Remove legend
    axis.text.x = element_text(hjust = -1, vjust = -0.5),  # Adjust horizontal and vertical position fo
    axis.text.y = element_text(hjust = 0.5, vjust = -1.5)    # Adjust horizontal and vertical position
  ) +
  scale_x_continuous(limits = c(0, 100), breaks = seq(0, 100, by = 20)) +  # Set x-axis limits and labe
  scale_y_continuous(limits = c(0, 100), breaks = seq(0, 100, by = 20)) +  # Set y-axis limits and labe
  # coord_fixed(ratio = 1) +   # Fix the aspect ratio to ensure alignment of diagonals and axes
  ggtitle("Cluster 2 (269 proteins)") +
  labs(x = "Protein (%)", y = "Carbohydrate (%)") +

  # Adjust the "Fat" label position
  annotate("text", x = 55, y = 55, label = "Fat (%)", color = "black", angle = -45, hjust = 1)

# Display the plot
C2_surface
```

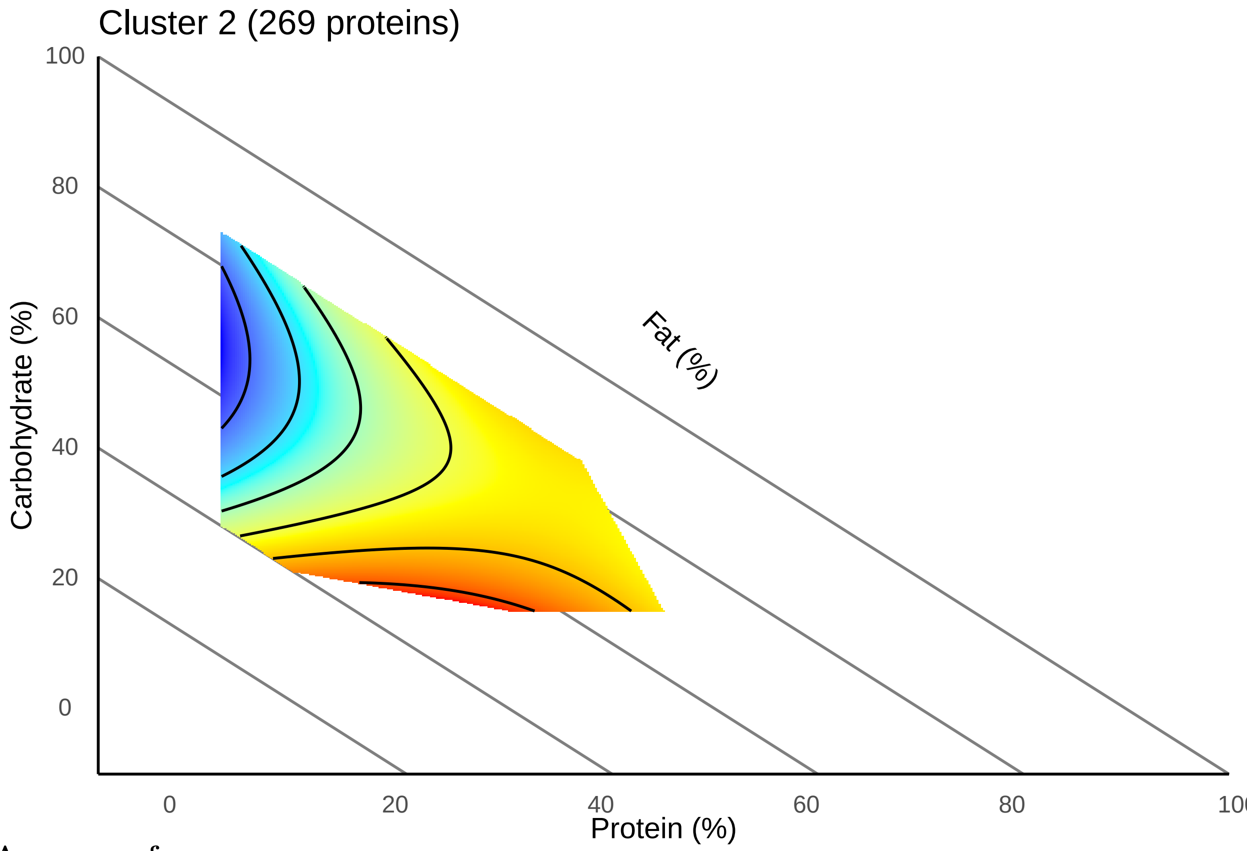

###### 4.1.2.1 Average surface

###### 4.1.2.2 Enrichment analysis

We now detect significantly enriched BPs in this cluster

```
valid_ids <- keys(org.Mm.eg.db, keytype = "ENTREZID")

protein_list <- Cluster2$Protein
background <- protein_tibble_fathers$proteins

# Convert Uniprot IDs to Entrez IDs
entrez_ids_c2 <- bitr(protein_list, fromType = "SYMBOL", toType = "ENTREZID", OrgDb = "org.Mm.eg.db")

# Extract the Entrez IDs
entrez_list <- entrez_ids_c2$ENTREZID

# removing missing values
entrez_list <- entrez_list[!is.na(entrez_list) & entrez_list != ""]
entrez_list <- entrez_list[entrez_list %in% valid_ids]

entrez_ids_background <- bitr(background, fromType = "SYMBOL", toType = "ENTREZID",
    OrgDb = "org.Mm.eg.db")

# Extract the Entrez IDs
entrez_list_background <- entrez_ids_background$ENTREZID

# removing missing values
entrez_list_background <- entrez_list_background[!is.na(entrez_list_background) &
    entrez_list_background != ""]
entrez_list_background <- entrez_list_background[entrez_list_background %in% valid_ids]

# GO analysis go_cluster2_all <- enrichGO(gene = entrez_list, OrgDb =
# org.Mm.eg.db, keyType = 'ENTREZID', ont = 'all', universe =
# entrez_list_background, pAdjustMethod = 'BH', pvalueCutoff = 0.05) #
# go_cluster2_all_simplified <- simplify(go_cluster2_all, cutoff = 0.7, by =
# 'p.adjust', select_fun = min, measure = 'Wang')

go_cluster2_BP <- enrichGO(gene = entrez_list, OrgDb = org.Mm.eg.db, keyType = "ENTREZID",
    ont = "BP", universe = entrez_list_background, pAdjustMethod = "BH", pvalueCutoff = 0.05)
#
go_cluster2_BP_simplified <- clusterProfiler::simplify(go_cluster2_BP, cutoff = 0.7,
    by = "p.adjust", select_fun = min, measure = "Wang")
```

```
bar_data_c2 <- go_cluster2_BP_simplified@result
top_bar_data_c2 <- bar_data_c2[order(bar_data_c2$p.adjust), ][1:5, ]
top_bar_data_c2$log10_p_adjust <- -log10(top_bar_data_c2$p.adjust)
top_bar_data_c2$percent_proteins <- ((top_bar_data_c2$Count/242) * 100)

# top_bar_data <- bar_data[order(bar_data$log10_p_adjust), ][1:10]

# enrichedgenes_cluster2 <-
# unique(unlist(strsplit(go_cluster2_BP_simplified$geneID, '/')))
# length(enrichedgenes_cluster2)

C2_bar <- ggplot(top_bar_data_c2, aes(x = log10_p_adjust, y = reorder(Description,
    log10_p_adjust), fill = percent_proteins)) + geom_bar(stat = "identity") + xlim(0,
```

```
    10) + scale_fill_gradientn(colors = c("lightblue", "blue", "darkblue"), limits = c(0,
    50), breaks = seq(0, 60, by = 20)) + labs(x = NULL, y = NULL, fill = NULL) +
    theme_minimal() + ggtitle("Cluster 1 (104 proteins)") + theme(axis.text.y = element_text(size = 12)
    axis.text.x = element_text(size = 12), legend.position = "none")
C2_bar
```

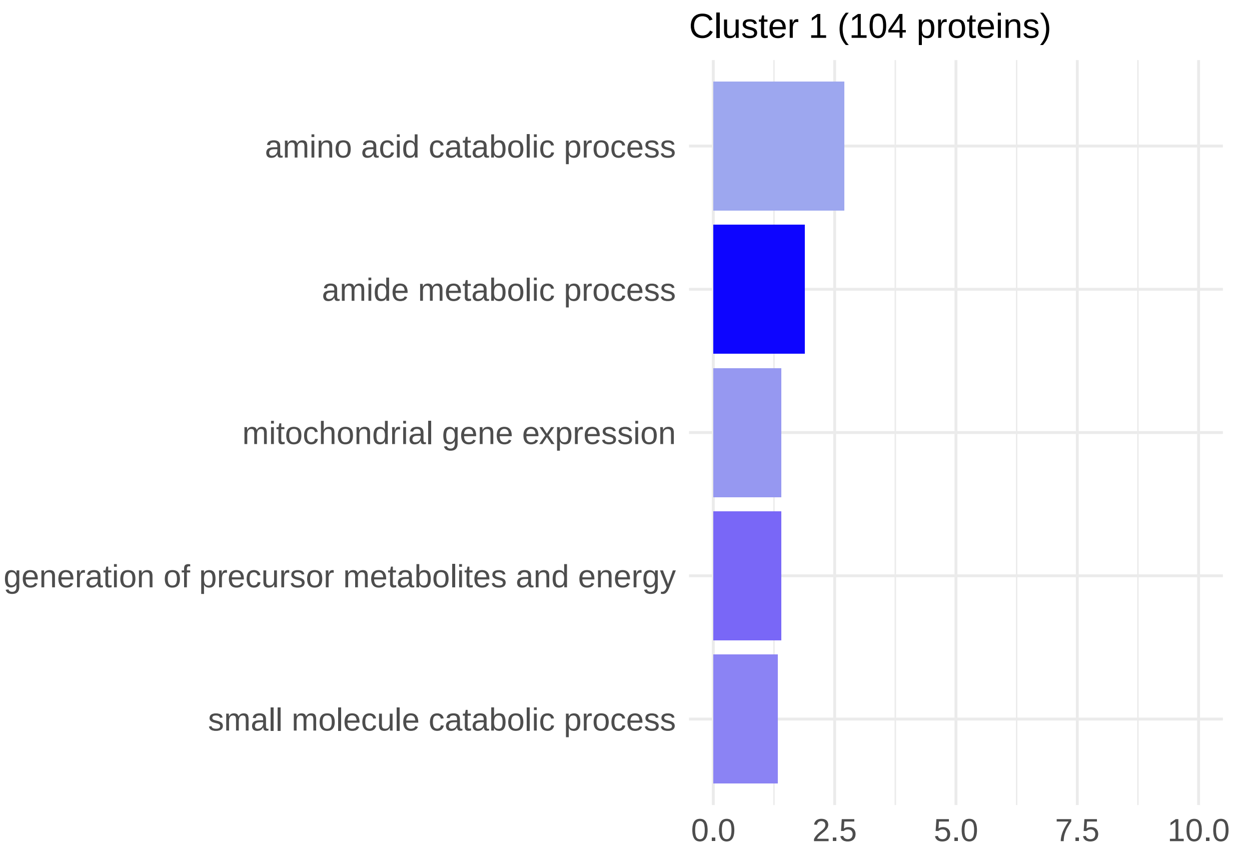

##### 4.1.3 Cluster 3

```
# need this to create surface
data = metadata
p_P = "Percent_prot"
p_C = "Percent_carb"
p_F = "Percent_fat"

coeffs = C3_descriptiveStats$Mean

mixture.surface <- function(data, coeffs, p_P = "p_P", p_C = "p_C", p_F = "p_F") {
}

# Set the resolution of the surface
surface.resolution <- 501
```

```r
# How many values to round surface
round.surf <- 3

# This specifies the color scheme for surface - it is actually a function that
# returns a function
rgb.palette <- colorRampPalette(c("blue", "cyan", "yellow", "red"), space = "Lab",
    interpolate = "linear")

# How many different colours should we use on the plot
no.cols <- 256

# Get the colors to use from the palette specified above
map <- rgb.palette(no.cols)

# How many levels should there be on the surface
nlev <- 5

# Labels for each
labels <- c("Protein (%)", "Carbohydrate (%)", "Fat (%)")

## plot the RMT surface
iso.lines <- seq(1, 0, -0.2)

# Set the layout
par(mfrow = c(1, 1), mar = c(5, 5, 5, 1))

# Make sure the proportions are closed off to 1
total <- (data[, p_P] + data[, p_C] + data[, p_F])
data$p_P <- data[, p_P]/total
data$p_C <- data[, p_C]/total
data$p_F <- data[, p_F]/total

## estimate convex hull and predict
mdff2 <- findConvex.prop(data$p_P, data$p_C, c("p_P", "p_C"), surface.resolution)
mdff2$p_F <- with(mdff2, 1 - p_P - p_C)

# Get the predicted surface
X <- model.matrix(~0 + p_P + p_C + p_F + p_P:p_C + p_P:p_F + p_C:p_F, data = mdff2)
Y <- (X %*% coeffs)
plot_data <- as.data.frame(X)
plot_data$Y <- Y
plot_data$p_P <- plot_data$p_P * 100
plot_data$p_C <- plot_data$p_C * 100
plot_data$p_F <- plot_data$p_F * 100
contour_use <- signif((max(Y) - min(Y))/5, 1)
```

```r

  C3_surface <- ggplot(plot_data) +
  # Add diagonal lines with solid lines, ensuring they cover the whole plot area
  geom_abline(intercept = 100, slope = -1, color = "grey50") +
  geom_abline(intercept = 80, slope = -1, color = "grey50") +
```

```
  geom_abline(intercept = 60, slope = -1, color = "grey50") +
  geom_abline(intercept = 40, slope = -1, color = "grey50") +
  geom_abline(intercept = 20, slope = -1, color = "grey50") +

  # Add other plot layers on top of the lines
  geom_tile(aes(x = p_P, y = p_C, fill = Y)) +
  scale_fill_gradientn(colors = map) +
  geom_contour(data = plot_data, aes(x = p_P, y = p_C, z = Y), na.rm = TRUE, color = "black", binwidth =
  # geom_label_contour(data = plot_data, aes(x = p_P, y = p_C, z = Y), size = 3, binwidth = contour_use

  # Set theme and labels, remove grid lines and specific axis lines
  theme_minimal() +
  theme(
    panel.grid = element_blank(),       # Remove grid lines
    panel.border = element_blank(),     # Remove the border around the plot panel
    axis.line = element_line(color = "black"),       # Add back the x and y axis lines
    legend.position = "none",            # Remove legend
    axis.text.x = element_text(hjust = -1, vjust = -0.5),  # Adjust horizontal and vertical position fo
    axis.text.y = element_text(hjust = 0.5, vjust = -1.5)    # Adjust horizontal and vertical position
  ) +
  scale_x_continuous(limits = c(0, 100), breaks = seq(0, 100, by = 20)) +  # Set x-axis limits and labe
  scale_y_continuous(limits = c(0, 100), breaks = seq(0, 100, by = 20)) +  # Set y-axis limits and labe
  # coord_fixed(ratio = 1) +   # Fix the aspect ratio to ensure alignment of diagonals and axes
  ggtitle("Cluster 3 (239 proteins)") +
  labs(x = "Protein (%)", y = "Carbohydrate (%)") +

  # Adjust the "Fat" label position
  annotate("text", x = 55, y = 55, label = "Fat (%)", color = "black", angle = -45, hjust = 1)

# Display the plot
C3_surface
```

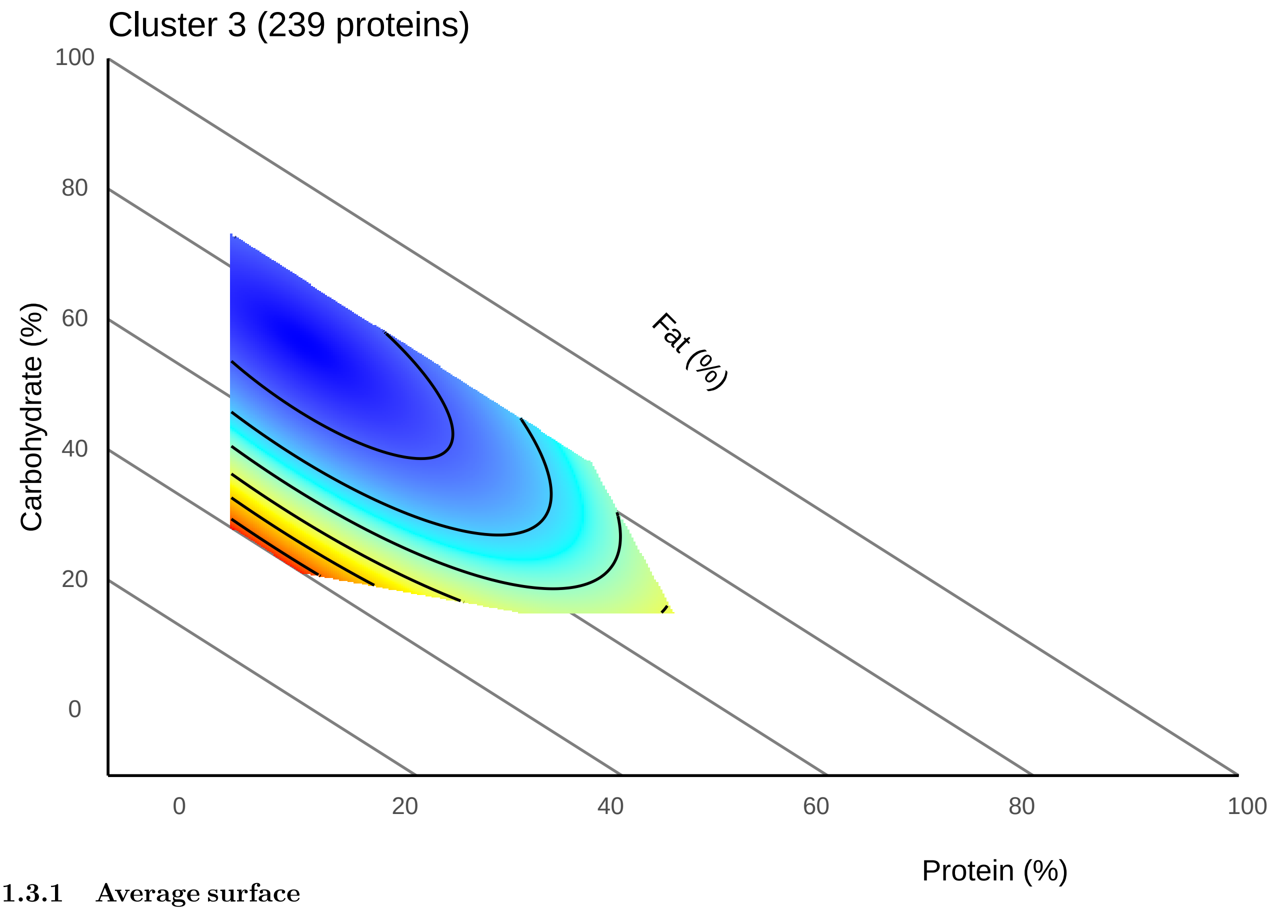

###### 4.1.3.1 Average surface

```
valid_ids <- keys(org.Mm.eg.db, keytype = "ENTREZID")

protein_list <- Cluster3$Protein
background <- protein_tibble_fathers$proteins

# Convert Uniprot IDs to Entrez IDs
entrez_ids_c3 <- bitr(protein_list, fromType = "SYMBOL", toType = "ENTREZID", OrgDb = "org.Mm.eg.db")

# Extract the Entrez IDs
entrez_list <- entrez_ids_c3$ENTREZID

# removing missing values
entrez_list <- entrez_list[!is.na(entrez_list) & entrez_list != ""]
entrez_list <- entrez_list[entrez_list %in% valid_ids]

entrez_ids_background <- bitr(background, fromType = "SYMBOL", toType = "ENTREZID",
    OrgDb = "org.Mm.eg.db")

# Extract the Entrez IDs
entrez_list_background <- entrez_ids_background$ENTREZID

# removing missing values
entrez_list_background <- entrez_list_background[!is.na(entrez_list_background) &
```

```
    entrez_list_background != ""]
entrez_list_background <- entrez_list_background[entrez_list_background %in% valid_ids]

# GO analysis go_cluster3_all <- enrichGO(gene = entrez_list, OrgDb =
# org.Mm.eg.db, keyType = 'ENTREZID', ont = 'all', universe =
# entrez_list_background, pAdjustMethod = 'BH', pvalueCutoff = 0.05)
# go_cluster3_all_simplified <- simplify(go_cluster3_all, cutoff = 0.7, by =
# 'p.adjust', select_fun = min, measure = 'Wang')

go_cluster3_BP <- enrichGO(gene = entrez_list, OrgDb = org.Mm.eg.db, keyType = "ENTREZID",
    ont = "BP", universe = entrez_list_background, pAdjustMethod = "BH", pvalueCutoff = 0.05)

go_cluster3_BP_simplified <- clusterProfiler::simplify(go_cluster3_BP, cutoff = 0.7,
    by = "p.adjust", select_fun = min, measure = "Wang")
# nothing when we simplify

# go_cluster3_CC <- enrichGO(gene = entrez_list, OrgDb = org.Mm.eg.db, keyType
# = 'ENTREZID', ont = 'CC', universe = entrez_list_background, pAdjustMethod =
# 'BH', pvalueCutoff = 0.05) go_cluster3_CC_simplified <-
# simplify(go_cluster3_CC, cutoff = 0.7, by = 'p.adjust', select_fun = min,
# measure = 'Wang') go_cluster3_MF <- enrichGO(gene = entrez_list, OrgDb =
# org.Mm.eg.db, keyType = 'ENTREZID', ont = 'MF', universe =
# entrez_list_background, pAdjustMethod = 'BH', pvalueCutoff = 0.05) #
# go_cluster3_MF_simplified <- simplify(go_cluster3_MF, cutoff = 0.7, by =
# 'p.adjust', select_fun = min, measure = 'Wang')
```

```
# GO_cluster3 <-clusterProfiler::dotplot(merge_result(list( 'Biological
# Process' = go_cluster3_BP_simplified,'Cellular Component' =
# go_cluster3_CC_simplified, 'Molecular Function' =
# go_cluster3_BP_simplified)), showCategory = 5) + xlab(NULL) # GO_cluster3 <-
# clusterProfiler::dotplot(go_cluster3_all_simplified, showCategory =5)
# GO_cluster3

bar_data_c3 <- go_cluster3_BP@result
top_bar_data_c3 <- bar_data_c3[order(bar_data_c3$p.adjust), ][1:5, ]
top_bar_data_c3$log10_p_adjust <- -log10(top_bar_data_c3$p.adjust)
top_bar_data_c3$percent_proteins <- ((top_bar_data_c3$Count/236) * 100)
# enrichedgenes_cluster3 <-
# unique(unlist(strsplit(go_cluster3_BP_simplified$geneID, '/')))
# length(enrichedgenes_cluster3)

C3_bar <- ggplot(top_bar_data_c3, aes(x = log10_p_adjust, y = reorder(Description,
    log10_p_adjust), fill = percent_proteins)) + geom_bar(stat = "identity") + xlim(0,
    10) + scale_fill_gradientn(colors = c("lightblue", "blue", "darkblue"), limits = c(0,
    50), breaks = seq(0, 60, by = 20)) + labs(x = NULL, y = NULL, fill = "Gene ratio") +
    theme_minimal() + ggtitle("Cluster 3 (175 proteins)") + theme(axis.text.y = element_text(size = 12)
    axis.text.x = element_text(size = 12), legend.position = "none")
C3_bar
```

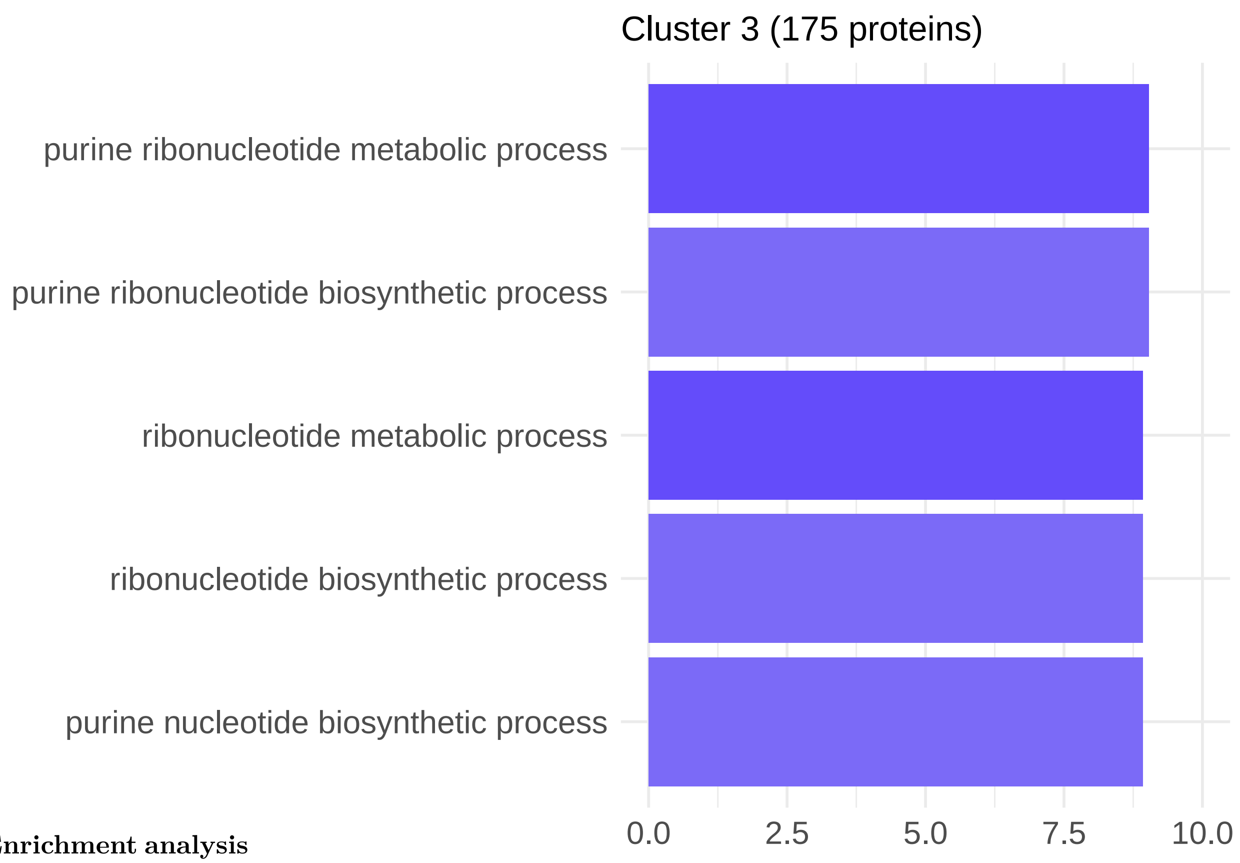

###### 4.1.3.2 Enrichment analysis

##### 4.1.4 Cluster 4

```
# need this to create surface
data = metadata
p_P = "Percent_prot"
p_C = "Percent_carb"
p_F = "Percent_fat"

coeffs = C4_descriptiveStats$Mean

mixture.surface <- function(data, coeffs, p_P = "p_P", p_C = "p_C", p_F = "p_F") {
}

# Set the resolution of the surface
surface.resolution <- 501

# How many values to round surface
round.surf <- 3

# This specifies the color scheme for surface - it is actually a function that
# returns a function
rgb.palette <- colorRampPalette(c("blue", "cyan", "yellow", "red"), space = "Lab",
    interpolate = "linear")
```

```r
# How many different colours should we use on the plot
no.cols <- 256

# Get the colors to use from the palette specified above
map <- rgb.palette(no.cols)

# How many levels should there be on the surface
nlev <- 5

# Labels for each
labels <- c("Protein (%)", "Carbohydrate (%)", "Fat (%)")

## plot the RMT surface
iso.lines <- seq(1, 0, -0.2)

# Set the layout
par(mfrow = c(1, 1), mar = c(5, 5, 5, 1))

# Make sure the proportions are closed off to 1
total <- (data[, p_P] + data[, p_C] + data[, p_F])
data$p_P <- data[, p_P]/total
data$p_C <- data[, p_C]/total
data$p_F <- data[, p_F]/total

## estimate convex hull and predict
mdff2 <- findConvex.prop(data$p_P, data$p_C, c("p_P", "p_C"), surface.resolution)
mdff2$p_F <- with(mdff2, 1 - p_P - p_C)

# Get the predicted surface
X <- model.matrix(~0 + p_P + p_C + p_F + p_P:p_C + p_P:p_F + p_C:p_F, data = mdff2)
Y <- (X %*% coeffs)
plot_data <- as.data.frame(X)
plot_data$Y <- Y
plot_data$p_P <- plot_data$p_P * 100
plot_data$p_C <- plot_data$p_C * 100
plot_data$p_F <- plot_data$p_F * 100
contour_use <- signif((max(Y) - min(Y))/5, 1)
```

```r
  C4_surface <- ggplot(plot_data) +
  # Add diagonal lines with solid lines, ensuring they cover the whole plot area
  geom_abline(intercept = 100, slope = -1, color = "grey50") +
  geom_abline(intercept = 80, slope = -1, color = "grey50") +
  geom_abline(intercept = 60, slope = -1, color = "grey50") +
  geom_abline(intercept = 40, slope = -1, color = "grey50") +
  geom_abline(intercept = 20, slope = -1, color = "grey50") +

  # Add other plot layers on top of the lines
  geom_tile(aes(x = p_P, y = p_C, fill = Y)) +
  scale_fill_gradientn(colors = map) +
  geom_contour(data = plot_data, aes(x = p_P, y = p_C, z = Y), na.rm = TRUE, color = "black", binwidth =
  # geom_label_contour(data = plot_data, aes(x = p_P, y = p_C, z = Y), size = 3, binwidth = contour_use
```

```
  # Set theme and labels, remove grid lines and specific axis lines
  theme_minimal() +
  theme(
    panel.grid = element_blank(),       # Remove grid lines
    panel.border = element_blank(),     # Remove the border around the plot panel
    axis.line = element_line(color = "black"),       # Add back the x and y axis lines
    legend.position = "none",              # Remove legend
    axis.text.x = element_text(hjust = -1, vjust = -0.5),  # Adjust horizontal and vertical position fo
    axis.text.y = element_text(hjust = 0.5, vjust = -1.5)    # Adjust horizontal and vertical position 
  ) +
  scale_x_continuous(limits = c(0, 100), breaks = seq(0, 100, by = 20)) +  # Set x-axis limits and labe
  scale_y_continuous(limits = c(0, 100), breaks = seq(0, 100, by = 20)) +  # Set y-axis limits and labe
  # coord_fixed(ratio = 1) +   # Fix the aspect ratio to ensure alignment of diagonals and axes
  ggtitle("Cluster 4 (362 proteins)") +
  labs(x = "Protein (%)", y = "Carbohydrate (%)") +

  # Adjust the "Fat" label position
  annotate("text", x = 55, y = 55, label = "Fat (%)", color = "black", angle = -45, hjust = 1)

# Display the plot
C4_surface
```

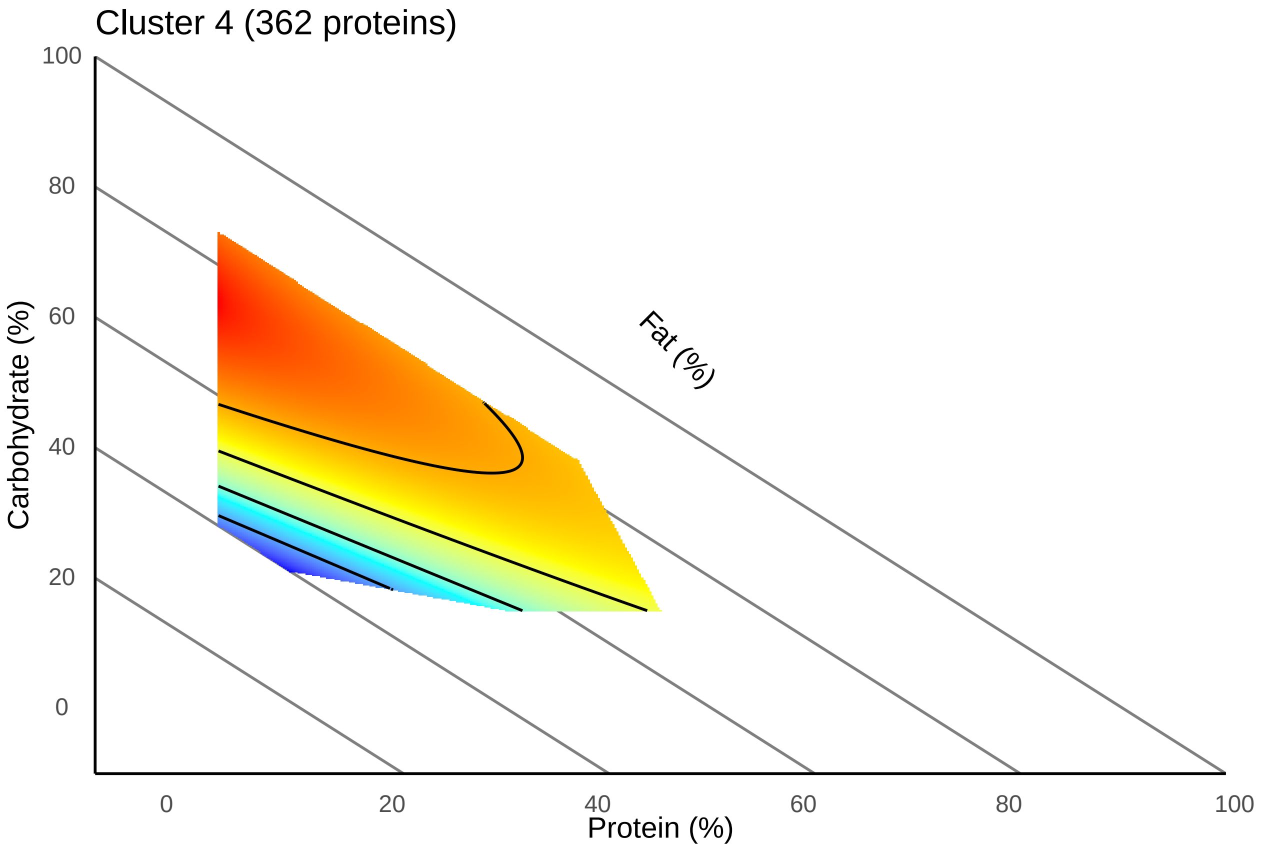

###### 4.1.4.1 Average surface

```r
valid_ids <- keys(org.Mm.eg.db, keytype = "ENTREZID")

protein_list <- Cluster4$Protein
background <- protein_tibble_fathers$proteins

# Convert Uniprot IDs to Entrez IDs
entrez_ids_c4 <- bitr(protein_list, fromType = "SYMBOL", toType = "ENTREZID", OrgDb = "org.Mm.eg.db")

# Extract the Entrez IDs
entrez_list <- entrez_ids_c4$ENTREZID

# removing missing values
entrez_list <- entrez_list[!is.na(entrez_list) & entrez_list != ""]
entrez_list <- entrez_list[entrez_list %in% valid_ids]

entrez_ids_background <- bitr(background, fromType = "SYMBOL", toType = "ENTREZID",
    OrgDb = "org.Mm.eg.db")

# Extract the Entrez IDs
entrez_list_background <- entrez_ids_background$ENTREZID

# removing missing values
entrez_list_background <- entrez_list_background[!is.na(entrez_list_background) &
    entrez_list_background != ""]
entrez_list_background <- entrez_list_background[entrez_list_background %in% valid_ids]

# GO analysis go_cluster4_all <- enrichGO(gene = entrez_list, OrgDb =
# org.Mm.eg.db, keyType = 'ENTREZID', ont = 'all', universe =
# entrez_list_background, pAdjustMethod = 'BH', pvalueCutoff = 0.05)
# go_cluster4_all_simplified <- simplify(go_cluster4_all, cutoff = 0.7, by =
# 'p.adjust', select_fun = min, measure = 'Wang')

go_cluster4_BP <- enrichGO(gene = entrez_list, OrgDb = org.Mm.eg.db, keyType = "ENTREZID",
    ont = "BP", universe = entrez_list_background, pAdjustMethod = "BH", pvalueCutoff = 0.05)

go_cluster4_BP_simplified <- clusterProfiler::simplify(go_cluster4_BP, cutoff = 0.7,
    by = "p.adjust", select_fun = min, measure = "Wang")

# go_cluster4_CC <- enrichGO(gene = entrez_list, OrgDb = org.Mm.eg.db, keyType
# = 'ENTREZID', ont = 'CC', universe = entrez_list_background, pAdjustMethod =
# 'BH', pvalueCutoff = 0.05) go_cluster4_CC_simplified <-
# simplify(go_cluster4_CC, cutoff = 0.7, by = 'p.adjust', select_fun = min,
# measure = 'Wang') go_cluster4_MF <- enrichGO(gene = entrez_list, OrgDb =
# org.Mm.eg.db, keyType = 'ENTREZID', ont = 'MF', universe =
# entrez_list_background, pAdjustMethod = 'BH', pvalueCutoff = 0.05)
# go_cluster4_MF_simplified <- simplify(go_cluster4_MF, cutoff = 0.7, by =
# 'p.adjust', select_fun = min, measure = 'Wang')

# Finding Ucp1 = entrezID = 22227
```

```r
cluster4_mapping <- as.data.frame(go_cluster4_BP_simplified@result)
Ucp1 <- cluster4_mapping[grep(22227, cluster4_mapping$geneID), ]
# not in simplified GO
```

```r
# GO_cluster4 <-clusterProfiler::dotplot(merge_result(list( 'Biological
# Process' = go_cluster4_BP_simplified,'Cellular Component' =
# go_cluster4_CC_simplified, 'Molecular Function' =
# go_cluster4_MF_simplified)), showCategory = 5) + xlab(NULL) # GO_cluster4 <-
# clusterProfiler::dotplot(go_cluster4_all_simplified, showCategory =5)
# GO_cluster4

bar_data_c4 <- go_cluster4_BP_simplified@result
top_bar_data_c4 <- bar_data_c4[order(bar_data_c4$p.adjust), ][1:3, ]
top_bar_data_c4$log10_p_adjust <- -log10(top_bar_data_c4$p.adjust)
top_bar_data_c4$percent_proteins <- ((top_bar_data_c4$Count/322) * 100)

# enrichedgenes_cluster4 <-
# unique(unlist(strsplit(go_cluster4_BP_simplified$geneID, '/')))
# length(enrichedgenes_cluster4)

C4_bar <- ggplot(top_bar_data_c4, aes(x = log10_p_adjust, y = reorder(Description,
    log10_p_adjust), fill = percent_proteins)) + geom_bar(stat = "identity") + xlim(0,
    10) + scale_fill_gradientn(colors = c("lightblue", "blue", "darkblue"), limits = c(0,
    50), breaks = seq(0, 60, by = 20)) + labs(x = NULL, y = NULL, fill = NULL) +
    theme_minimal() + ggtitle("Cluster 4 (40 proteins)") + theme(axis.text.y = element_text(size = 12),
    axis.text.x = element_text(size = 12), legend.position = "none")
C4_bar
```

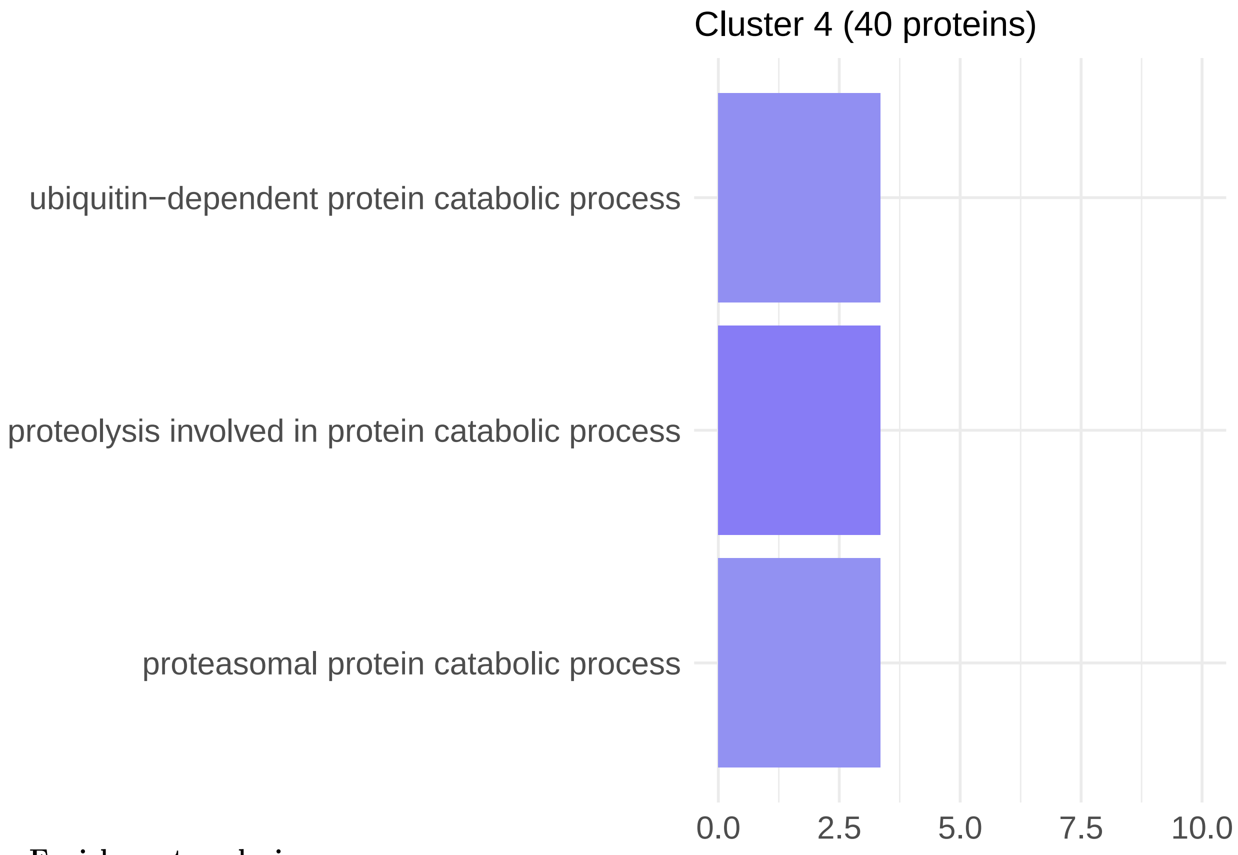

###### 4.1.4.2 Enrichment analysis

##### 4.1.5 Cluster 5

```
# need this to create surface
data = metadata
p_P = "Percent_prot"
p_C = "Percent_carb"
p_F = "Percent_fat"

coeffs = C5_descriptiveStats$Mean

mixture.surface <- function(data, coeffs, p_P = "p_P", p_C = "p_C", p_F = "p_F") {
}

# Set the resolution of the surface
surface.resolution <- 501

# How many values to round surface
round.surf <- 3

# This specifies the color scheme for surface - it is actually a function that
# returns a function
rgb.palette <- colorRampPalette(c("blue", "cyan", "yellow", "red"), space = "Lab",
    interpolate = "linear")
```

```r
# How many different colours should we use on the plot
no.cols <- 256

# Get the colors to use from the palette specified above
map <- rgb.palette(no.cols)

# How many levels should there be on the surface
nlev <- 5

# Labels for each
labels <- c("Protein (%)", "Carbohydrate (%)", "Fat (%)")

## plot the RMT surface
iso.lines <- seq(1, 0, -0.2)

# Set the layout
par(mfrow = c(1, 1), mar = c(5, 5, 5, 1))

# Make sure the proportions are closed off to 1
total <- (data[, p_P] + data[, p_C] + data[, p_F])
data$p_P <- data[, p_P]/total
data$p_C <- data[, p_C]/total
data$p_F <- data[, p_F]/total

## estimate convex hull and predict
mdff2 <- findConvex.prop(data$p_P, data$p_C, c("p_P", "p_C"), surface.resolution)
mdff2$p_F <- with(mdff2, 1 - p_P - p_C)

# Get the predicted surface
X <- model.matrix(~0 + p_P + p_C + p_F + p_P:p_C + p_P:p_F + p_C:p_F, data = mdff2)
Y <- (X %*% coeffs)
plot_data <- as.data.frame(X)
plot_data$Y <- Y
plot_data$p_P <- plot_data$p_P * 100
plot_data$p_C <- plot_data$p_C * 100
plot_data$p_F <- plot_data$p_F * 100
contour_use <- signif((max(Y) - min(Y))/5, 1)
```

```r
  C5_surface <- ggplot(plot_data) +
  # Add diagonal lines with solid lines, ensuring they cover the whole plot area
  geom_abline(intercept = 100, slope = -1, color = "grey50") +
  geom_abline(intercept = 80, slope = -1, color = "grey50") +
  geom_abline(intercept = 60, slope = -1, color = "grey50") +
  geom_abline(intercept = 40, slope = -1, color = "grey50") +
  geom_abline(intercept = 20, slope = -1, color = "grey50") +

  # Add other plot layers on top of the lines
  geom_tile(aes(x = p_P, y = p_C, fill = Y)) +
  scale_fill_gradientn(colors = map) +
```

```
  geom_contour(data = plot_data, aes(x = p_P, y = p_C, z = Y), na.rm = TRUE, color = "black", binwidth =
  # geom_label_contour(data = plot_data, aes(x = p_P, y = p_C, z = Y), size = 3, binwidth = contour_use
  #
  # Set theme and labels, remove grid lines and specific axis lines
  theme_minimal() +
  theme(
    panel.grid = element_blank(),       # Remove grid lines
    panel.border = element_blank(),     # Remove the border around the plot panel
    axis.line = element_line(color = "black"),       # Add back the x and y axis lines
    legend.position = "none",             # Remove legend
    axis.text.x = element_text(hjust = -1, vjust = -0.5),  # Adjust horizontal and vertical position fo
    axis.text.y = element_text(hjust = 0.5, vjust = -1.5)    # Adjust horizontal and vertical position
  ) +
  scale_x_continuous(limits = c(0, 100), breaks = seq(0, 100, by = 20)) +  # Set x-axis limits and labe
  scale_y_continuous(limits = c(0, 100), breaks = seq(0, 100, by = 20)) +  # Set y-axis limits and labe
  # coord_fixed(ratio = 1) +   # Fix the aspect ratio to ensure alignment of diagonals and axes
  ggtitle("Cluster 5 (260 proteins)") +
  labs(x = "Protein (%)", y = "Carbohydrate (%)") +

  # Adjust the "Fat" label position
  annotate("text", x = 55, y = 55, label = "Fat (%)", color = "black", angle = -45, hjust = 1)

# Display the plot
C5_surface
```

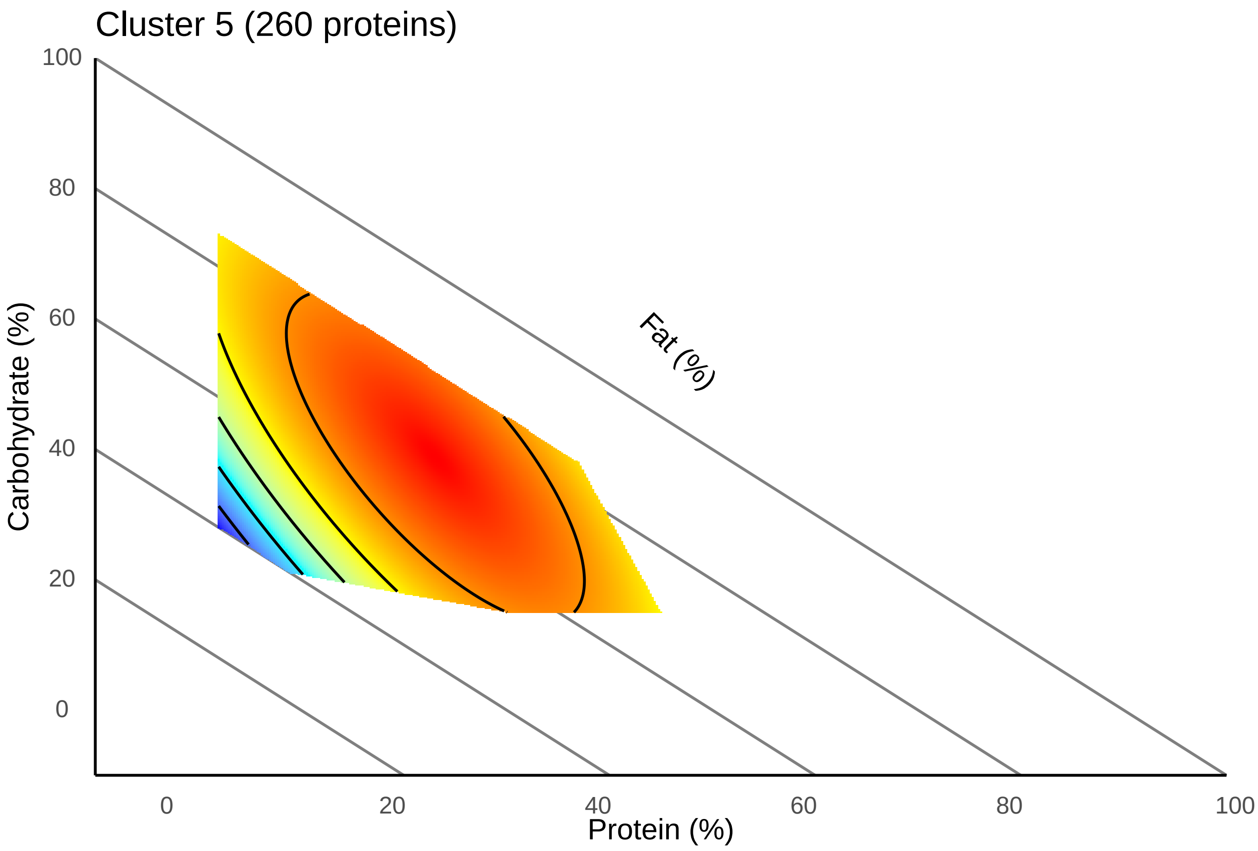

###### 4.1.5.1 Average surface

```r
valid_ids <- keys(org.Mm.eg.db, keytype = "ENTREZID")

protein_list <- Cluster5$Protein
background <- protein_tibble_fathers$proteins

# Convert Uniprot IDs to Entrez IDs
entrez_ids_c5 <- bitr(protein_list, fromType = "SYMBOL", toType = "ENTREZID", OrgDb = "org.Mm.eg.db")

# Extract the Entrez IDs
entrez_list <- entrez_ids_c5$ENTREZID

# removing missing values
entrez_list <- entrez_list[!is.na(entrez_list) & entrez_list != ""]
entrez_list <- entrez_list[entrez_list %in% valid_ids]

entrez_ids_background <- bitr(background, fromType = "SYMBOL", toType = "ENTREZID",
    OrgDb = "org.Mm.eg.db")

# Extract the Entrez IDs
entrez_list_background <- entrez_ids_background$ENTREZID

# removing missing values
entrez_list_background <- entrez_list_background[!is.na(entrez_list_background) &
    entrez_list_background != ""]
entrez_list_background <- entrez_list_background[entrez_list_background %in% valid_ids]

# GO analysis
go_cluster5_BP <- enrichGO(gene = entrez_list, OrgDb = org.Mm.eg.db, keyType = "ENTREZID",
    ont = "BP", universe = entrez_list_background, pAdjustMethod = "BH", pvalueCutoff = 0.05)
#
go_cluster5_BP_simplified <- clusterProfiler::simplify(go_cluster5_BP, cutoff = 0.7,
    by = "p.adjust", select_fun = min, measure = "Wang")

# go_cluster5_CC <- enrichGO(gene = entrez_list, OrgDb = org.Mm.eg.db, keyType
# = 'ENTREZID', ont = 'CC', universe = entrez_list_background, pAdjustMethod =
# 'BH', pvalueCutoff = 0.05) # go_cluster5_CC_simplified <-
# simplify(go_cluster5_CC, cutoff = 0.7, by = 'p.adjust', select_fun = min,
# measure = 'Wang') go_cluster5_MF <- enrichGO(gene = entrez_list, OrgDb =
# org.Mm.eg.db, keyType = 'ENTREZID', ont = 'MF', universe =
# entrez_list_background, pAdjustMethod = 'BH', pvalueCutoff = 0.05) #
# go_cluster5_MF_simplified <- simplify(go_cluster5_MF, cutoff = 0.7, by =
# 'p.adjust', select_fun = min, measure = 'Wang')
```

```r
# GO_cluster5 <-clusterProfiler::dotplot(merge_result(list( 'Biological
# Process' = go_cluster5_BP_simplified,'Cellular Component' =
# go_cluster5_CC_simplified, 'Molecular Function' =
# go_cluster5_MF_simplified)), showCategory = 5) + xlab(NULL) # GO_cluster5 <-
# clusterProfiler::dotplot(go_cluster5_all_simplified, showCategory =5)
# GO_cluster5
```

```r
bar_data_c5 <- go_cluster5_BP_simplified@result
top_bar_data_c5 <- bar_data_c5[order(bar_data_c5$p.adjust), ][1:5, ]
top_bar_data_c5$log10_p_adjust <- -log10(top_bar_data_c5$p.adjust)
top_bar_data_c5$percent_proteins <- ((top_bar_data_c5$Count/256) * 100)

# top_bar_data <- bar_data[order(bar_data$log10_p_adjust), ][1:10]

# enrichedgenes_cluster5 <-
# unique(unlist(strsplit(go_cluster5_BP_simplified$geneID, '/')))
# length(enrichedgenes_cluster5)

C5_bar <- ggplot(top_bar_data_c5, aes(x = log10_p_adjust, y = reorder(Description,
    log10_p_adjust), fill = percent_proteins)) + geom_bar(stat = "identity") + xlim(0,
    10) + scale_fill_gradientn(colors = c("lightblue", "blue", "darkblue"), limits = c(0,
    50), breaks = seq(0, 60, by = 20)) + labs(x = "-log(10) p-adjust", y = NULL,
    fill = NULL) + theme_minimal() + ggtitle("Cluster 5 (188 proteins)") + theme(axis.text.y = element_
    axis.text.x = element_text(size = 12), legend.position = "none")
C5_bar
```

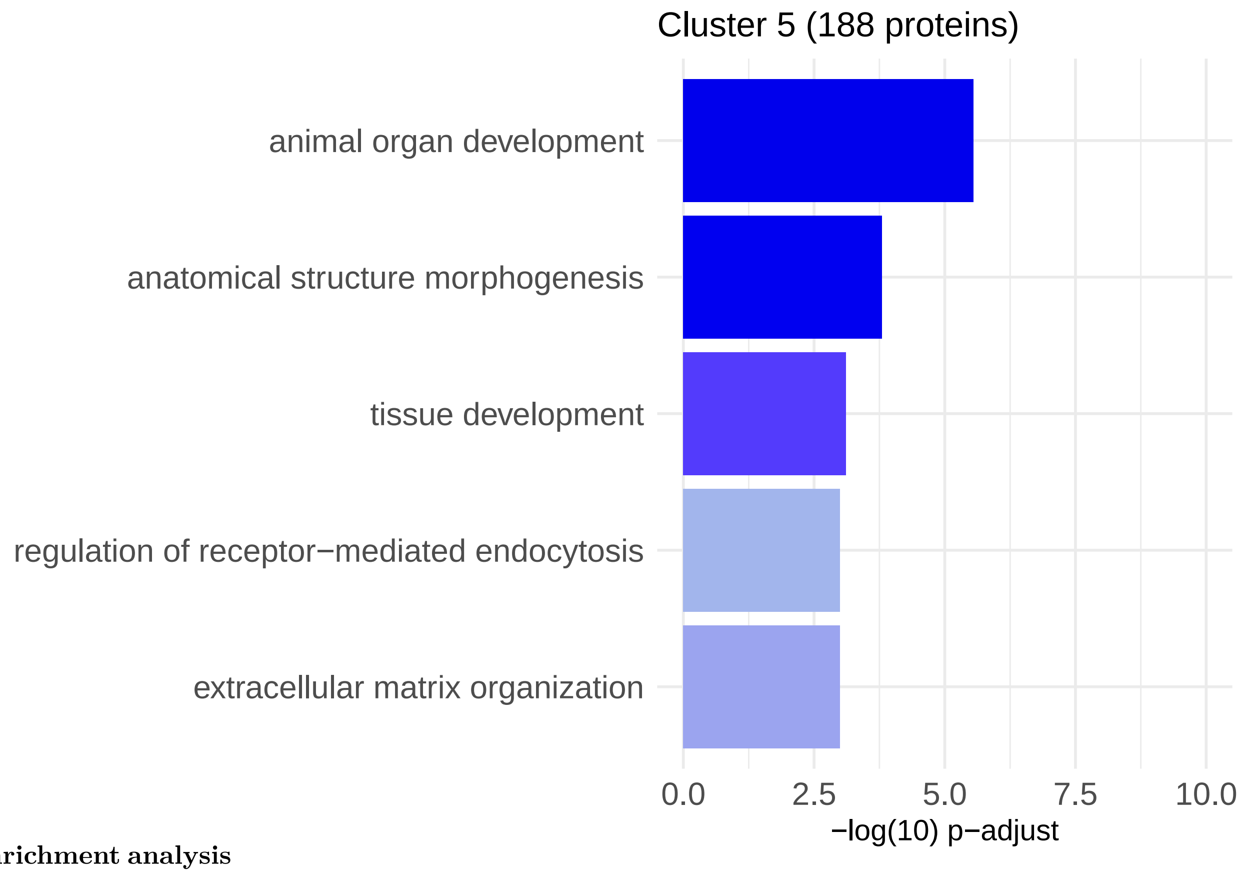

###### 4.1.5.2 Enrichment analysis

#### 4.2 Individual surfaces of 'hub' proteins

##### 4.2.1 Cluster 1

**4.2.1.1 CYC1** This protein is no longer in our significant 'set' of proteins when including dietary intake

```r
Cyc1 <- subset(Cluster1_imported, Protein == "Cyc1")
Cyc1_joined <- tidy_protein_table_fathers[tidy_protein_table_fathers$proteins %in%
    Cyc1, ]

Cyc1_joined$pP <- (Cyc1_joined$Percent_prot/100)
Cyc1_joined$pC <- (Cyc1_joined$Percent_carb/100)
Cyc1_joined$pF <- (Cyc1_joined$Percent_fat/100)

Cyc1_joined$LogIntensity_z <- scale(Cyc1_joined$LogIntensity)
model <- lm(LogIntensity_z ~ 0 + pP + pC + pF + pP:pC + pP:pF + pC:pF, data = Cyc1_joined)

coeffs <- coef(model)
data = metadata
p_P = "Percent_prot"
p_C = "Percent_carb"
p_F = "Percent_fat"

mixture.surface <- function(data, coeffs, p_P = "p_P", p_C = "p_C", p_F = "p_F") {
}

# Set the resolution of the surface
surface.resolution <- 501

# How many values to round surface
round.surf <- 3

# This specifies the color scheme for surface - it is actually a function that
# returns a function
rgb.palette <- colorRampPalette(c("blue", "cyan", "yellow", "red"), space = "Lab",
    interpolate = "linear")

# How many different colours should we use on the plot
no.cols <- 256

# Get the colors to use from the palette specified above
map <- rgb.palette(no.cols)

# How many levels should there be on the surface
nlev <- 5

# Labels for each
labels <- c("Protein (%)", "Carbohydrate (%)", "Fat (%)")

## plot the RMT surface
iso.lines <- seq(1, 0, -0.2)

# Set the layout
par(mfrow = c(1, 1), mar = c(5, 5, 5, 1))

# Make sure the proportions are closed off to 1
total <- (data[, p_P] + data[, p_C] + data[, p_F])
```

```
data$p_P <- data[, p_P]/total
data$p_C <- data[, p_C]/total
data$p_F <- data[, p_F]/total

## estimate convex hull and predict
mdff2 <- findConvex.prop(data$p_P, data$p_C, c("p_P", "p_C"), surface.resolution)
mdff2$p_F <- with(mdff2, 1 - p_P - p_C)

# Get the predicted surface
X <- model.matrix(~0 + p_P + p_C + p_F + p_P:p_C + p_P:p_F + p_C:p_F, data = mdff2)
Y <- (X %*% coeffs)
plot_data <- as.data.frame(X)
plot_data$Y <- Y
plot_data$p_P <- plot_data$p_P * 100
plot_data$p_C <- plot_data$p_C * 100
plot_data$p_F <- plot_data$p_F * 100
contour_use <- signif((max(Y) - min(Y))/5, 1)
```

```
  Cyc1_fathers <- ggplot(plot_data) +
  # Add diagonal lines with solid lines, ensuring they cover the whole plot area
  geom_abline(intercept = 100, slope = -1, color = "grey50") +
  geom_abline(intercept = 80, slope = -1, color = "grey50") +
  geom_abline(intercept = 60, slope = -1, color = "grey50") +
  geom_abline(intercept = 40, slope = -1, color = "grey50") +
  geom_abline(intercept = 20, slope = -1, color = "grey50") +

  # Add other plot layers on top of the lines
  geom_tile(aes(x = p_P, y = p_C, fill = Y)) +
  scale_fill_gradientn(colors = map) +
  geom_contour(data = plot_data, aes(x = p_P, y = p_C, z = Y), na.rm = TRUE, color = "black", binwidth =
  geom_label_contour(data = plot_data, aes(x = p_P, y = p_C, z = Y), size = 3, binwidth = contour_use) +

  # Set theme and labels, remove grid lines and specific axis lines
  theme_minimal() +
  theme(
    panel.grid = element_blank(),       # Remove grid lines
    panel.border = element_blank(),     # Remove the border around the plot panel
    axis.line = element_line(color = "black"),       # Add back the x and y axis lines
    legend.position = "none",             # Remove legend
    axis.text.x = element_text(hjust = -1, vjust = -0.5),  # Adjust horizontal and vertical position fo
    axis.text.y = element_text(hjust = 0.5, vjust = -1.5)    # Adjust horizontal and vertical position
  ) +
  scale_x_continuous(limits = c(0, 100), breaks = seq(0, 100, by = 20)) +  # Set x-axis limits and labe
  scale_y_continuous(limits = c(0, 100), breaks = seq(0, 100, by = 20)) +  # Set y-axis limits and labe
  coord_fixed(ratio = 1) +   # Fix the aspect ratio to ensure alignment of diagonals and axes
  ggtitle("CYC1 (cluster 1)") +
  labs(x = "Protein (%)", y = "Carbohydrate (%)") +

  # Adjust the "Fat" label position
  annotate("text", x = 55, y = 55, label = "Fat (%)", color = "black", angle = -45, hjust = 1)

# Display the plot
Cyc1_fathers
```

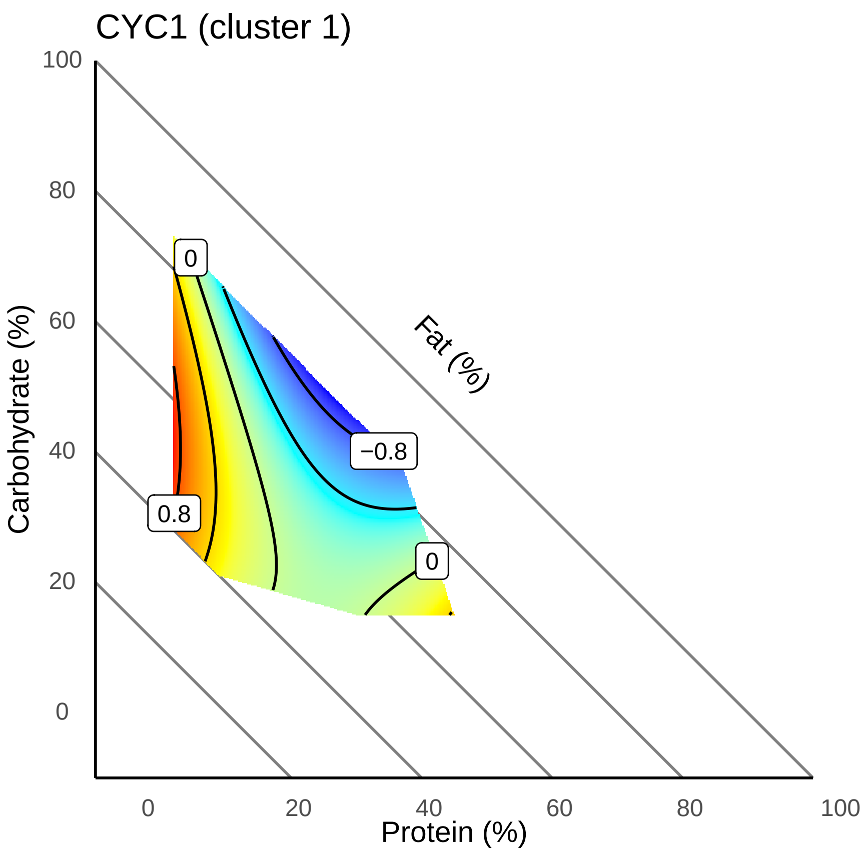

**4.2.1.2 UQCRC1** This protein is no longer in our significant 'set' of proteins when including dietary intake

```
Uqcrc1 <- subset(Cluster1_imported, Protein == "Uqcrc1")
Uqcrc1_joined <- tidy_protein_table_fathers[tidy_protein_table_fathers$proteins %in%
    Uqcrc1, ]

Uqcrc1_joined$pP <- (Uqcrc1_joined$Percent_prot/100)
Uqcrc1_joined$pC <- (Uqcrc1_joined$Percent_carb/100)
Uqcrc1_joined$pF <- (Uqcrc1_joined$Percent_fat/100)

Uqcrc1_joined$LogIntensity_z <- scale(Uqcrc1_joined$LogIntensity)
model <- lm(LogIntensity_z ~ 0 + pP + pC + pF + pP:pC + pP:pF + pC:pF, data = Uqcrc1_joined)

coeffs <- coef(model)
data = metadata
p_P = "Percent_prot"
p_C = "Percent_carb"
p_F = "Percent_fat"

mixture.surface <- function(data, coeffs, p_P = "p_P", p_C = "p_C", p_F = "p_F") {
}
```

```r
# Set the resolution of the surface
surface.resolution <- 501

# How many values to round surface
round.surf <- 3

# This specifies the color scheme for surface - it is actually a function that
# returns a function
rgb.palette <- colorRampPalette(c("blue", "cyan", "yellow", "red"), space = "Lab",
    interpolate = "linear")

# How many different colours should we use on the plot
no.cols <- 256

# Get the colors to use from the palette specified above
map <- rgb.palette(no.cols)

# How many levels should there be on the surface
nlev <- 5

# Labels for each
labels <- c("Protein (%)", "Carbohydrate (%)", "Fat (%)")

## plot the RMT surface
iso.lines <- seq(1, 0, -0.2)

# Set the layout
par(mfrow = c(1, 1), mar = c(5, 5, 5, 1))

# Make sure the proportions are closed off to 1
total <- (data[, p_P] + data[, p_C] + data[, p_F])
data$p_P <- data[, p_P]/total
data$p_C <- data[, p_C]/total
data$p_F <- data[, p_F]/total

## estimate convex hull and predict
mdff2 <- findConvex.prop(data$p_P, data$p_C, c("p_P", "p_C"), surface.resolution)
mdff2$p_F <- with(mdff2, 1 - p_P - p_C)

# Get the predicted surface
X <- model.matrix(~0 + p_P + p_C + p_F + p_P:p_C + p_P:p_F + p_C:p_F, data = mdff2)
Y <- (X %*% coeffs)
plot_data <- as.data.frame(X)
plot_data$Y <- Y
plot_data$p_P <- plot_data$p_P * 100
plot_data$p_C <- plot_data$p_C * 100
plot_data$p_F <- plot_data$p_F * 100
contour_use <- signif((max(Y) - min(Y))/5, 1)
```

```r
  Uqcrc1_fathers <- ggplot(plot_data) +
  # Add diagonal lines with solid lines, ensuring they cover the whole plot area
  geom_abline(intercept = 100, slope = -1, color = "grey50") +
  geom_abline(intercept = 80, slope = -1, color = "grey50") +
```

```r
  geom_abline(intercept = 60, slope = -1, color = "grey50") +
  geom_abline(intercept = 40, slope = -1, color = "grey50") +
  geom_abline(intercept = 20, slope = -1, color = "grey50") +

  # Add other plot layers on top of the lines
  geom_tile(aes(x = p_P, y = p_C, fill = Y)) +
  scale_fill_gradientn(colors = map) +
  geom_contour(data = plot_data, aes(x = p_P, y = p_C, z = Y), na.rm = TRUE, color = "black", binwidth =
  geom_label_contour(data = plot_data, aes(x = p_P, y = p_C, z = Y), size = 3, binwidth = contour_use) +

  # Set theme and labels, remove grid lines and specific axis lines
  theme_minimal() +
  theme(
    panel.grid = element_blank(),        # Remove grid lines
    panel.border = element_blank(),      # Remove the border around the plot panel
    axis.line = element_line(color = "black"),       # Add back the x and y axis lines
    legend.position = "none",             # Remove legend
    axis.text.x = element_text(hjust = -1, vjust = -0.5),  # Adjust horizontal and vertical position fo
    axis.text.y = element_text(hjust = 0.5, vjust = -1.5)    # Adjust horizontal and vertical position
  ) +
  scale_x_continuous(limits = c(0, 100), breaks = seq(0, 100, by = 20)) +  # Set x-axis limits and labe
  scale_y_continuous(limits = c(0, 100), breaks = seq(0, 100, by = 20)) +  # Set y-axis limits and labe
  coord_fixed(ratio = 1) +   # Fix the aspect ratio to ensure alignment of diagonals and axes
  ggtitle("UQCRC1 (cluster 1)") +
  labs(x = "Protein (%)", y = "Carbohydrate (%)") +

  # Adjust the "Fat" label position
  annotate("text", x = 55, y = 55, label = "Fat (%)", color = "black", angle = -45, hjust = 1)

# Display the plot
Uqcrc1_fathers
```

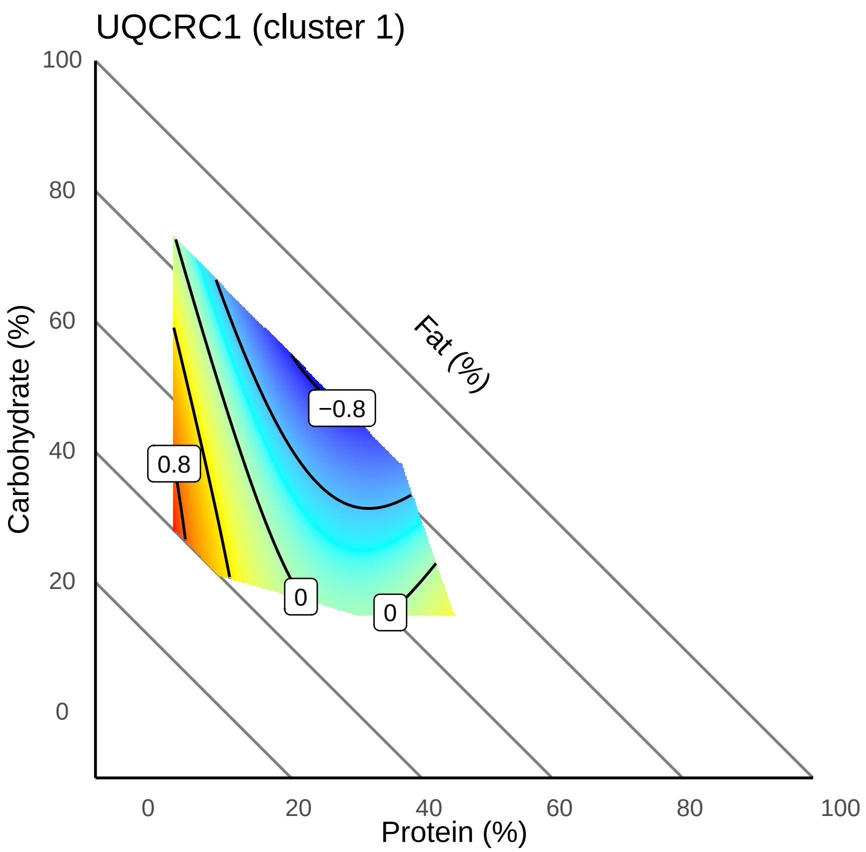

```
p <- (Cyc1_fathers | Uqcrc1_fathers)
p
```

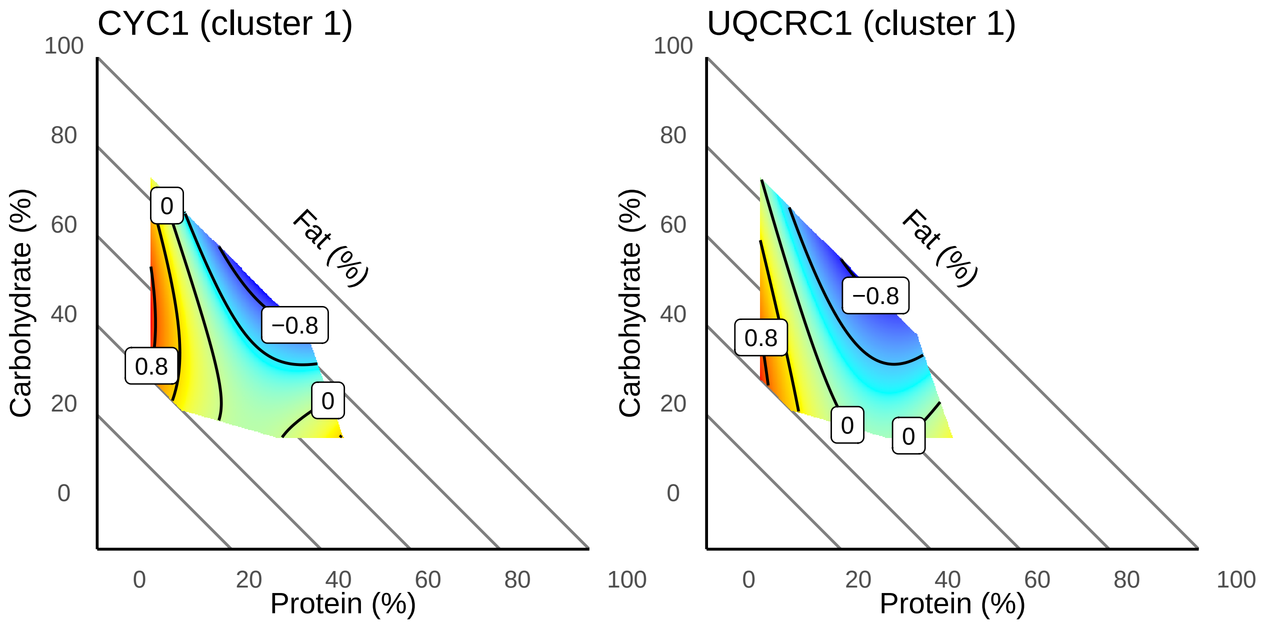

###### 4.2.1.3 Combined (Fig. S3)

##### 4.2.2 Cluster 2

###### 4.2.2.1 GLUT4 (Fig. S4)

Gene name is Slc2a4

```
GLUT4 <- subset(Cluster2, Protein == "Slc2a4")

GLUT4_joined <- tidy_protein_table_fathers[tidy_protein_table_fathers$proteins %in%
    GLUT4, ]

GLUT4_joined$pP <- (GLUT4_joined$Percent_prot/100)
GLUT4_joined$pC <- (GLUT4_joined$Percent_carb/100)
GLUT4_joined$pF <- (GLUT4_joined$Percent_fat/100)

GLUT4_joined$LogIntensity_z <- scale(GLUT4_joined$LogIntensity)
model <- lm(LogIntensity_z ~ 0 + pP + pC + pF + pP:pC + pP:pF + pC:pF, data = GLUT4_joined)

coeffs <- coef(model)
data = metadata
p_P = "Percent_prot"
p_C = "Percent_carb"
p_F = "Percent_fat"

mixture.surface <- function(data, coeffs, p_P = "p_P", p_C = "p_C", p_F = "p_F") {
```

```
}

# Set the resolution of the surface
surface.resolution <- 501

# How many values to round surface
round.surf <- 3

# This specifies the color scheme for surface - it is actually a function that
# returns a function
rgb.palette <- colorRampPalette(c("blue", "cyan", "yellow", "red"), space = "Lab",
    interpolate = "linear")

# How many different colours should we use on the plot
no.cols <- 256

# Get the colors to use from the palette specified above
map <- rgb.palette(no.cols)

# How many levels should there be on the surface
nlev <- 5

# Labels for each
labels <- c("Protein (%)", "Carbohydrate (%)", "Fat (%)")

## plot the RMT surface
iso.lines <- seq(1, 0, -0.2)

# Set the layout
par(mfrow = c(1, 1), mar = c(5, 5, 5, 1))

# Make sure the proportions are closed off to 1
total <- (data[, p_P] + data[, p_C] + data[, p_F])
data$p_P <- data[, p_P]/total
data$p_C <- data[, p_C]/total
data$p_F <- data[, p_F]/total

## estimate convex hull and predict
mdff2 <- findConvex.prop(data$p_P, data$p_C, c("p_P", "p_C"), surface.resolution)
mdff2$p_F <- with(mdff2, 1 - p_P - p_C)

# Get the predicted surface
X <- model.matrix(~0 + p_P + p_C + p_F + p_P:p_C + p_P:p_F + p_C:p_F, data = mdff2)
Y <- (X %*% coeffs)
plot_data <- as.data.frame(X)
plot_data$Y <- Y
plot_data$p_P <- plot_data$p_P * 100
plot_data$p_C <- plot_data$p_C * 100
plot_data$p_F <- plot_data$p_F * 100
contour_use <- signif((max(Y) - min(Y))/5, 1)
```

```
  Glut4_fathers <- ggplot(plot_data) +
  # Add diagonal lines with solid lines, ensuring they cover the whole plot area
```

```
  geom_abline(intercept = 100, slope = -1, color = "grey50") +
  geom_abline(intercept = 80, slope = -1, color = "grey50") +
  geom_abline(intercept = 60, slope = -1, color = "grey50") +
  geom_abline(intercept = 40, slope = -1, color = "grey50") +
  geom_abline(intercept = 20, slope = -1, color = "grey50") +

  # Add other plot layers on top of the lines
  geom_tile(aes(x = p_P, y = p_C, fill = Y)) +
  scale_fill_gradientn(colors = map) +
  geom_contour(data = plot_data, aes(x = p_P, y = p_C, z = Y), na.rm = TRUE, color = "black", binwidth =
  geom_label_contour(data = plot_data, aes(x = p_P, y = p_C, z = Y), size = 3, binwidth = contour_use) +

  # Set theme and labels, remove grid lines and specific axis lines
  theme_minimal() +
  theme(
    panel.grid = element_blank(),       # Remove grid lines
    panel.border = element_blank(),     # Remove the border around the plot panel
    axis.line = element_line(color = "black"),       # Add back the x and y axis lines
    legend.position = "none",             # Remove legend
    axis.text.x = element_text(hjust = -1, vjust = -0.5),  # Adjust horizontal and vertical position fo
    axis.text.y = element_text(hjust = 0.5, vjust = -1.5)    # Adjust horizontal and vertical position
  ) +
  scale_x_continuous(limits = c(0, 100), breaks = seq(0, 100, by = 20)) +  # Set x-axis limits and labe
  scale_y_continuous(limits = c(0, 100), breaks = seq(0, 100, by = 20)) +  # Set y-axis limits and labe
  coord_fixed(ratio = 1) +   # Fix the aspect ratio to ensure alignment of diagonals and axes
  ggtitle("GLUT4 (cluster 2)") +
  labs(x = "Protein (%)", y = "Carbohydrate (%)") +

  # Adjust the "Fat" label position
  annotate("text", x = 55, y = 55, label = "Fat (%)", color = "black", angle = -45, hjust = 1)

# Display the plot
Glut4_fathers
```

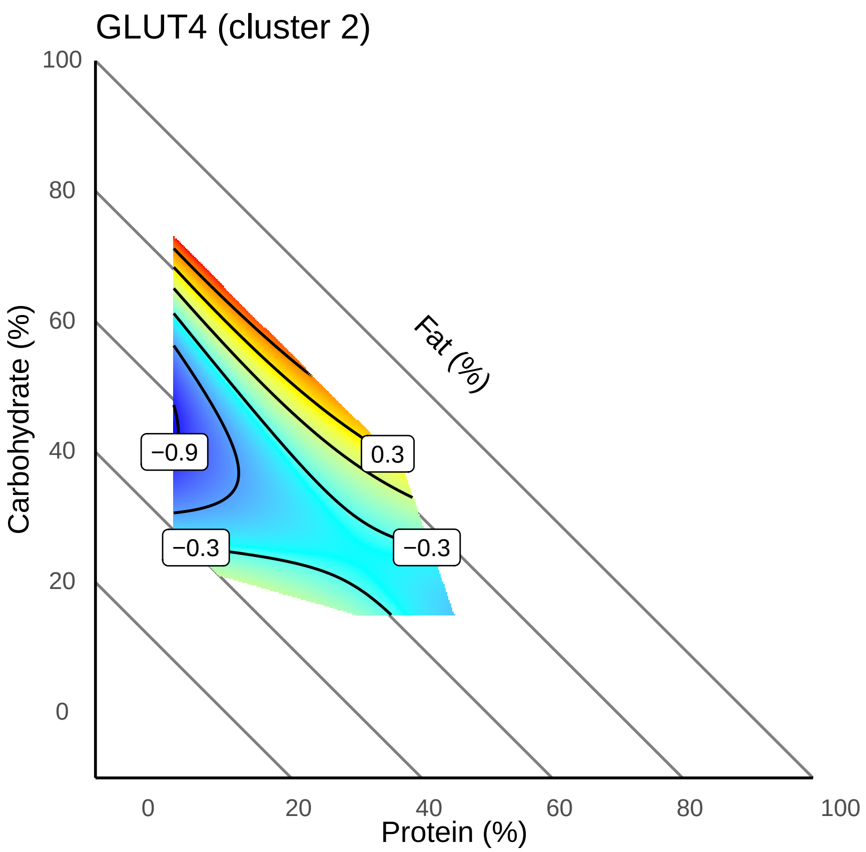

###### 4.2.3 Cluster 3

```
Cs <- subset(Cluster3, Protein == "Cs")
Cs_joined <- tidy_protein_table_fathers[tidy_protein_table_fathers$proteins %in%
    Cs, ]

Cs_joined$pP <- (Cs_joined$Percent_prot/100)
Cs_joined$pC <- (Cs_joined$Percent_carb/100)
Cs_joined$pF <- (Cs_joined$Percent_fat/100)

Cs_joined$LogIntensity_z <- scale(Cs_joined$LogIntensity)
model <- lm(LogIntensity_z ~ 0 + pP + pC + pF + pP:pC + pP:pF + pC:pF, data = Cs_joined)

coeffs <- coef(model)
data = metadata
p_P = "Percent_prot"
p_C = "Percent_carb"
p_F = "Percent_fat"

mixture.surface <- function(data, coeffs, p_P = "p_P", p_C = "p_C", p_F = "p_F") {
}
```

```r
# Set the resolution of the surface
surface.resolution <- 501

# How many values to round surface
round.surf <- 3

# This specifies the color scheme for surface - it is actually a function that
# returns a function
rgb.palette <- colorRampPalette(c("blue", "cyan", "yellow", "red"), space = "Lab",
    interpolate = "linear")

# How many different colours should we use on the plot
no.cols <- 256

# Get the colors to use from the palette specified above
map <- rgb.palette(no.cols)

# How many levels should there be on the surface
nlev <- 5

# Labels for each
labels <- c("Protein (%)", "Carbohydrate (%)", "Fat (%)")

## plot the RMT surface
iso.lines <- seq(1, 0, -0.2)

# Set the layout
par(mfrow = c(1, 1), mar = c(5, 5, 5, 1))

# Make sure the proportions are closed off to 1
total <- (data[, p_P] + data[, p_C] + data[, p_F])
data$p_P <- data[, p_P]/total
data$p_C <- data[, p_C]/total
data$p_F <- data[, p_F]/total

## estimate convex hull and predict
mdff2 <- findConvex.prop(data$p_P, data$p_C, c("p_P", "p_C"), surface.resolution)
mdff2$p_F <- with(mdff2, 1 - p_P - p_C)

# Get the predicted surface
X <- model.matrix(~0 + p_P + p_C + p_F + p_P:p_C + p_P:p_F + p_C:p_F, data = mdff2)
Y <- (X %*% coeffs)
plot_data <- as.data.frame(X)
plot_data$Y <- Y
plot_data$p_P <- plot_data$p_P * 100
plot_data$p_C <- plot_data$p_C * 100
plot_data$p_F <- plot_data$p_F * 100
contour_use <- signif((max(Y) - min(Y))/5, 1)
```

```r
  Cs_fathers <- ggplot(plot_data) +
  # Add diagonal lines with solid lines, ensuring they cover the whole plot area
```

```
  geom_abline(intercept = 100, slope = -1, color = "grey50") +
  geom_abline(intercept = 80, slope = -1, color = "grey50") +
  geom_abline(intercept = 60, slope = -1, color = "grey50") +
  geom_abline(intercept = 40, slope = -1, color = "grey50") +
  geom_abline(intercept = 20, slope = -1, color = "grey50") +

  # Add other plot layers on top of the lines
  geom_tile(aes(x = p_P, y = p_C, fill = Y)) +
  scale_fill_gradientn(colors = map) +
  geom_contour(data = plot_data, aes(x = p_P, y = p_C, z = Y), na.rm = TRUE, color = "black", binwidth =
  geom_label_contour(data = plot_data, aes(x = p_P, y = p_C, z = Y), size = 3, binwidth = contour_use) +

  # Set theme and labels, remove grid lines and specific axis lines
  theme_minimal() +
  theme(
    panel.grid = element_blank(),       # Remove grid lines
    panel.border = element_blank(),     # Remove the border around the plot panel
    axis.line = element_line(color = "black"),       # Add back the x and y axis lines
    legend.position = "none",            # Remove legend
    axis.text.x = element_text(hjust = -1, vjust = -0.5),  # Adjust horizontal and vertical position fo
    axis.text.y = element_text(hjust = 0.5, vjust = -1.5)    # Adjust horizontal and vertical position f
  ) +
  scale_x_continuous(limits = c(0, 100), breaks = seq(0, 100, by = 20)) +  # Set x-axis limits and labe
  scale_y_continuous(limits = c(0, 100), breaks = seq(0, 100, by = 20)) +  # Set y-axis limits and labe
  coord_fixed(ratio = 1) +   # Fix the aspect ratio to ensure alignment of diagonals and axes
  ggtitle("CS (cluster 3)") +
  labs(x = "Protein (%)", y = "Carbohydrate (%)") +

  # Adjust the "Fat" label position
  annotate("text", x = 55, y = 55, label = "Fat (%)", color = "black", angle = -45, hjust = 1)

# Display the plot
Cs_fathers
```

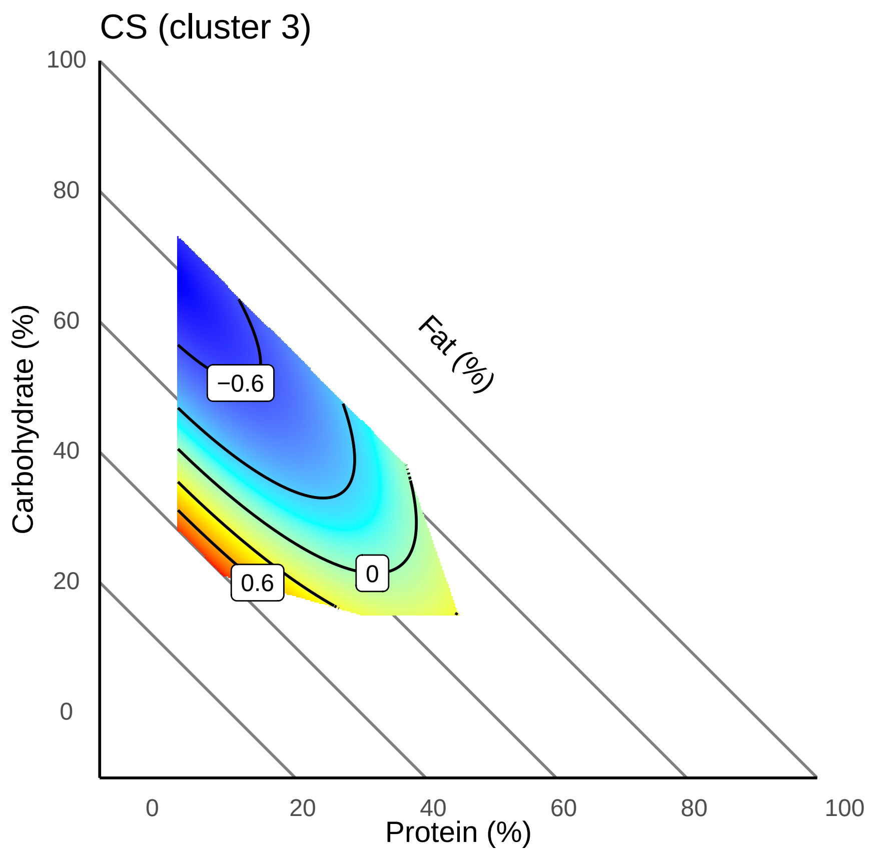

#### 4.2.3.1 CS

```
Ndufs2 <- subset(Cluster3, Protein == "Ndufs2")
Ndufs2_joined <- tidy_protein_table_fathers[tidy_protein_table_fathers$proteins %in%
    Ndufs2, ]

Ndufs2_joined$pP <- (Ndufs2_joined$Percent_prot/100)
Ndufs2_joined$pC <- (Ndufs2_joined$Percent_carb/100)
Ndufs2_joined$pF <- (Ndufs2_joined$Percent_fat/100)

Ndufs2_joined$LogIntensity_z <- scale(Ndufs2_joined$LogIntensity)
model <- lm(LogIntensity_z ~ 0 + pP + pC + pF + pP:pC + pP:pF + pC:pF, data = Ndufs2_joined)

coeffs <- coef(model)
data = metadata
p_P = "Percent_prot"
p_C = "Percent_carb"
p_F = "Percent_fat"

mixture.surface <- function(data, coeffs, p_P = "p_P", p_C = "p_C", p_F = "p_F") {
}

# Set the resolution of the surface
```

```r
surface.resolution <- 501

# How many values to round surface
round.surf <- 3

# This specifies the color scheme for surface - it is actually a function that
# returns a function
rgb.palette <- colorRampPalette(c("blue", "cyan", "yellow", "red"), space = "Lab",
    interpolate = "linear")

# How many different colours should we use on the plot
no.cols <- 256

# Get the colors to use from the palette specified above
map <- rgb.palette(no.cols)

# How many levels should there be on the surface
nlev <- 5

# Labels for each
labels <- c("Protein (%)", "Carbohydrate (%)", "Fat (%)")

## plot the RMT surface
iso.lines <- seq(1, 0, -0.2)

# Set the layout
par(mfrow = c(1, 1), mar = c(5, 5, 5, 1))

# Make sure the proportions are closed off to 1
total <- (data[, p_P] + data[, p_C] + data[, p_F])
data$p_P <- data[, p_P]/total
data$p_C <- data[, p_C]/total
data$p_F <- data[, p_F]/total

## estimate convex hull and predict
mdff2 <- findConvex.prop(data$p_P, data$p_C, c("p_P", "p_C"), surface.resolution)
mdff2$p_F <- with(mdff2, 1 - p_P - p_C)

# Get the predicted surface
X <- model.matrix(~0 + p_P + p_C + p_F + p_P:p_C + p_P:p_F + p_C:p_F, data = mdff2)
Y <- (X %*% coeffs)
plot_data <- as.data.frame(X)
plot_data$Y <- Y
plot_data$p_P <- plot_data$p_P * 100
plot_data$p_C <- plot_data$p_C * 100
plot_data$p_F <- plot_data$p_F * 100
contour_use <- signif((max(Y) - min(Y))/5, 1)
```

```r
  Ndufs2_fathers <- ggplot(plot_data) +
  # Add diagonal lines with solid lines, ensuring they cover the whole plot area
  geom_abline(intercept = 100, slope = -1, color = "grey50") +
  geom_abline(intercept = 80, slope = -1, color = "grey50") +
```

```
  geom_abline(intercept = 60, slope = -1, color = "grey50") +
  geom_abline(intercept = 40, slope = -1, color = "grey50") +
  geom_abline(intercept = 20, slope = -1, color = "grey50") +

  # Add other plot layers on top of the lines
  geom_tile(aes(x = p_P, y = p_C, fill = Y)) +
  scale_fill_gradientn(colors = map) +
  geom_contour(data = plot_data, aes(x = p_P, y = p_C, z = Y), na.rm = TRUE, color = "black", binwidth =
  geom_label_contour(data = plot_data, aes(x = p_P, y = p_C, z = Y), size = 3, binwidth = contour_use) +

  # Set theme and labels, remove grid lines and specific axis lines
  theme_minimal() +
  theme(
    panel.grid = element_blank(),       # Remove grid lines
    panel.border = element_blank(),     # Remove the border around the plot panel
    axis.line = element_line(color = "black"),       # Add back the x and y axis lines
    legend.position = "none",             # Remove legend
    axis.text.x = element_text(hjust = -1, vjust = -0.5),  # Adjust horizontal and vertical position fo
    axis.text.y = element_text(hjust = 0.5, vjust = -1.5)    # Adjust horizontal and vertical position
  ) +
  scale_x_continuous(limits = c(0, 100), breaks = seq(0, 100, by = 20)) +  # Set x-axis limits and labe
  scale_y_continuous(limits = c(0, 100), breaks = seq(0, 100, by = 20)) +  # Set y-axis limits and labe
  coord_fixed(ratio = 1) +   # Fix the aspect ratio to ensure alignment of diagonals and axes
  ggtitle("NDUFS2 (cluster 3)") +
  labs(x = "Protein (%)", y = "Carbohydrate (%)") +

  # Adjust the "Fat" label position
  annotate("text", x = 55, y = 55, label = "Fat (%)", color = "black", angle = -45, hjust = 1)

# Display the plot
Ndufs2_fathers
```

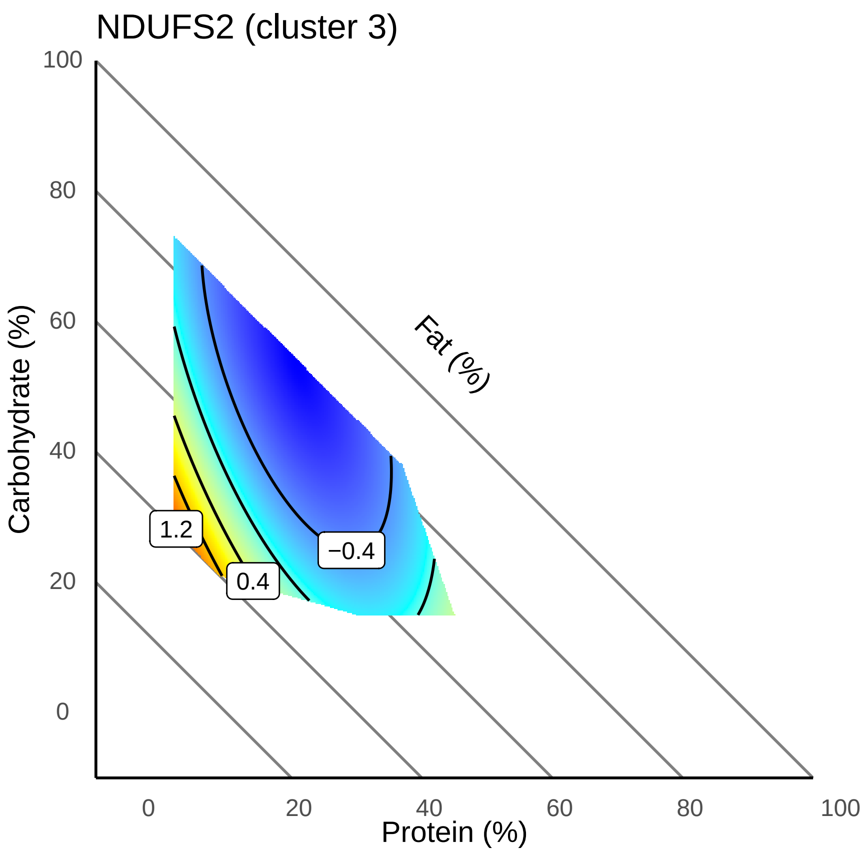

###### 4.2.3.2 NDUFS2

```
p <- (Cs_fathers | Ndufs2_fathers)
p
```

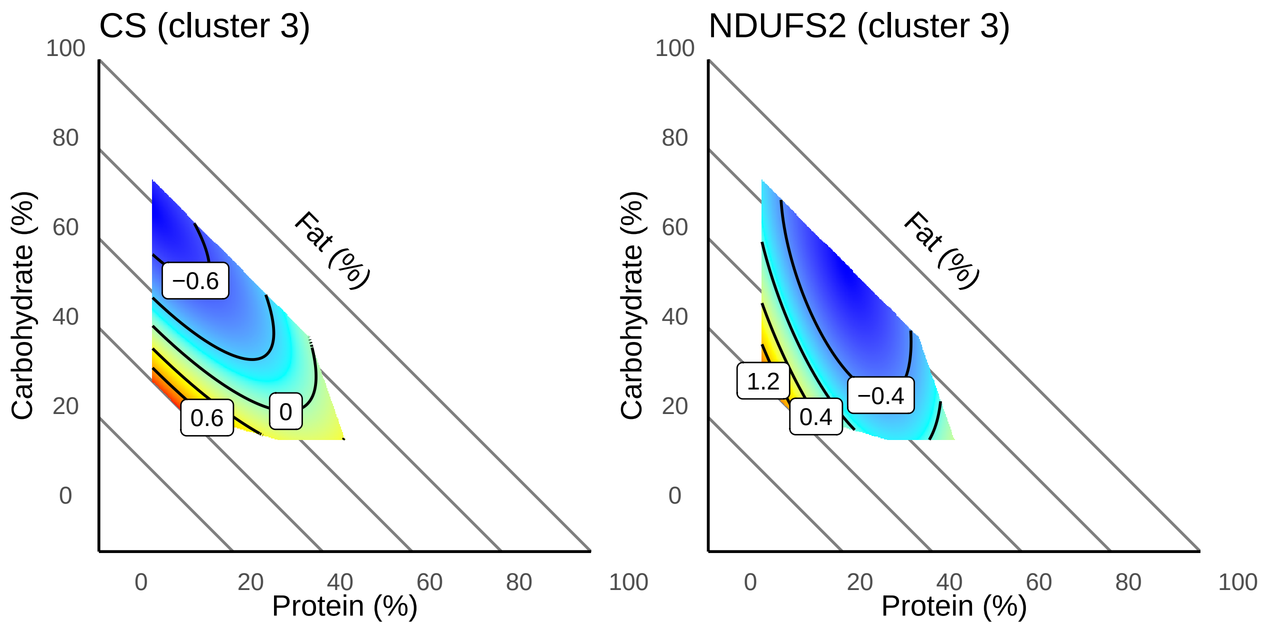

###### 4.2.3.3 Combined (Fig. S5)

##### 4.2.4 Cluster 4

These are presented in the main text

##### 4.2.5 Cluster 5

```
Apoa1 <- subset(Cluster5, Protein == "Apoa1")
Apoa1_joined <- tidy_protein_table_fathers[tidy_protein_table_fathers$proteins %in%
    Apoa1, ]

Apoa1_joined$pP <- (Apoa1_joined$Percent_prot/100)
Apoa1_joined$pC <- (Apoa1_joined$Percent_carb/100)
Apoa1_joined$pF <- (Apoa1_joined$Percent_fat/100)

Apoa1_joined$LogIntensity_z <- scale(Apoa1_joined$LogIntensity)
model <- lm(LogIntensity_z ~ 0 + pP + pC + pF + pP:pC + pP:pF + pC:pF, data = Apoa1_joined)

coeffs <- coef(model)
data = metadata
p_P = "Percent_prot"
p_C = "Percent_carb"
```

```r
p_F = "Percent_fat"

mixture.surface <- function(data, coeffs, p_P = "p_P", p_C = "p_C", p_F = "p_F") {
}

# Set the resolution of the surface
surface.resolution <- 501

# How many values to round surface
round.surf <- 3

# This specifies the color scheme for surface - it is actually a function that
# returns a function
rgb.palette <- colorRampPalette(c("blue", "cyan", "yellow", "red"), space = "Lab",
    interpolate = "linear")

# How many different colours should we use on the plot
no.cols <- 256

# Get the colors to use from the palette specified above
map <- rgb.palette(no.cols)

# How many levels should there be on the surface
nlev <- 5

# Labels for each
labels <- c("Protein (%)", "Carbohydrate (%)", "Fat (%)")

## plot the RMT surface
iso.lines <- seq(1, 0, -0.2)

# Set the layout
par(mfrow = c(1, 1), mar = c(5, 5, 5, 1))

# Make sure the proportions are closed off to 1
total <- (data[, p_P] + data[, p_C] + data[, p_F])
data$p_P <- data[, p_P]/total
data$p_C <- data[, p_C]/total
data$p_F <- data[, p_F]/total

## estimate convex hull and predict
mdff2 <- findConvex.prop(data$p_P, data$p_C, c("p_P", "p_C"), surface.resolution)
mdff2$p_F <- with(mdff2, 1 - p_P - p_C)

# Get the predicted surface
X <- model.matrix(~0 + p_P + p_C + p_F + p_P:p_C + p_P:p_F + p_C:p_F, data = mdff2)
Y <- (X %*% coeffs)
plot_data <- as.data.frame(X)
plot_data$Y <- Y
plot_data$p_P <- plot_data$p_P * 100
plot_data$p_C <- plot_data$p_C * 100
plot_data$p_F <- plot_data$p_F * 100
```

```r
contour_use <- signif((max(Y) - min(Y))/5, 1)
```

```r

  Apoa1_fathers <- ggplot(plot_data) +
  # Add diagonal lines with solid lines, ensuring they cover the whole plot area
  geom_abline(intercept = 100, slope = -1, color = "grey50") +
  geom_abline(intercept = 80, slope = -1, color = "grey50") +
  geom_abline(intercept = 60, slope = -1, color = "grey50") +
  geom_abline(intercept = 40, slope = -1, color = "grey50") +
  geom_abline(intercept = 20, slope = -1, color = "grey50") +

  # Add other plot layers on top of the lines
  geom_tile(aes(x = p_P, y = p_C, fill = Y)) +
  scale_fill_gradientn(colors = map) +
  geom_contour(data = plot_data, aes(x = p_P, y = p_C, z = Y), na.rm = TRUE, color = "black", binwidth =
  geom_label_contour(data = plot_data, aes(x = p_P, y = p_C, z = Y), size = 3, binwidth = contour_use) +

  # Set theme and labels, remove grid lines and specific axis lines
  theme_minimal() +
  theme(
    panel.grid = element_blank(),       # Remove grid lines
    panel.border = element_blank(),     # Remove the border around the plot panel
    axis.line = element_line(color = "black"),       # Add back the x and y axis lines
    legend.position = "none",            # Remove legend
    axis.text.x = element_text(hjust = -1, vjust = -0.5),  # Adjust horizontal and vertical position fo
    axis.text.y = element_text(hjust = 0.5, vjust = -1.5)    # Adjust horizontal and vertical position
  ) +
  scale_x_continuous(limits = c(0, 100), breaks = seq(0, 100, by = 20)) +  # Set x-axis limits and labe
  scale_y_continuous(limits = c(0, 100), breaks = seq(0, 100, by = 20)) +  # Set y-axis limits and labe
  coord_fixed(ratio = 1) +   # Fix the aspect ratio to ensure alignment of diagonals and axes
  ggtitle("APOA1 (cluster 5)") +
  labs(x = "Protein (%)", y = "Carbohydrate (%)") +

  # Adjust the "Fat" label position
  annotate("text", x = 55, y = 55, label = "Fat (%)", color = "black", angle = -45, hjust = 1)

# Display the plot
Apoa1_fathers
```

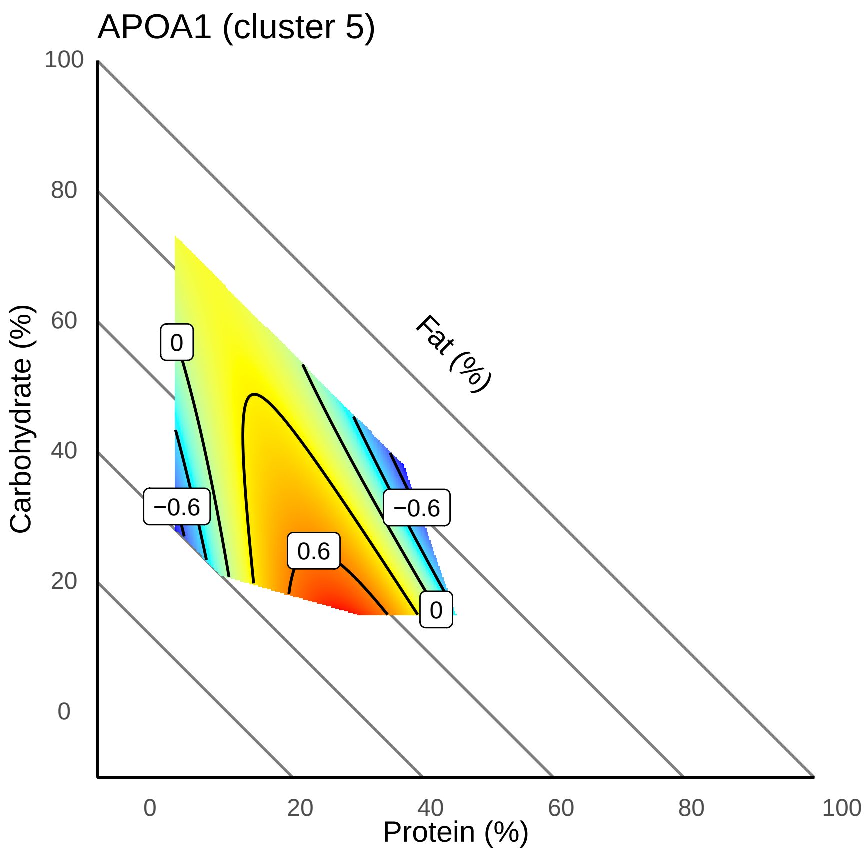

###### 4.2.5.1 APOA1

```
Fn1 <- subset(Cluster5, Protein == "Fn1")
Fn1_joined <- tidy_protein_table_fathers[tidy_protein_table_fathers$proteins %in%
    Fn1, ]

Fn1_joined$pP <- (Fn1_joined$Percent_prot/100)
Fn1_joined$pC <- (Fn1_joined$Percent_carb/100)
Fn1_joined$pF <- (Fn1_joined$Percent_fat/100)

Fn1_joined$LogIntensity_z <- scale(Fn1_joined$LogIntensity)
model <- lm(LogIntensity_z ~ 0 + pP + pC + pF + pP:pC + pP:pF + pC:pF, data = Fn1_joined)

coeffs <- coef(model)
data = metadata
p_P = "Percent_prot"
p_C = "Percent_carb"
p_F = "Percent_fat"

mixture.surface <- function(data, coeffs, p_P = "p_P", p_C = "p_C", p_F = "p_F") {
}

# Set the resolution of the surface
```

```r
surface.resolution <- 501

# How many values to round surface
round.surf <- 3

# This specifies the color scheme for surface - it is actually a function that
# returns a function
rgb.palette <- colorRampPalette(c("blue", "cyan", "yellow", "red"), space = "Lab",
    interpolate = "linear")

# How many different colours should we use on the plot
no.cols <- 256

# Get the colors to use from the palette specified above
map <- rgb.palette(no.cols)

# How many levels should there be on the surface
nlev <- 5

# Labels for each
labels <- c("Protein (%)", "Carbohydrate (%)", "Fat (%)")

## plot the RMT surface
iso.lines <- seq(1, 0, -0.2)

# Set the layout
par(mfrow = c(1, 1), mar = c(5, 5, 5, 1))

# Make sure the proportions are closed off to 1
total <- (data[, p_P] + data[, p_C] + data[, p_F])
data$p_P <- data[, p_P]/total
data$p_C <- data[, p_C]/total
data$p_F <- data[, p_F]/total

## estimate convex hull and predict
mdff2 <- findConvex.prop(data$p_P, data$p_C, c("p_P", "p_C"), surface.resolution)
mdff2$p_F <- with(mdff2, 1 - p_P - p_C)

# Get the predicted surface
X <- model.matrix(~0 + p_P + p_C + p_F + p_P:p_C + p_P:p_F + p_C:p_F, data = mdff2)
Y <- (X %*% coeffs)
plot_data <- as.data.frame(X)
plot_data$Y <- Y
plot_data$p_P <- plot_data$p_P * 100
plot_data$p_C <- plot_data$p_C * 100
plot_data$p_F <- plot_data$p_F * 100
contour_use <- signif((max(Y) - min(Y))/5, 1)
```

```r
  Fn1_fathers <- ggplot(plot_data) +
  # Add diagonal lines with solid lines, ensuring they cover the whole plot area
  geom_abline(intercept = 100, slope = -1, color = "grey50") +
  geom_abline(intercept = 80, slope = -1, color = "grey50") +
```

```
  geom_abline(intercept = 60, slope = -1, color = "grey50") +
  geom_abline(intercept = 40, slope = -1, color = "grey50") +
  geom_abline(intercept = 20, slope = -1, color = "grey50") +

  # Add other plot layers on top of the lines
  geom_tile(aes(x = p_P, y = p_C, fill = Y)) +
  scale_fill_gradientn(colors = map) +
  geom_contour(data = plot_data, aes(x = p_P, y = p_C, z = Y), na.rm = TRUE, color = "black", binwidth =
  geom_label_contour(data = plot_data, aes(x = p_P, y = p_C, z = Y), size = 3, binwidth = contour_use) +

  # Set theme and labels, remove grid lines and specific axis lines
  theme_minimal() +
  theme(
    panel.grid = element_blank(),       # Remove grid lines
    panel.border = element_blank(),     # Remove the border around the plot panel
    axis.line = element_line(color = "black"),       # Add back the x and y axis lines
    legend.position = "none",             # Remove legend
    axis.text.x = element_text(hjust = -1, vjust = -0.5),  # Adjust horizontal and vertical position fo
    axis.text.y = element_text(hjust = 0.5, vjust = -1.5)    # Adjust horizontal and vertical position
  ) +
  scale_x_continuous(limits = c(0, 100), breaks = seq(0, 100, by = 20)) +  # Set x-axis limits and labe
  scale_y_continuous(limits = c(0, 100), breaks = seq(0, 100, by = 20)) +  # Set y-axis limits and labe
  coord_fixed(ratio = 1) +   # Fix the aspect ratio to ensure alignment of diagonals and axes
  ggtitle("FN1 (cluster 5)") +
  labs(x = "Protein (%)", y = "Carbohydrate (%)") +

  # Adjust the "Fat" label position
  annotate("text", x = 55, y = 55, label = "Fat (%)", color = "black", angle = -45, hjust = 1)

# Display the plot
Fn1_fathers
```

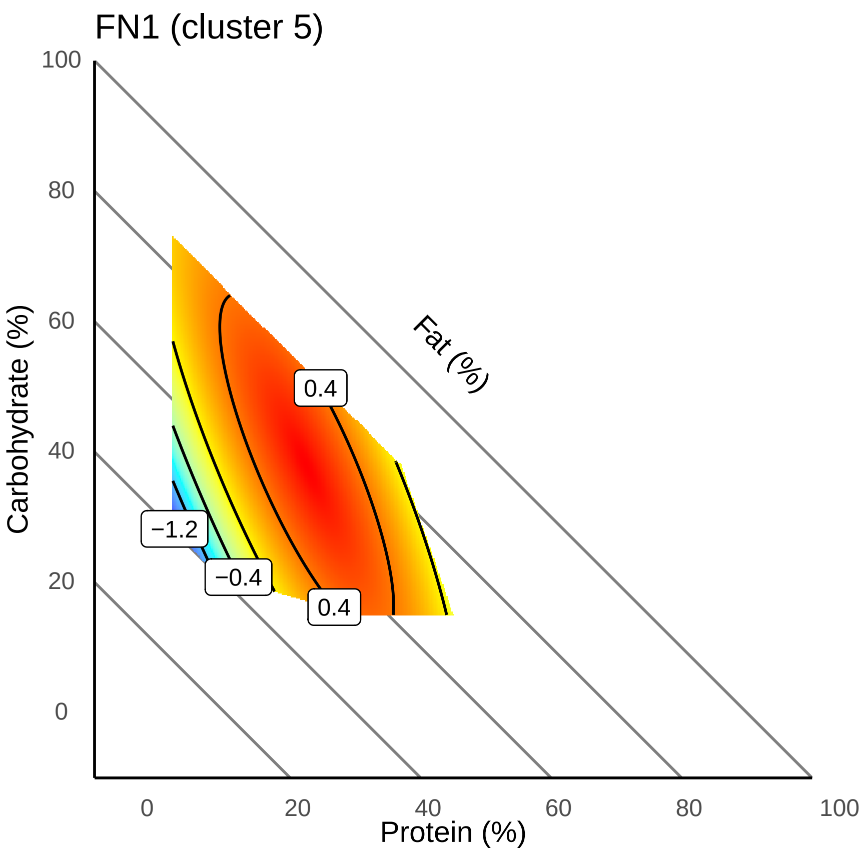

#### 4.2.5.2 FN1

```
p <- (Fn1_fathers | Apoa1_fathers)
p
```

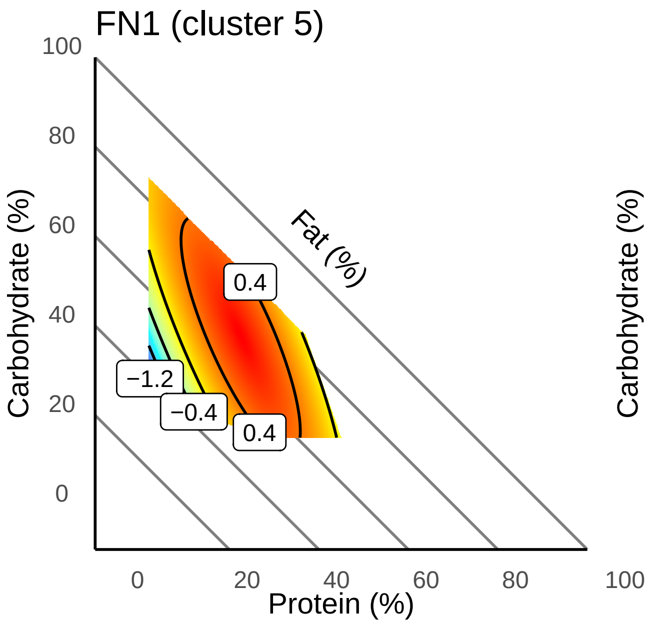

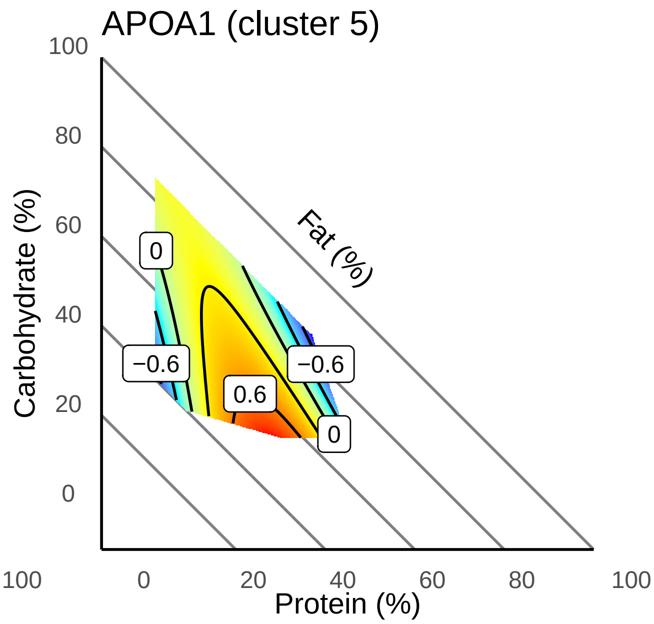

###### 4.2.5.3 Combined

### 5 F1 female offspring

This section contains all the additive/linear and interaction/quadratic models. Including models with dietary intake as a co-variate.

There were no significantly affected proteins other than in the protein + intake model (reported in main text).

#### 5.1 Protein and carbs (%)

##### 5.1.1 Additive model

```
# create design matrix
Percent_prot = as.numeric(SummarizedExperiment_females$Percent_prot)
Percent_carb = as.numeric(SummarizedExperiment_females$Percent_carb)

design7 <- model.matrix(~Percent_prot + Percent_carb)

data_sample_names <- colnames(SummarizedExperiment_females)
design_sample_names <- rownames(design7)

rownames(design7) <- data_sample_names
```

```
fit7 <- lmFit(assay(SummarizedExperiment_females), design7)
fit7a <- eBayes(fit7)

table7 <- topTable(fit7a, sort = "none", number = Inf, adjust = "fdr")

count(table7$adj.P.Val < 0.05)
```

```
## [1] 0
```

```
hist(table7$P.Value)
```

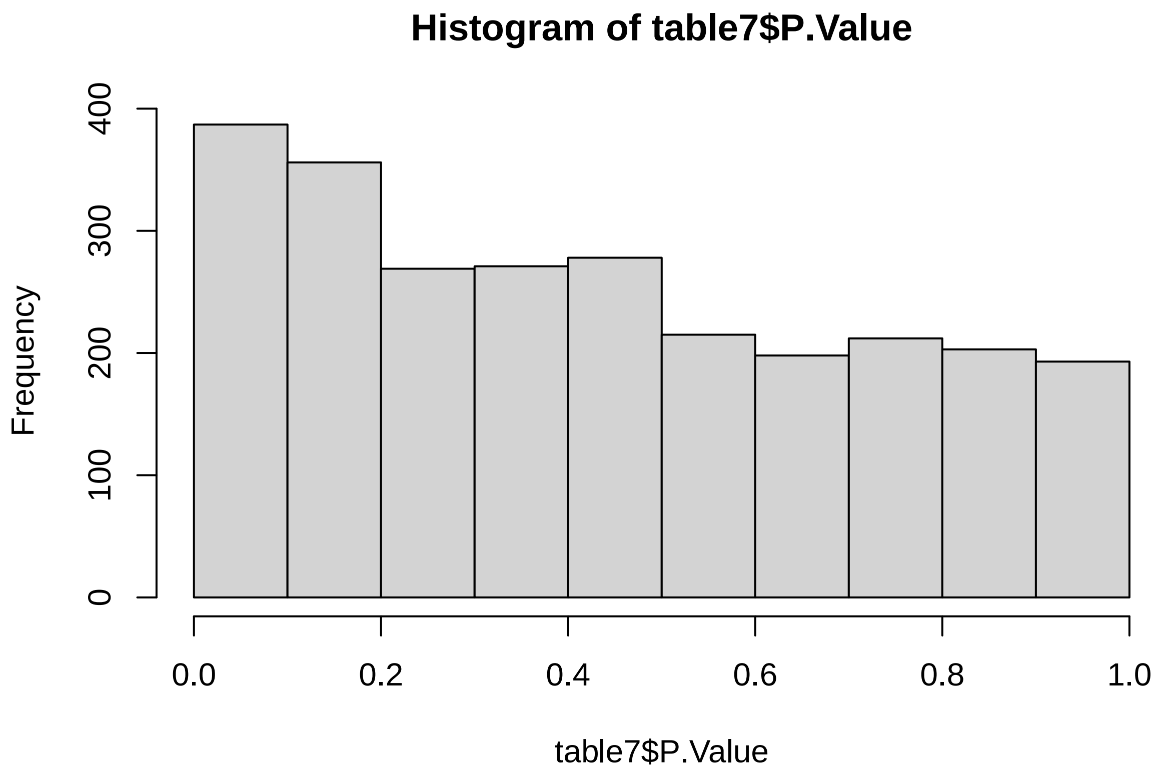

```
# 0 DE
```

##### 5.1.2 Additive model + intake

```
# create design matrix
Percent_prot = as.numeric(SummarizedExperiment_females$Percent_prot)
Percent_carb = as.numeric(SummarizedExperiment_females$Percent_carb)
intake = as.numeric(SummarizedExperiment_females$Food_intake)

design7a <- model.matrix(~Percent_prot + Percent_carb + intake)
```

```
data_sample_names <- colnames(SummarizedExperiment_females)
design_sample_names <- rownames(design7a)

rownames(design7a) <- data_sample_names

fit7a <- lmFit(assay(SummarizedExperiment_females), design7a)
fit7b <- eBayes(fit7a)

table7a <- topTable(fit7b, sort = "none", number = Inf, adjust = "fdr")

count(table7a$adj.P.Val < 0.05)
```

```
## [1] 0
```

```
hist(table7a$P.Value)
```

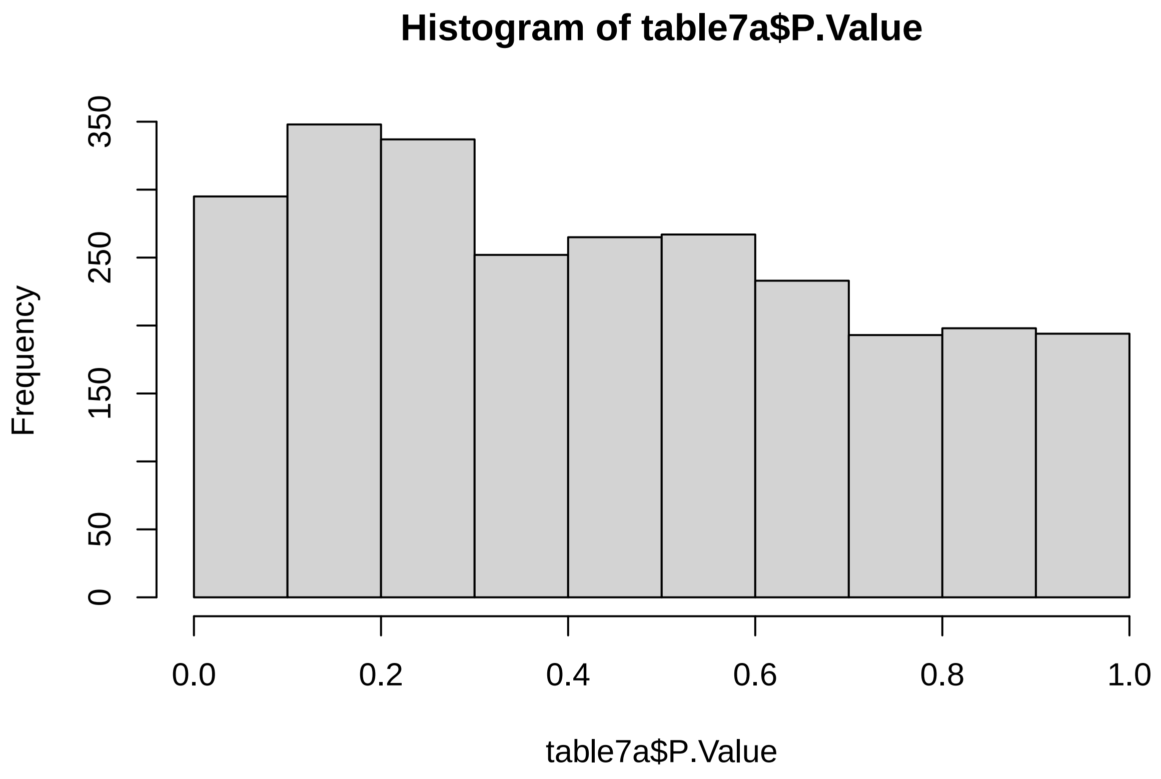

```
# 0 DE
```

###### 5.1.3 Interaction model

```
# create design matrix
Percent_prot = as.numeric(SummarizedExperiment_females$Percent_prot)
```

```r
Percent_carb = as.numeric(SummarizedExperiment_females$Percent_carb)

design8 <- model.matrix(~Percent_prot * Percent_carb)

data_sample_names <- colnames(SummarizedExperiment_females)
design_sample_names <- rownames(design8)

rownames(design8) <- data_sample_names

fit8 <- lmFit(assay(SummarizedExperiment_females), design8)
fit8a <- eBayes(fit8)

table8 <- topTable(fit8a, sort = "none", number = Inf, adjust = "fdr")
count(table8$adj.P.Val < 0.05)
```

```
## [1] 0
```

```r
hist(table8$P.Value)
```

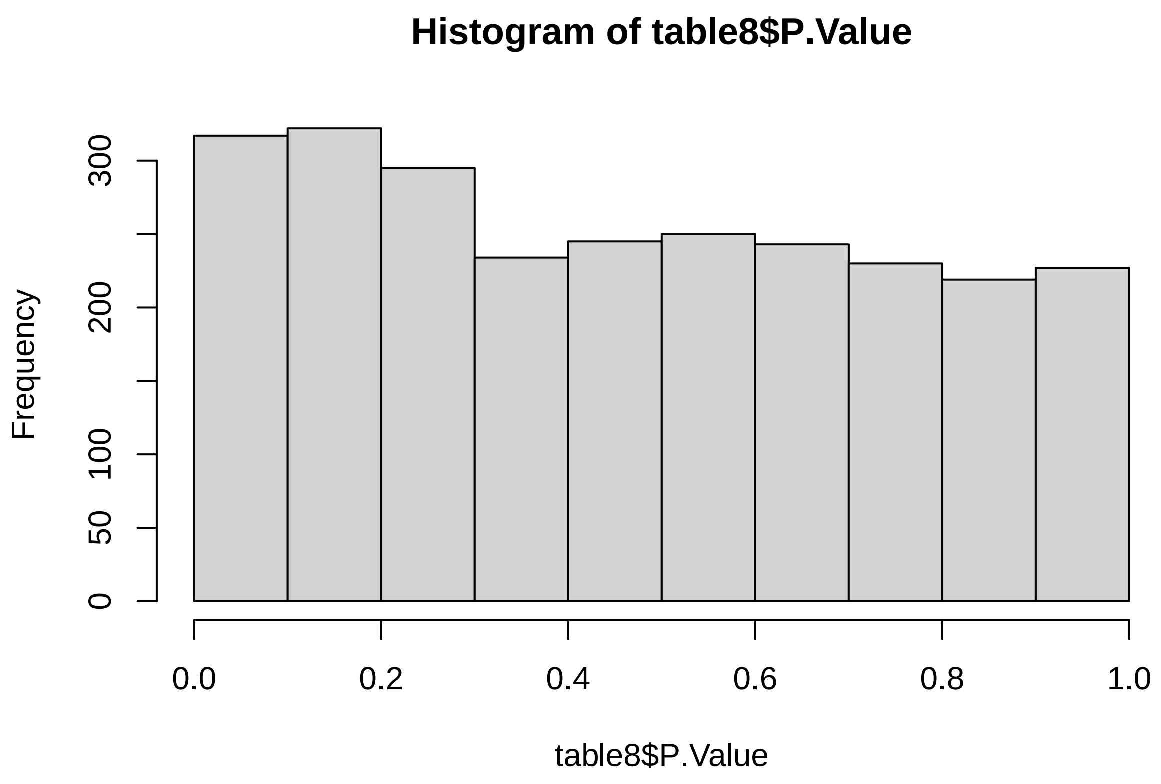

```r
# 0 DE
```

##### 5.1.4 Interaction model + intake

```
# create design matrix
Percent_prot = as.numeric(SummarizedExperiment_females$Percent_prot)
Percent_carb = as.numeric(SummarizedExperiment_females$Percent_carb)
intake = as.numeric(SummarizedExperiment_females$Food_intake)

design8a <- model.matrix(~Percent_prot * Percent_carb + intake)

data_sample_names <- colnames(SummarizedExperiment_females)
design_sample_names <- rownames(design8a)

rownames(design8a) <- data_sample_names

fit8b <- lmFit(assay(SummarizedExperiment_females), design8a)
fit8b <- eBayes(fit8b)

table8a <- topTable(fit8b, sort = "none", number = Inf, adjust = "fdr")
count(table8a$adj.P.Val < 0.05)
```

```
## [1] 0
```

```
hist(table8a$P.Value)
```

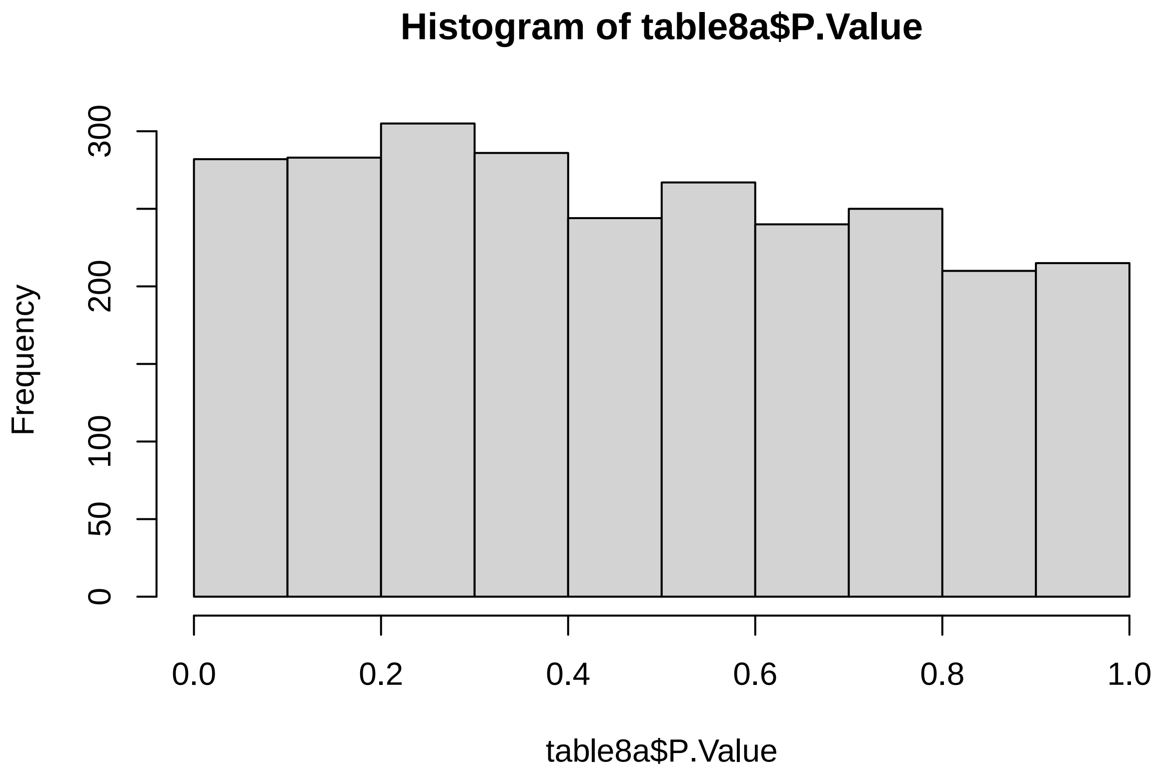

```
# 0 DE
```

#### 5.2 Protein and fat(%)

##### 5.2.1 Additive model

```
# create design matrix
Percent_prot = as.numeric(SummarizedExperiment_females$Percent_prot)
Percent_fat = as.numeric(SummarizedExperiment_females$Percent_fat)

design9 <- model.matrix(~Percent_prot + Percent_carb)

data_sample_names <- colnames(SummarizedExperiment_females)
design_sample_names <- rownames(design9)

rownames(design9) <- data_sample_names

fit9 <- lmFit(assay(SummarizedExperiment_females), design9)
fit9a <- eBayes(fit9)

table9 <- topTable(fit9a, sort = "none", number = Inf, adjust = "fdr")

count(table9$adj.P.Val < 0.05)
```

```
## [1] 0
```

```
hist(table9$P.Value)
```

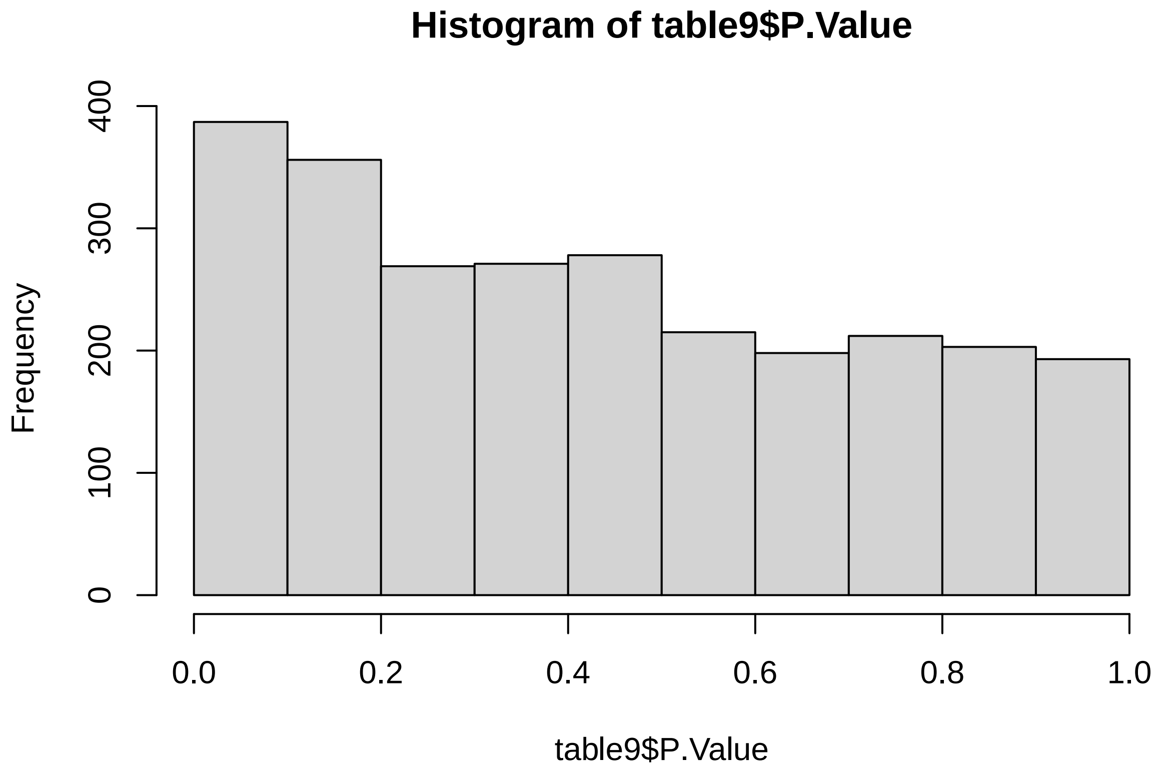

```
# 0 DE
```

##### 5.2.2 Additive model + intake

```
# create design matrix
Percent_prot = as.numeric(SummarizedExperiment_females$Percent_prot)
Percent_fat = as.numeric(SummarizedExperiment_females$Percent_fat)
intake = as.numeric(SummarizedExperiment_females$Food_intake)

design9a <- model.matrix(~Percent_prot + Percent_fat + intake)

data_sample_names <- colnames(SummarizedExperiment_females)
design_sample_names <- rownames(design9a)

rownames(design9a) <- data_sample_names

fit9a <- lmFit(assay(SummarizedExperiment_females), design9a)
fit9b <- eBayes(fit9a)

table9a <- topTable(fit9b, sort = "none", number = Inf, adjust = "fdr")

count(table9a$adj.P.Val < 0.05)
```

```
## [1] 0
```

```
hist(table9a$P.Value)
```

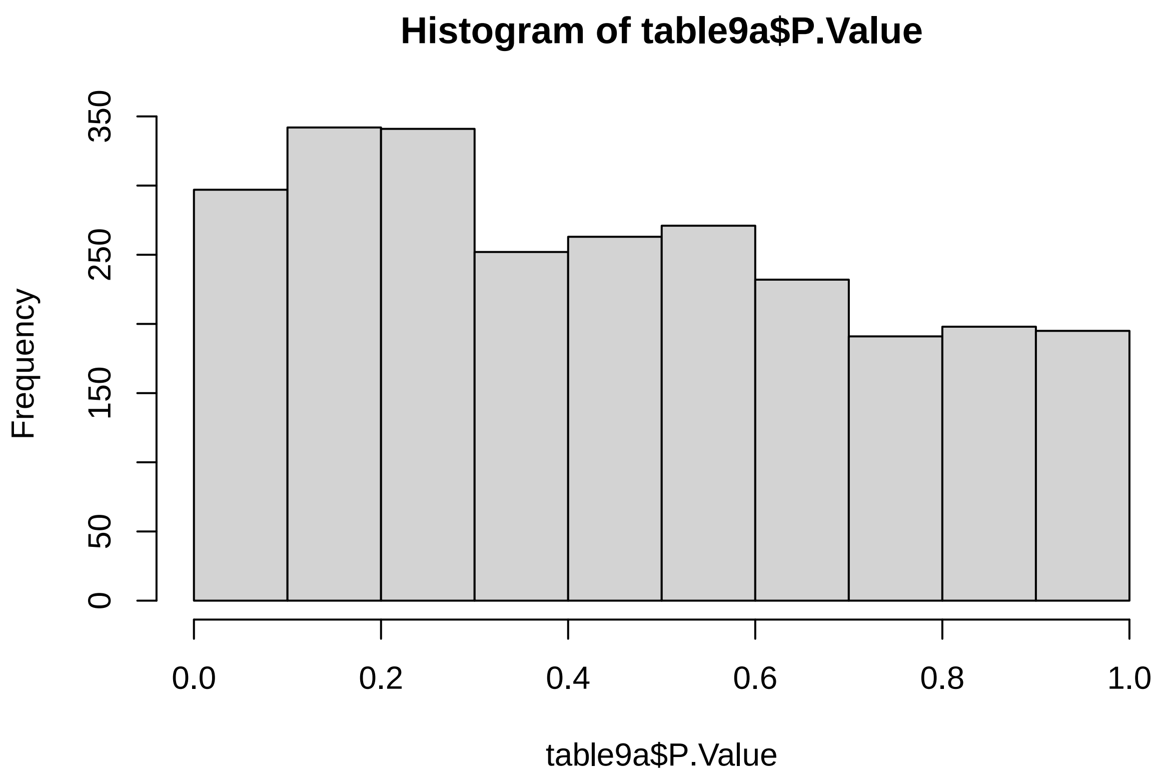

```
# 0 DE
```

##### 5.2.3 Interaction model

```
# create design matrix
Percent_prot = as.numeric(SummarizedExperiment_females$Percent_prot)
Percent_fat = as.numeric(SummarizedExperiment_females$Percent_fat)

design10 <- model.matrix(~Percent_prot * Percent_fat)

data_sample_names <- colnames(SummarizedExperiment_females)
design_sample_names <- rownames(design10)

rownames(design10) <- data_sample_names

fit10 <- lmFit(assay(SummarizedExperiment_females), design10)
fit10a <- eBayes(fit10)

table10 <- topTable(fit10a, sort = "none", number = Inf, adjust = "fdr")
count(table10$adj.P.Val < 0.05)
```

```
## [1] 0
```

```r
hist(table10$P.Value)
```

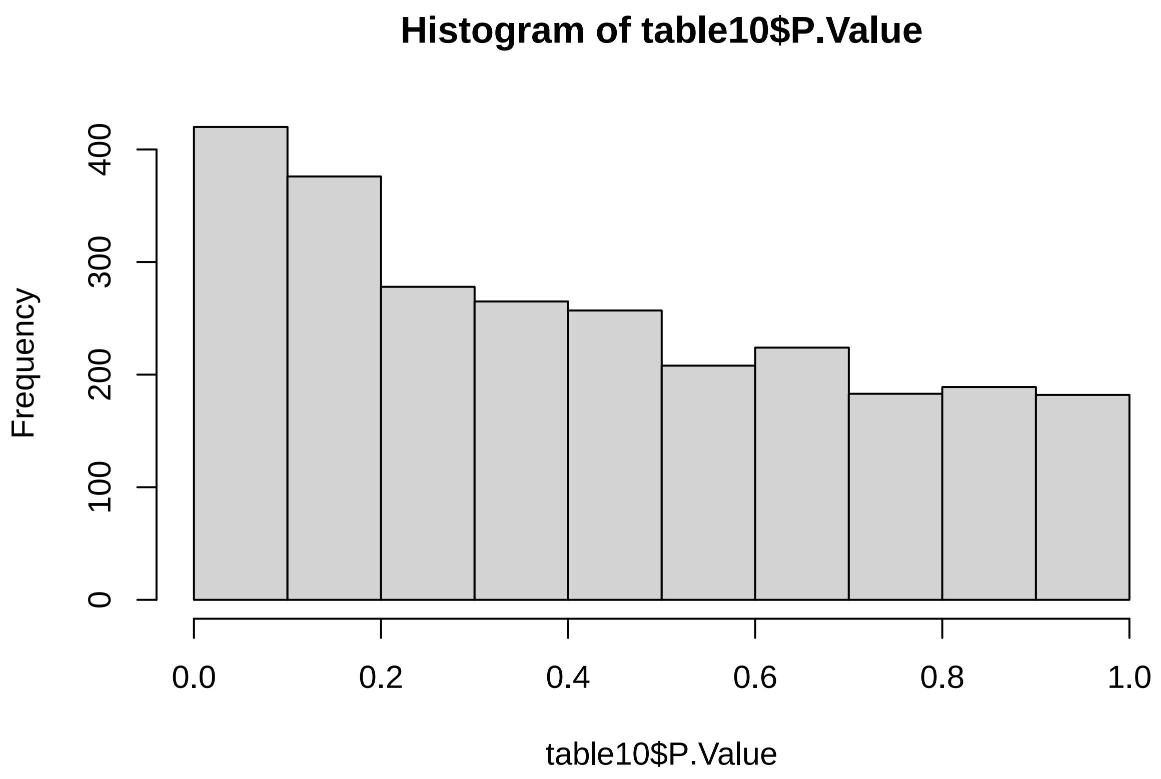

```r
# 0 DE
```

###### 5.2.4 Interaction model + intake

```r
# create design matrix
Percent_prot = as.numeric(SummarizedExperiment_females$Percent_prot)
Percent_fat = as.numeric(SummarizedExperiment_females$Percent_fat)
intake = as.numeric(SummarizedExperiment_females$Food_intake)

design10a <- model.matrix(~Percent_prot * Percent_fat + intake)

data_sample_names <- colnames(SummarizedExperiment_females)
design_sample_names <- rownames(design10a)

rownames(design10a) <- data_sample_names

fit10b <- lmFit(assay(SummarizedExperiment_females), design10a)
fit10b <- eBayes(fit10b)

table10a <- topTable(fit10b, sort = "none", number = Inf, adjust = "fdr")
count(table10a$adj.P.Val < 0.05)
```

```
## [1] 0
```

```r
hist(table10a$P.Value)
```

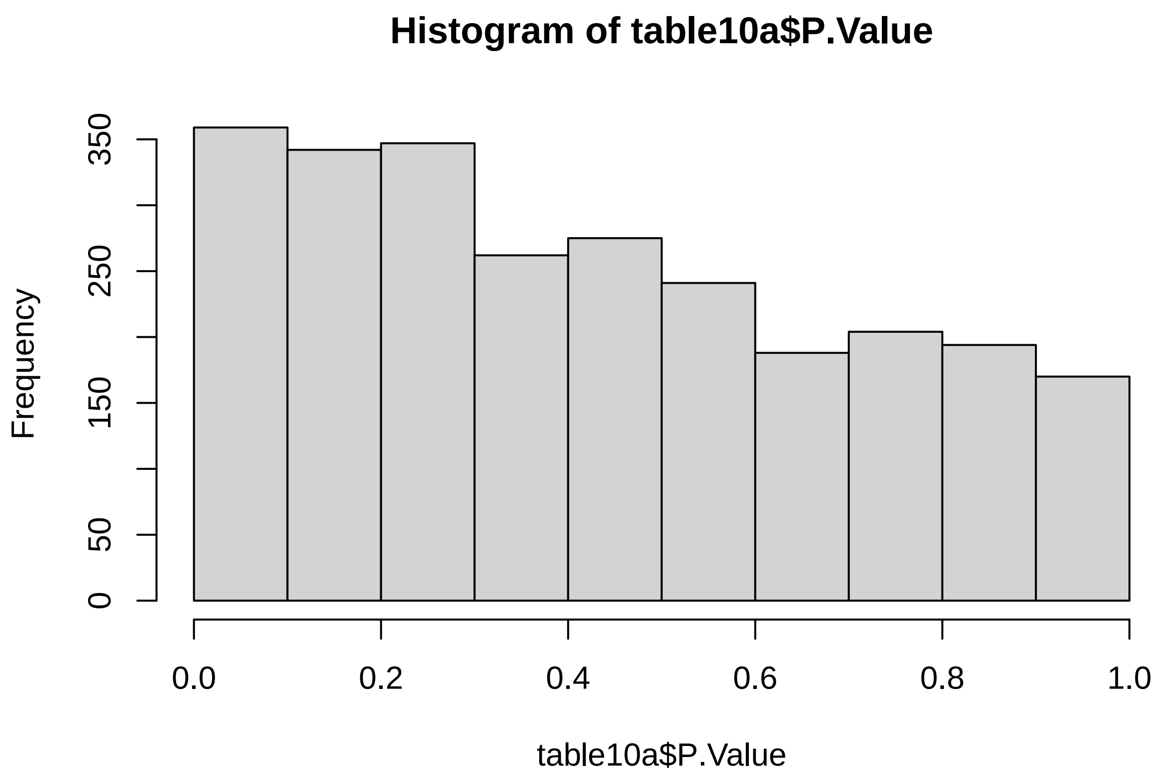

```r
# 0 DE
```

#### 5.3 Carbs and fat(%)

##### 5.3.1 Additive model

```r
# create design matrix
Percent_carb = as.numeric(SummarizedExperiment_females$Percent_carb)
Percent_fat = as.numeric(SummarizedExperiment_females$Percent_fat)

design11 <- model.matrix(~Percent_carb + Percent_fat)

data_sample_names <- colnames(SummarizedExperiment_females)
design_sample_names <- rownames(design11)

rownames(design11) <- data_sample_names

fit11 <- lmFit(assay(SummarizedExperiment_females), design11)
fit11a <- eBayes(fit11)
```

```
table11 <- topTable(fit11a, sort = "none", number = Inf, adjust = "fdr")

count(table11$adj.P.Val < 0.05)
```

```
## [1] 0
```

```
hist(table11$P.Value)
```

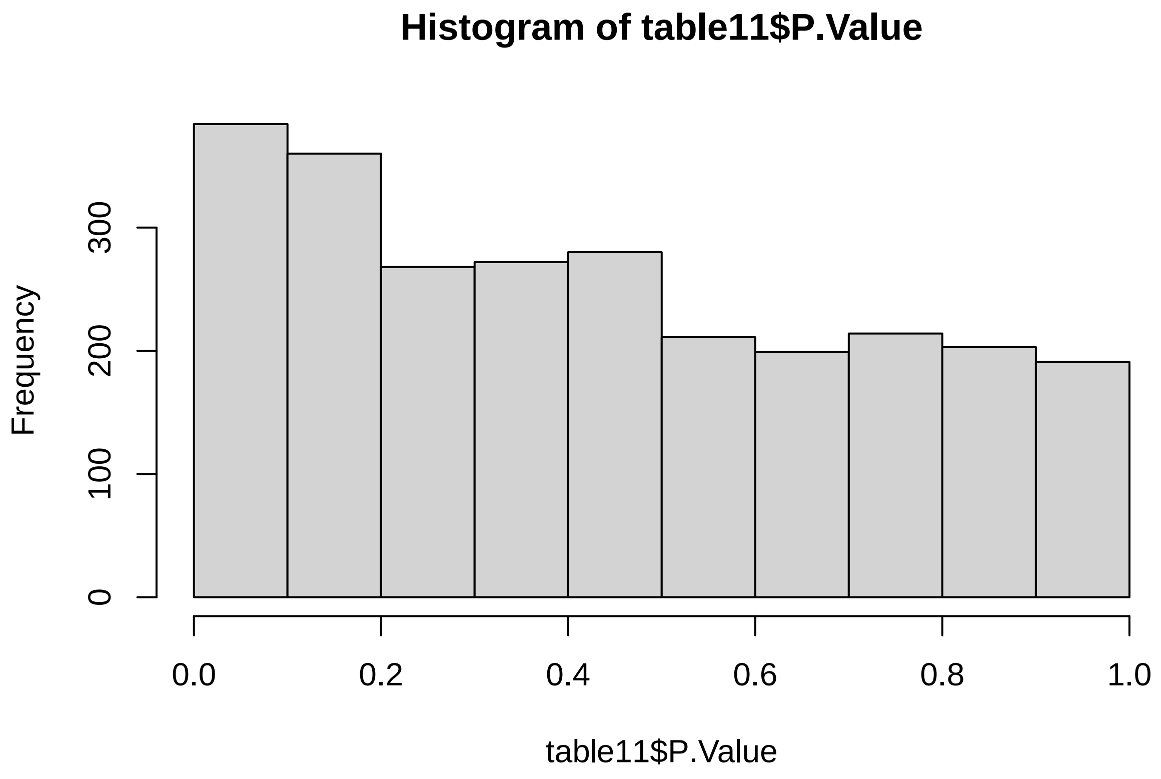

```
# 0 DE
```

##### 5.3.2 Additive model + intake

```
# create design matrix
Percent_fat = as.numeric(SummarizedExperiment_females$Percent_fat)
Percent_carb = as.numeric(SummarizedExperiment_females$Percent_carb)
intake = as.numeric(SummarizedExperiment_females$Food_intake)

design11a <- model.matrix(~Percent_carb + Percent_fat + intake)

data_sample_names <- colnames(SummarizedExperiment_females)
design_sample_names <- rownames(design11a)
```

```r
rownames(design11a) <- data_sample_names

fit11a <- lmFit(assay(SummarizedExperiment_females), design11a)
fit11b <- eBayes(fit11a)

table11a <- topTable(fit11b, sort = "none", number = Inf, adjust = "fdr")

count(table11a$adj.P.Val < 0.05)
```

```
## [1] 0
```

```r
hist(table11a$P.Value)
```

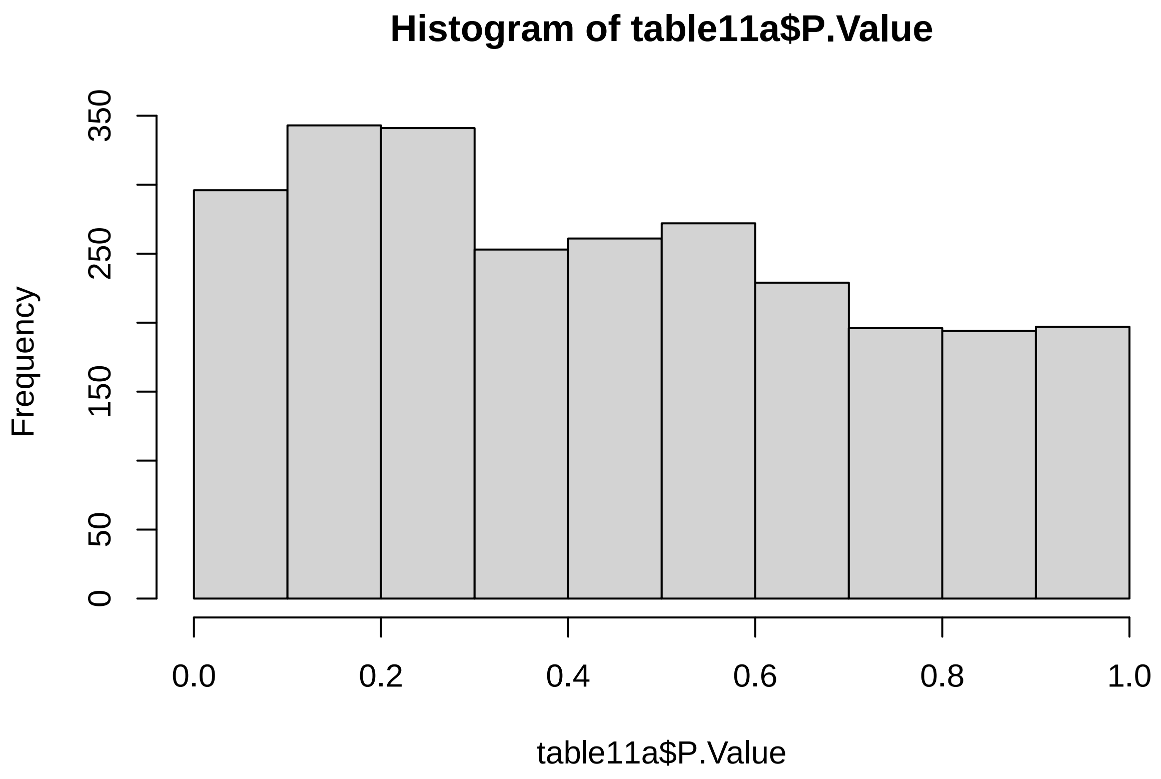

```r
# 0 DE
```

##### 5.3.3 Interaction model

```r
# create design matrix
Percent_fat = as.numeric(SummarizedExperiment_females$Percent_fat)
Percent_carb = as.numeric(SummarizedExperiment_females$Percent_carb)

design12 <- model.matrix(~Percent_carb + Percent_fat)
```

```r
data_sample_names <- colnames(SummarizedExperiment_females)
design_sample_names <- rownames(design12)

rownames(design12) <- data_sample_names

fit12 <- lmFit(assay(SummarizedExperiment_females), design12)
fit12a <- eBayes(fit12)

table12 <- topTable(fit12a, sort = "none", number = Inf, adjust = "fdr")
count(table12$adj.P.Val < 0.05)
```

```
## [1] 0
```

```r
hist(table12$P.Value)
```

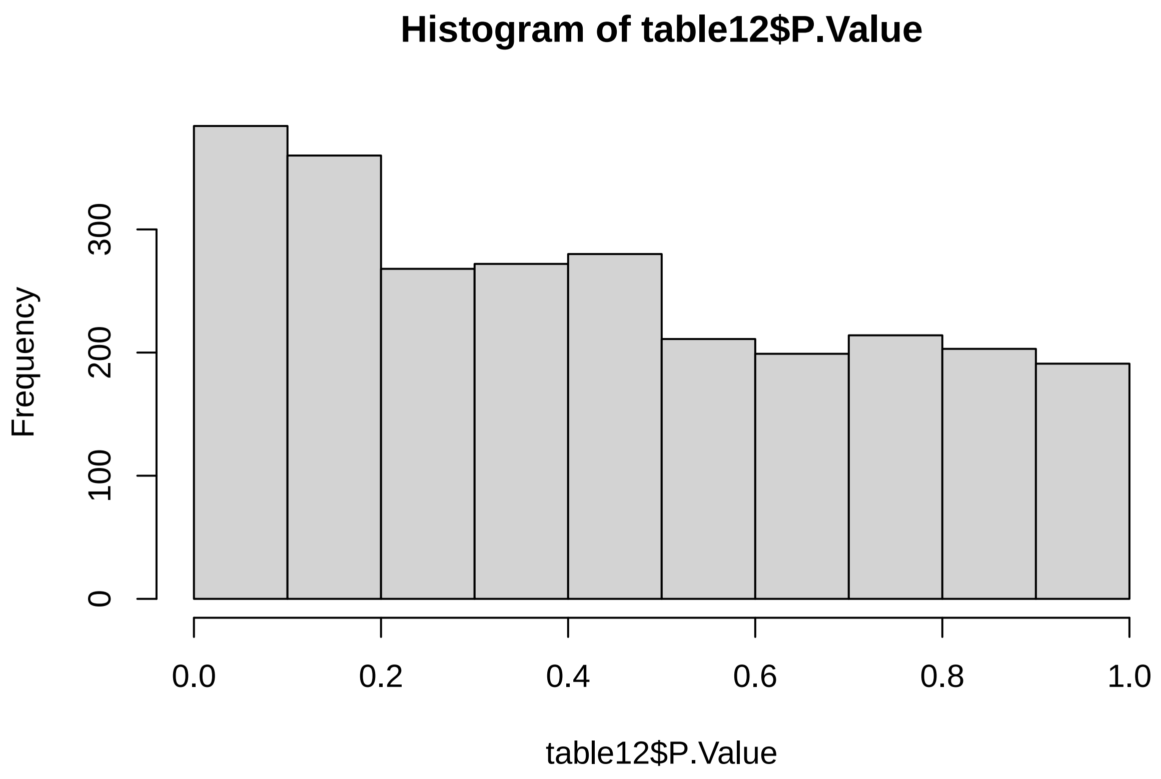

```r
# 0 DE
```

###### 5.3.4 Interaction model + intake

```r
# create design matrix
Percent_fat = as.numeric(SummarizedExperiment_females$Percent_fat)
Percent_carb = as.numeric(SummarizedExperiment_females$Percent_carb)
```

```r
intake = as.numeric(SummarizedExperiment_females$Food_intake)

design12a <- model.matrix(~Percent_carb * Percent_fat + intake)

data_sample_names <- colnames(SummarizedExperiment_females)
design_sample_names <- rownames(design12a)

rownames(design12a) <- data_sample_names

fit12b <- lmFit(assay(SummarizedExperiment_females), design12a)
fit12b <- eBayes(fit12b)

table12a <- topTable(fit12b, sort = "none", number = Inf, adjust = "fdr")
count(table12a$adj.P.Val < 0.05)
```

```
## [1] 0
```

```r
hist(table12a$P.Value)
```

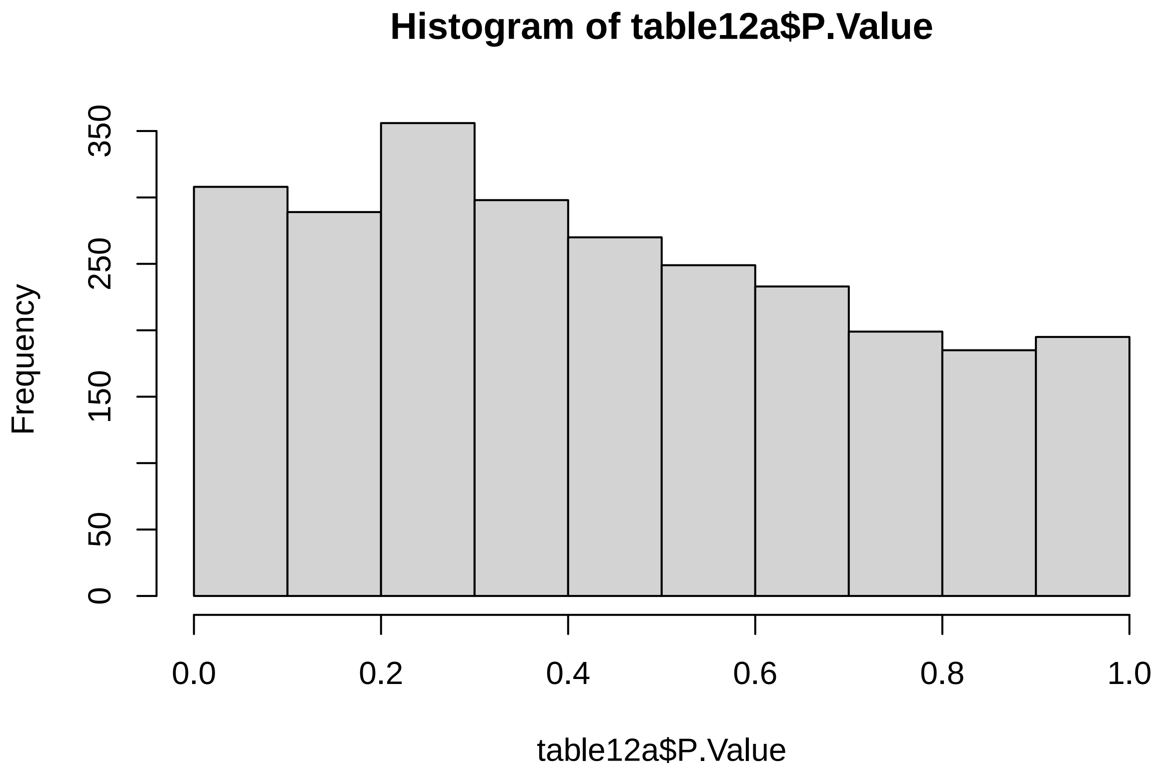

```r
# 0 DE
```

#### 5.4 Protein (%)

```
# create design matrix
Percent_prot = log2(as.numeric(SummarizedExperiment_females$Percent_prot))

design13 <- model.matrix(~Percent_prot)

data_sample_names <- colnames(SummarizedExperiment_females)
design_sample_names <- rownames(design13)

rownames(design13) <- data_sample_names

fit13 <- lmFit(assay(SummarizedExperiment_females), design13)
fit13a <- eBayes(fit13)

table13 <- topTable(fit13a, sort = "none", number = Inf, adjust = "fdr")

count(table13$adj.P.Val < 0.05)
```

```
## [1] 0
```

```
hist(table13$P.Value)
```

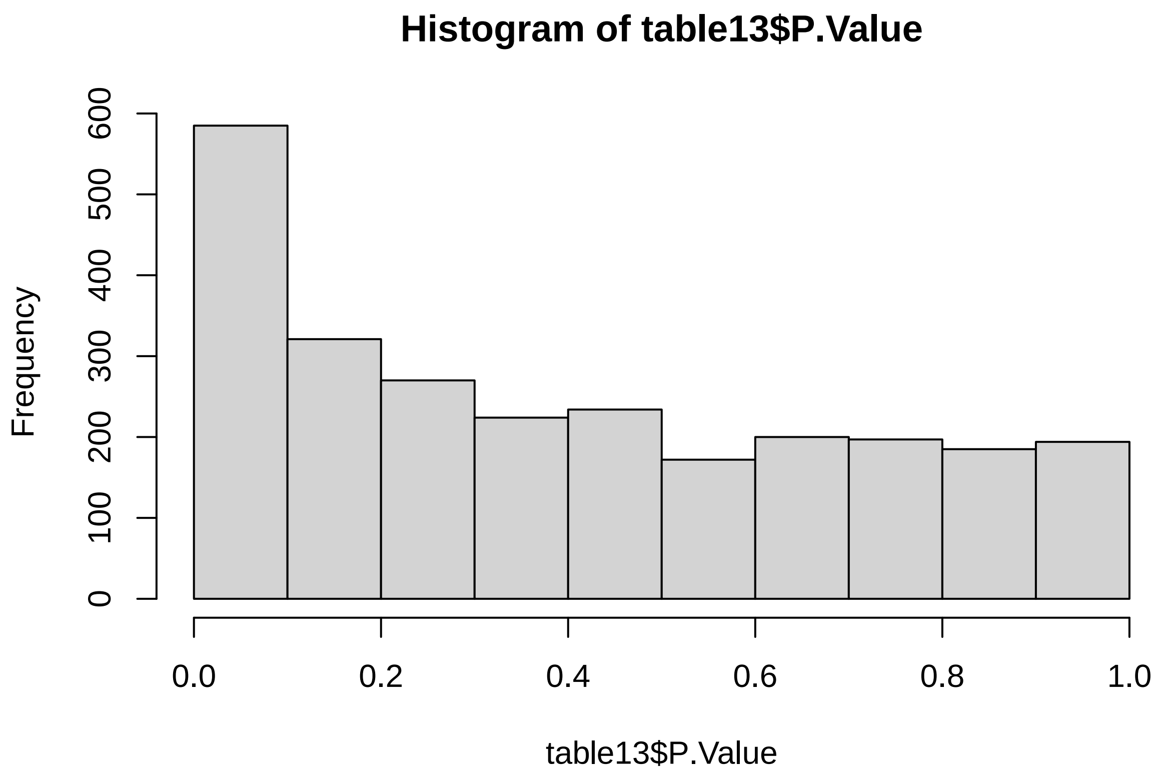

##### 5.4.1 Protein + intake

This is the model where Bsg is significantly affected by diet (see main text)

```
# create design matrix
Percent_prot = log2(as.numeric(SummarizedExperiment_females$Percent_prot))
intake = log2(as.numeric(SummarizedExperiment_females$Food_intake))

design13a <- model.matrix(~Percent_prot + intake)

data_sample_names <- colnames(SummarizedExperiment_females)
design_sample_names <- rownames(design13a)

rownames(design13a) <- data_sample_names

fit13a <- lmFit(assay(SummarizedExperiment_females), design13a)
fit13b <- eBayes(fit13a)

table13a <- topTable(fit13b, sort = "none", number = Inf, adjust = "fdr")

count(table13a$adj.P.Val < 0.05)
```

```
## [1] 1
```

```
hist(table13a$P.Value)
```

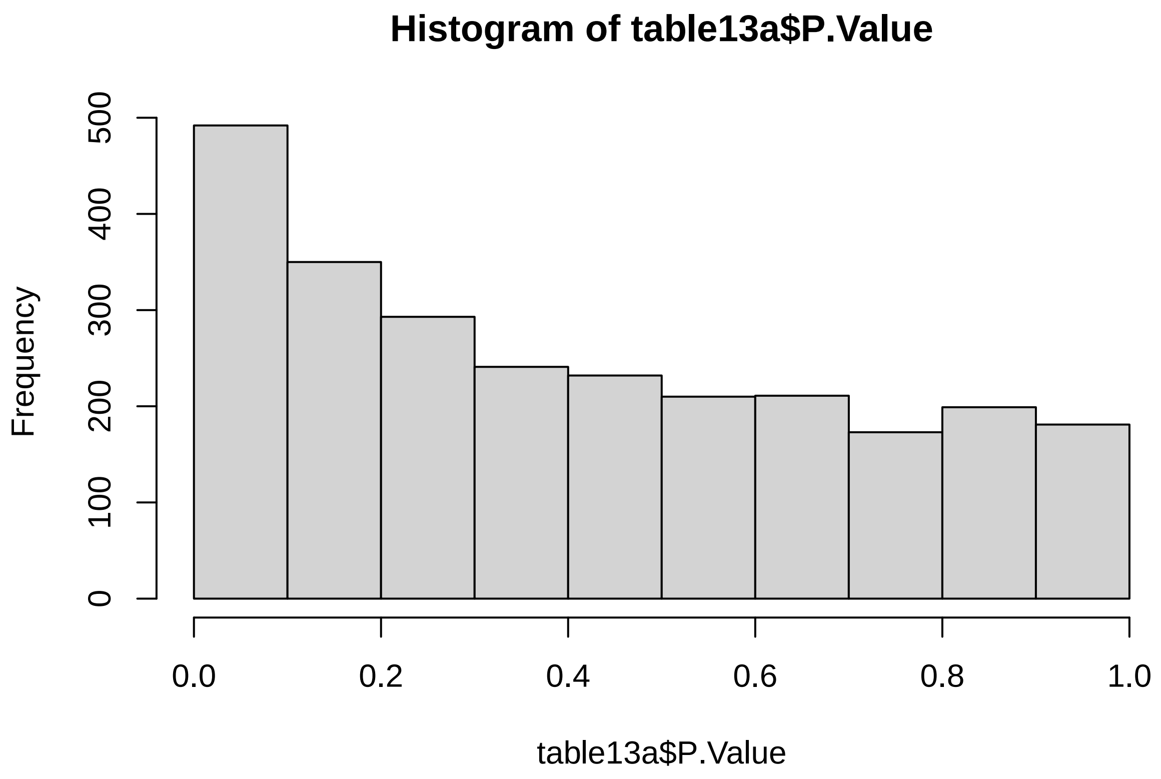

#### 5.5 Carbs (%)

```r
# create design matrix
Percent_carb = log2(as.numeric(SummarizedExperiment_females$Percent_carb))

design14 <- model.matrix(~Percent_carb)

data_sample_names <- colnames(SummarizedExperiment_females)
design_sample_names <- rownames(design14)

rownames(design14) <- data_sample_names

fit14 <- lmFit(assay(SummarizedExperiment_females), design14)
fit14a <- eBayes(fit14)

table14 <- topTable(fit14a, sort = "none", number = Inf, adjust = "fdr")

count(table14$adj.P.Val < 0.05)
```

```
## [1] 0
```

```r
hist(table14$P.Value)
```

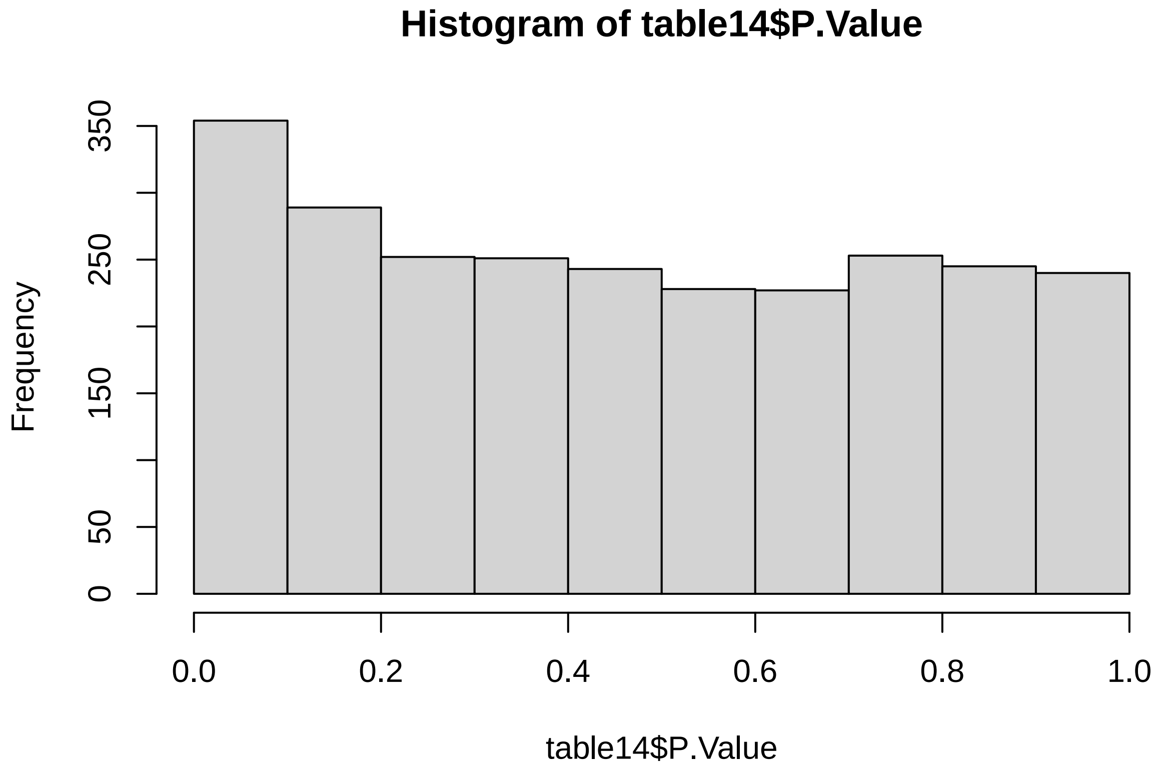

##### 5.5.1 Carb + intake

```
# create design matrix
Percent_carb = log2(as.numeric(SummarizedExperiment_females$Percent_carb))
intake = log2(as.numeric(SummarizedExperiment_females$Food_intake))

design14a <- model.matrix(~Percent_carb + intake)

data_sample_names <- colnames(SummarizedExperiment_females)
design_sample_names <- rownames(design14a)

rownames(design14a) <- data_sample_names

fit14a <- lmFit(assay(SummarizedExperiment_females), design14a)
fit14b <- eBayes(fit14a)

table14a <- topTable(fit14b, sort = "none", number = Inf, adjust = "fdr")

count(table14a$adj.P.Val < 0.05)
```

```
## [1] 0
```

```
hist(table14a$P.Value)
```

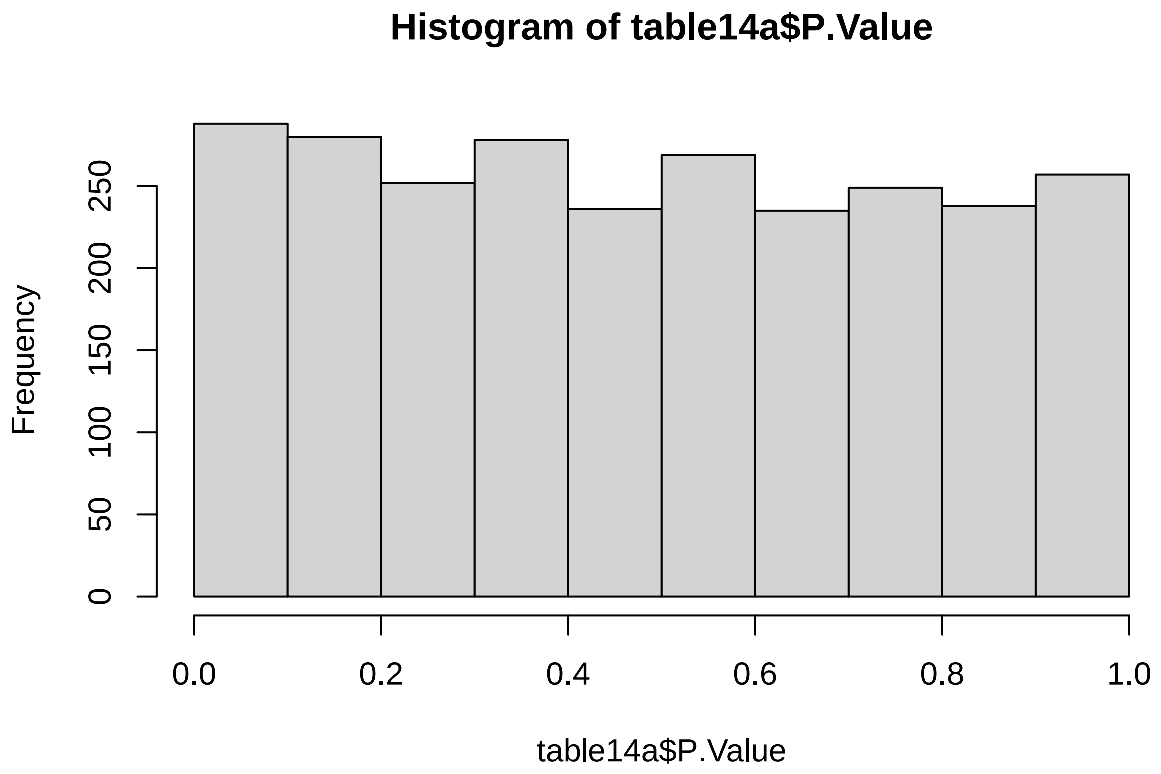

##### 5.6 Fat (%)

```
# create design matrix
Percent_fat = log2(as.numeric(SummarizedExperiment_females$Percent_fat))

design15 <- model.matrix(~Percent_fat)

data_sample_names <- colnames(SummarizedExperiment_females)
design_sample_names <- rownames(design15)

rownames(design15) <- data_sample_names

fit15 <- lmFit(assay(SummarizedExperiment_females), design15)
fit15a <- eBayes(fit15)

table15 <- topTable(fit15a, sort = "none", number = Inf, adjust = "fdr")

count(table15$adj.P.Val < 0.05)
```

```
## [1] 0
```

```
hist(table15$P.Value)
```

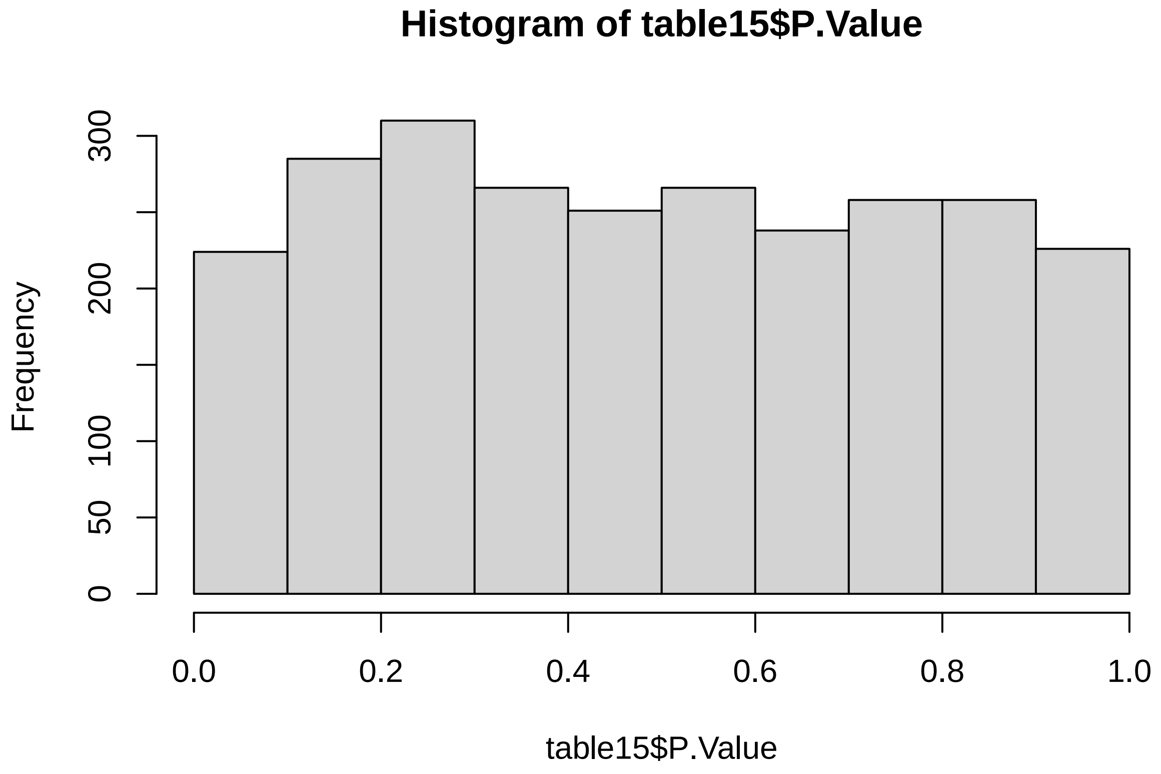

###### 5.6.1 Fat + intake

```
# create design matrix
Percent_fat = log2(as.numeric(SummarizedExperiment_females$Percent_fat))
intake = log2(as.numeric(SummarizedExperiment_females$Food_intake))

design15a <- model.matrix(~Percent_fat + intake)

data_sample_names <- colnames(SummarizedExperiment_females)
design_sample_names <- rownames(design14a)

rownames(design15a) <- data_sample_names

fit15a <- lmFit(assay(SummarizedExperiment_females), design15a)
fit15b <- eBayes(fit15a)

table15a <- topTable(fit15b, sort = "none", number = Inf, adjust = "fdr")

count(table15a$adj.P.Val < 0.05)
```

```
## [1] 0
```

```
hist(table15a$P.Value)
```

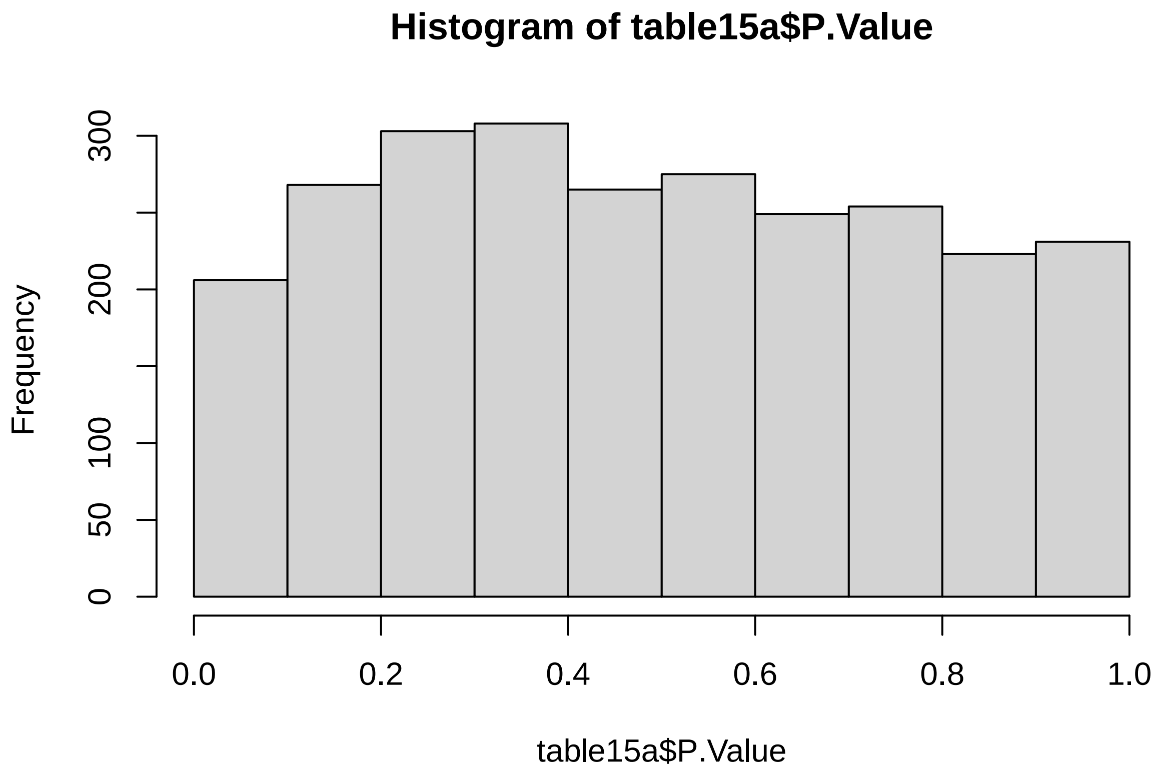

### 6 F1 male offspring

This section contains all the additive/linear and interaction/quadratic models. Including models with dietary intake as a co-variate.

#### 6.1 Protein and carbs (%)

##### 6.1.1 Additive model

```
# create design matrix
Percent_prot = as.numeric(SummarizedExperiment_males$Percent_prot)
Percent_carb = as.numeric(SummarizedExperiment_males$Percent_carb)

design16 <- model.matrix(~Percent_prot + Percent_carb)

data_sample_names <- colnames(SummarizedExperiment_males)
design_sample_names <- rownames(design16)

rownames(design16) <- data_sample_names

fit16 <- lmFit(assay(SummarizedExperiment_males), design16)
fit16a <- eBayes(fit16)

table16 <- topTable(fit16a, sort = "none", number = Inf, adjust = "fdr")

count(table16$adj.P.Val < 0.05)
```

```
## [1] 0
```

```
hist(table16$P.Value)
```

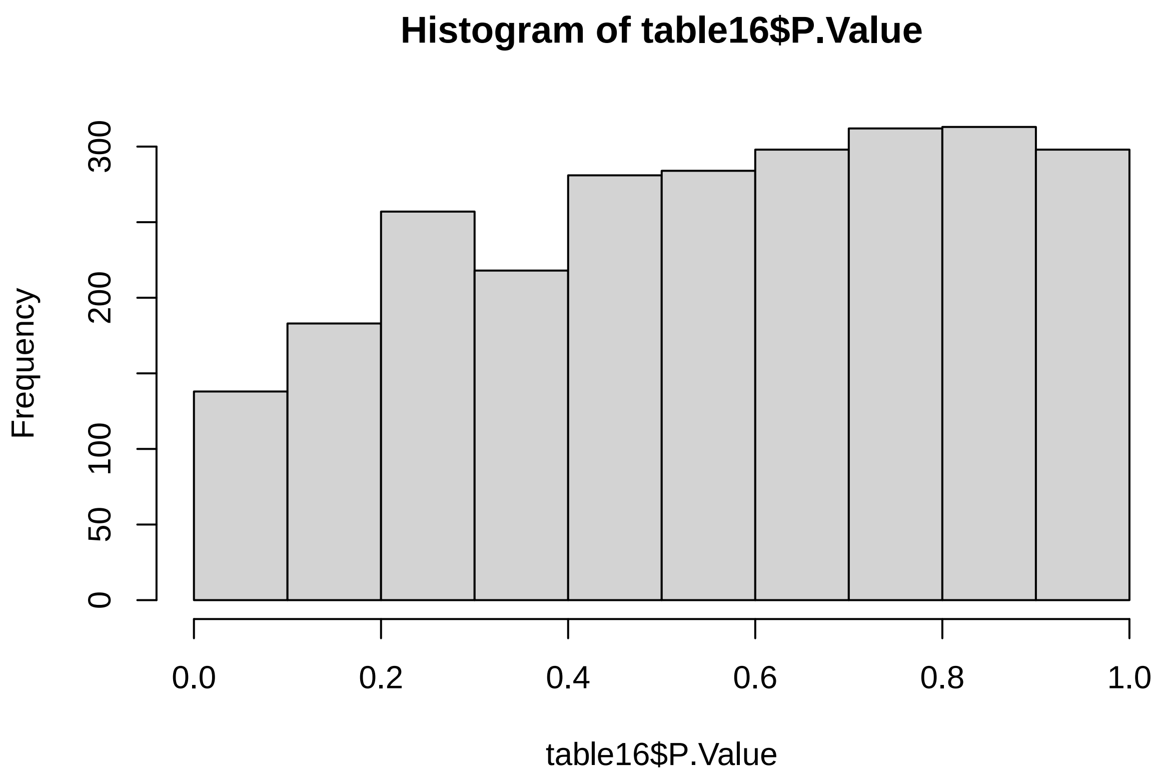

```
# 0 DE
```

###### 6.1.2 Additive model + intake

```
# create design matrix
Percent_prot = as.numeric(SummarizedExperiment_males$Percent_prot)
Percent_carb = as.numeric(SummarizedExperiment_males$Percent_carb)
intake = as.numeric(SummarizedExperiment_males$Food_intake)

design16a <- model.matrix(~Percent_prot + Percent_carb + intake)

data_sample_names <- colnames(SummarizedExperiment_males)
design_sample_names <- rownames(design16a)

rownames(design16a) <- data_sample_names

fit16a <- lmFit(assay(SummarizedExperiment_males), design16a)
fit16b <- eBayes(fit16a)

table16a <- topTable(fit16b, sort = "none", number = Inf, adjust = "fdr")

count(table16a$adj.P.Val < 0.05)
```

```
## [1] 0
```

```r
hist(table16a$P.Value)
```

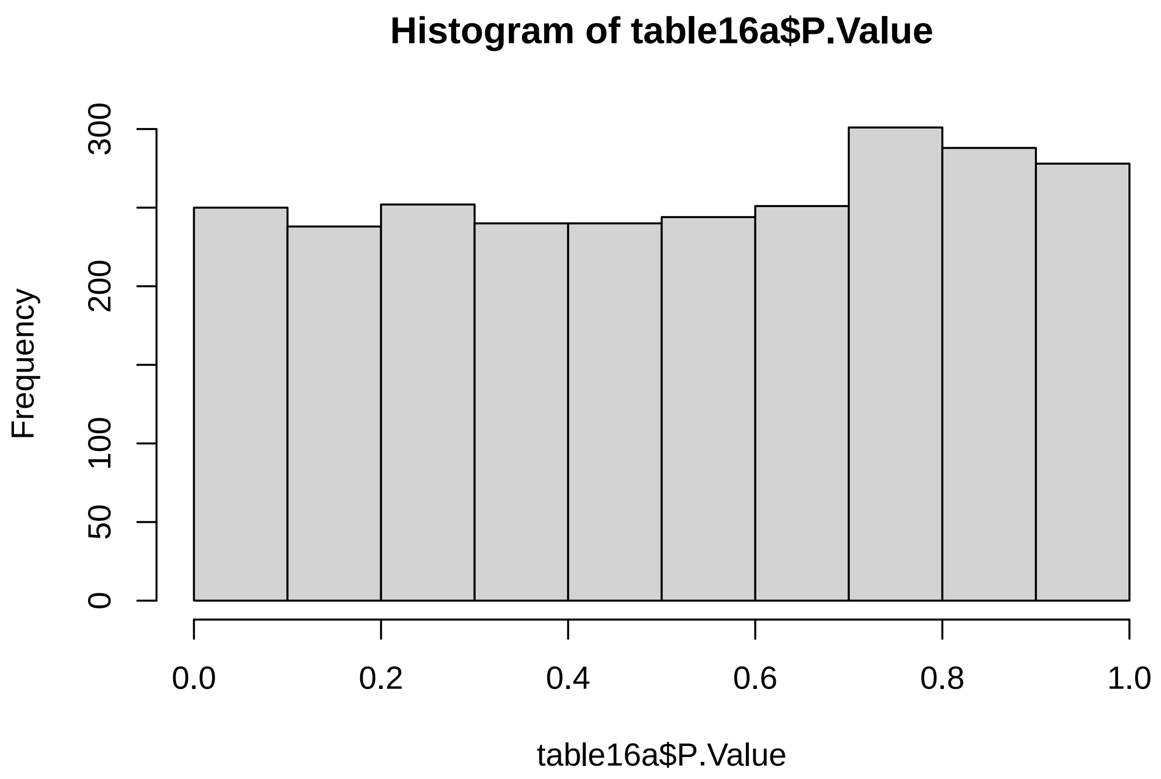

```r
# 0 DE
```

###### 6.1.3 Interaction model

```r
# create design matrix
Percent_prot = as.numeric(SummarizedExperiment_males$Percent_prot)
Percent_carb = as.numeric(SummarizedExperiment_males$Percent_carb)

design17 <- model.matrix(~Percent_prot * Percent_carb)

data_sample_names <- colnames(SummarizedExperiment_males)
design_sample_names <- rownames(design17)

rownames(design17) <- data_sample_names

fit17 <- lmFit(assay(SummarizedExperiment_males), design17)
fit17a <- eBayes(fit17)

table17 <- topTable(fit17a, sort = "none", number = Inf, adjust = "fdr")
count(table17$adj.P.Val < 0.05)
```

```
## [1] 0
```

```
hist(table17$P.Value)
```

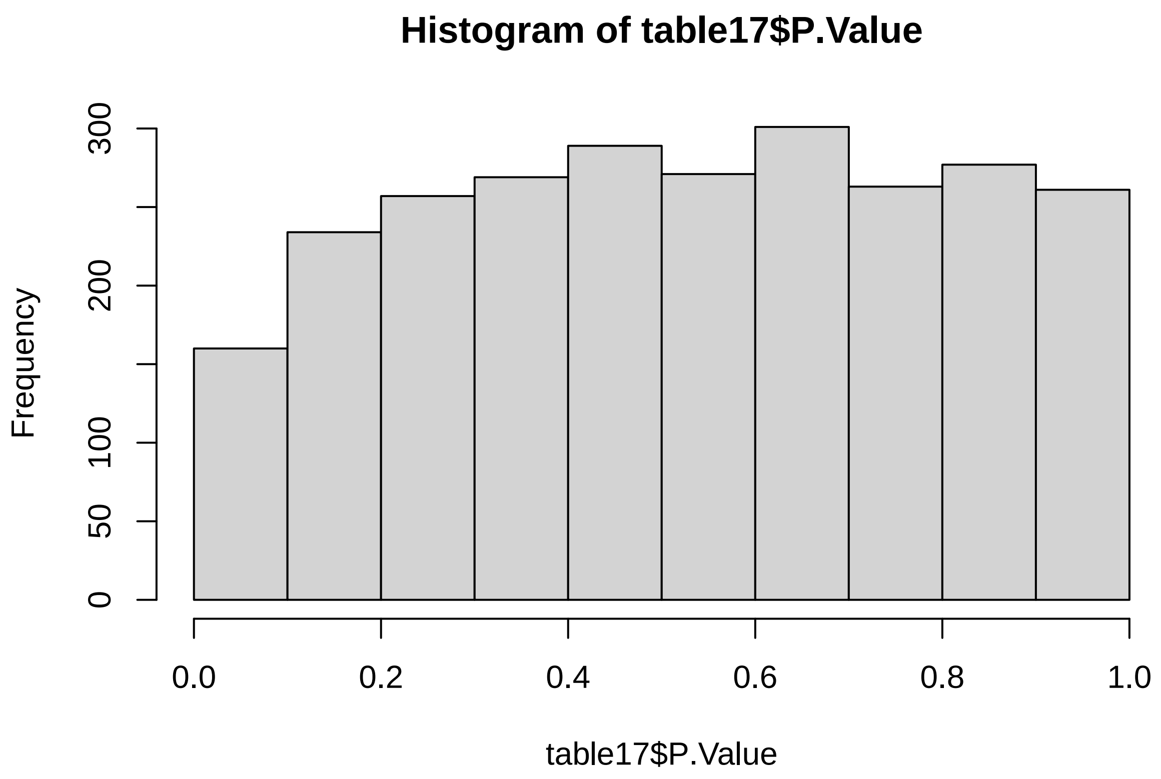

```
# 0 DE
```

###### 6.1.4 Interaction model + intake

```
# create design matrix
Percent_prot = as.numeric(SummarizedExperiment_males$Percent_prot)
Percent_carb = as.numeric(SummarizedExperiment_males$Percent_carb)
intake = as.numeric(SummarizedExperiment_males$Food_intake)

design17a <- model.matrix(~Percent_prot * Percent_carb + intake)

data_sample_names <- colnames(SummarizedExperiment_males)
design_sample_names <- rownames(design17a)

rownames(design17a) <- data_sample_names

fit17b <- lmFit(assay(SummarizedExperiment_males), design17a)
fit17b <- eBayes(fit17b)

table17a <- topTable(fit17b, sort = "none", number = Inf, adjust = "fdr")
count(table17a$adj.P.Val < 0.05)
```

```
## [1] 0
```

```r
hist(table17a$P.Value)
```

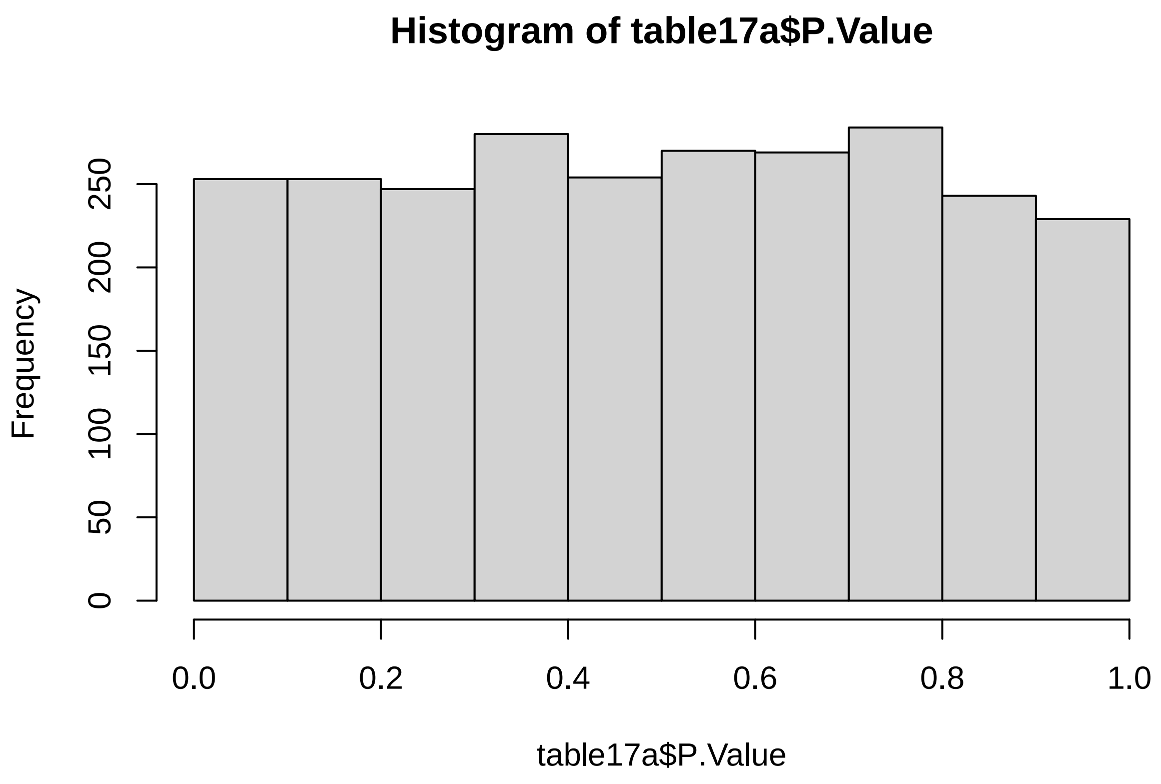

```r
# 0 DE
```

#### 6.2 Protein and fat(%)

##### 6.2.1 Additive model

```r
# create design matrix
Percent_prot = as.numeric(SummarizedExperiment_males$Percent_prot)
Percent_fat = as.numeric(SummarizedExperiment_males$Percent_fat)

design18 <- model.matrix(~Percent_prot + Percent_carb)

data_sample_names <- colnames(SummarizedExperiment_males)
design_sample_names <- rownames(design18)

rownames(design18) <- data_sample_names

fit18 <- lmFit(assay(SummarizedExperiment_males), design18)
fit18a <- eBayes(fit18)
```

```r
table18 <- topTable(fit18a, sort = "none", number = Inf, adjust = "fdr")

count(table18$adj.P.Val < 0.05)
```

```
## [1] 0
```

```r
hist(table18$P.Value)
```

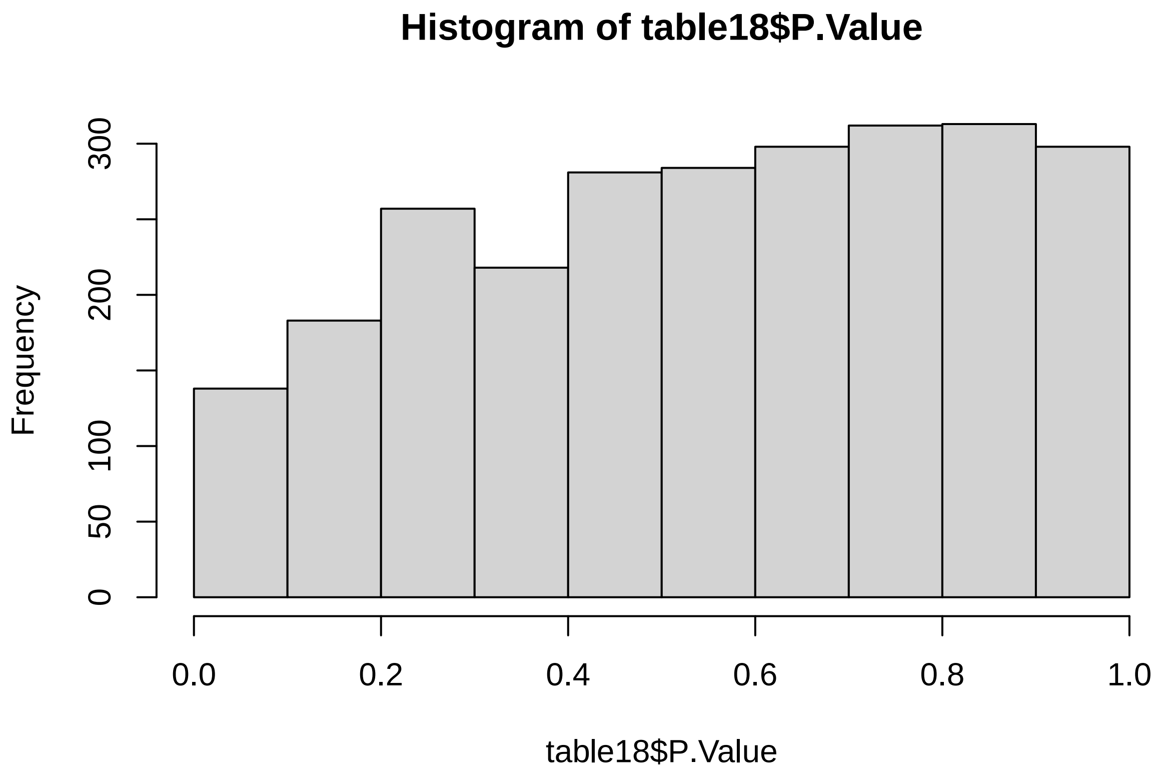

```r
# 0 DE
```

###### 6.2.2 Additive model + intake

```r
# create design matrix
Percent_prot = as.numeric(SummarizedExperiment_males$Percent_prot)
Percent_fat = as.numeric(SummarizedExperiment_males$Percent_fat)
intake = as.numeric(SummarizedExperiment_males$Food_intake)

design18a <- model.matrix(~Percent_prot + Percent_fat + intake)

data_sample_names <- colnames(SummarizedExperiment_males)
design_sample_names <- rownames(design18a)
```

```r
rownames(design18a) <- data_sample_names

fit18a <- lmFit(assay(SummarizedExperiment_males), design18a)
fit18b <- eBayes(fit18a)

table18a <- topTable(fit18b, sort = "none", number = Inf, adjust = "fdr")

count(table18a$adj.P.Val < 0.05)
```

```
## [1] 0
```

```r
hist(table18a$P.Value)
```

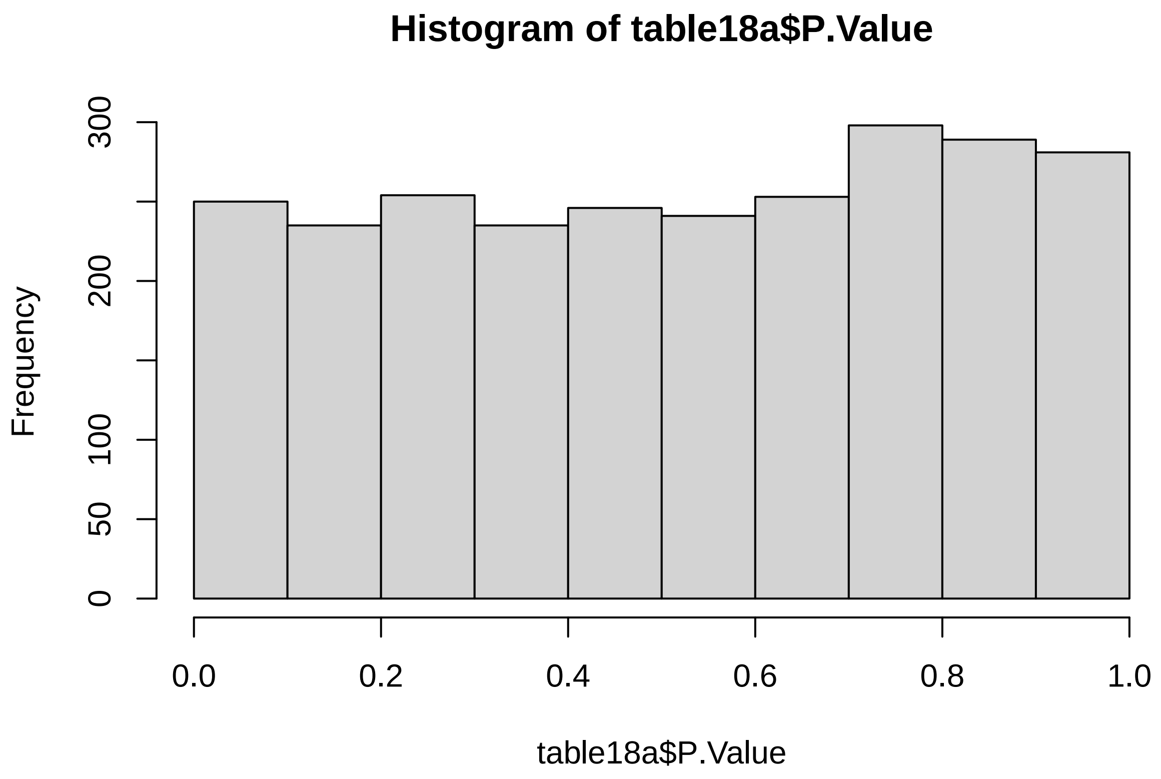

```r
# 0 DE
```

###### 6.2.3 Interaction model

```r
# create design matrix
Percent_prot = as.numeric(SummarizedExperiment_males$Percent_prot)
Percent_fat = as.numeric(SummarizedExperiment_males$Percent_fat)

design19 <- model.matrix(~Percent_prot * Percent_fat)
```

```
data_sample_names <- colnames(SummarizedExperiment_males)
design_sample_names <- rownames(design19)

rownames(design19) <- data_sample_names

fit19 <- lmFit(assay(SummarizedExperiment_males), design19)
fit19a <- eBayes(fit19)

table19 <- topTable(fit19a, sort = "none", number = Inf, adjust = "fdr")
count(table19$adj.P.Val < 0.05)
```

```
## [1] 0
```

```
hist(table19$P.Value)
```

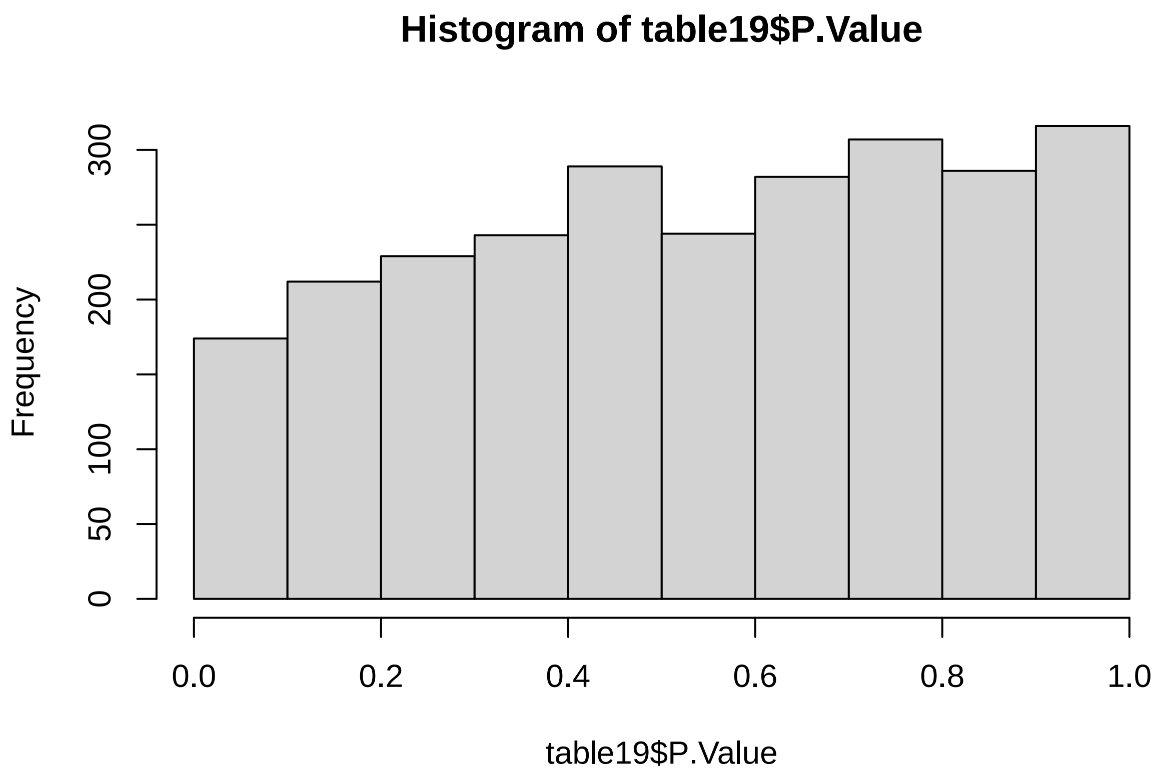

```
# 0 DE
```

###### 6.2.4 Interaction model + intake

```
# create design matrix
Percent_prot = as.numeric(SummarizedExperiment_males$Percent_prot)
Percent_fat = as.numeric(SummarizedExperiment_males$Percent_fat)
```

```r
intake = as.numeric(SummarizedExperiment_males$Food_intake)

design19a <- model.matrix(~Percent_prot * Percent_fat + intake)

data_sample_names <- colnames(SummarizedExperiment_males)
design_sample_names <- rownames(design19a)

rownames(design19a) <- data_sample_names

fit19b <- lmFit(assay(SummarizedExperiment_males), design19a)
fit19b <- eBayes(fit19b)

table19a <- topTable(fit19b, sort = "none", number = Inf, adjust = "fdr")
count(table19a$adj.P.Val < 0.05)
```

```
## [1] 0
```

```r
hist(table19a$P.Value)
```

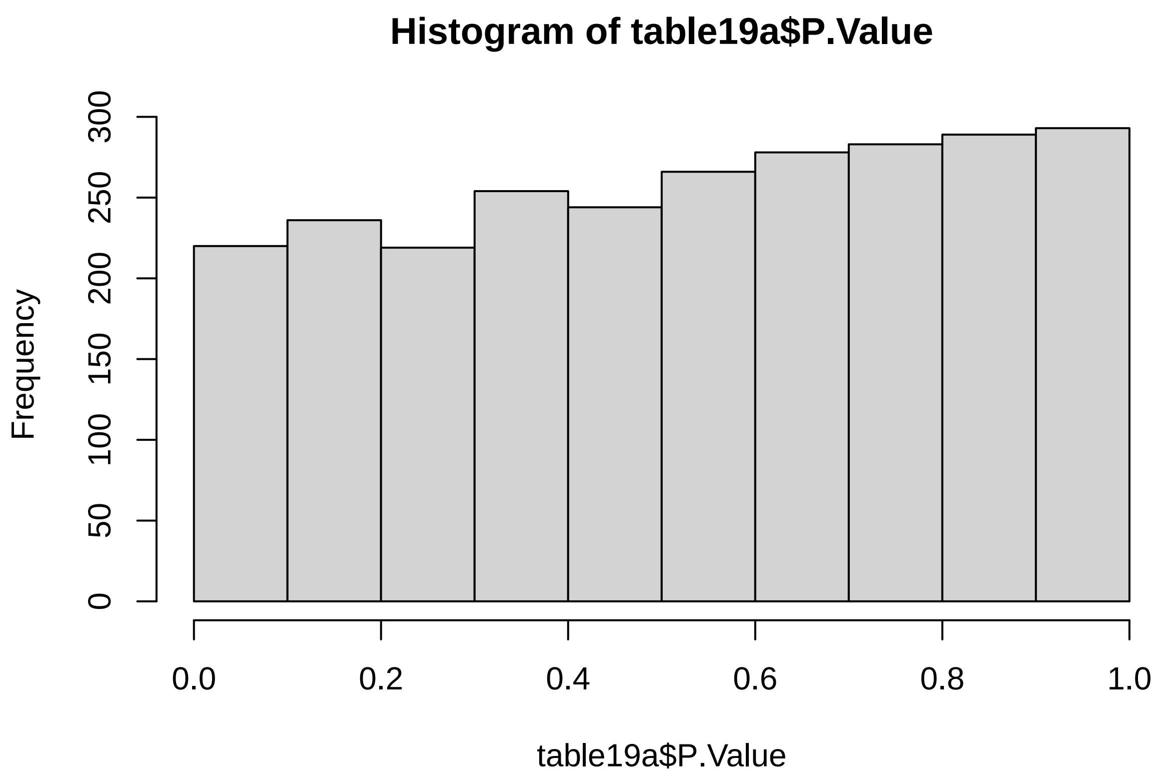

```r
# 0 DE
```

#### 6.3 Carbs and fat(%)

##### 6.3.1 Additive model

```
# create design matrix
Percent_carb = as.numeric(SummarizedExperiment_males$Percent_carb)
Percent_fat = as.numeric(SummarizedExperiment_males$Percent_fat)

design20 <- model.matrix(~Percent_carb + Percent_fat)

data_sample_names <- colnames(SummarizedExperiment_males)
design_sample_names <- rownames(design20)

rownames(design20) <- data_sample_names

fit20 <- lmFit(assay(SummarizedExperiment_males), design20)
fit20a <- eBayes(fit20)

table20 <- topTable(fit20a, sort = "none", number = Inf, adjust = "fdr")

count(table20$adj.P.Val < 0.05)
```

```
## [1] 0
```

```
hist(table20$P.Value)
```

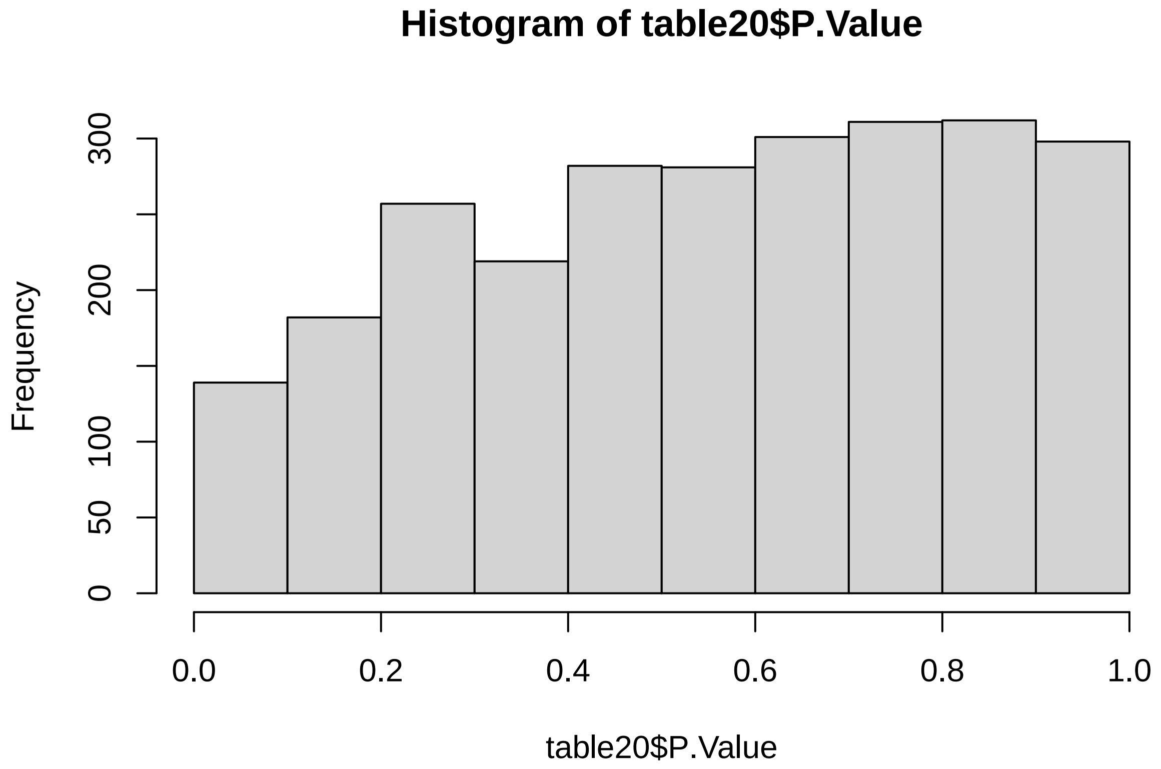

```
# 0 DE
```

##### 6.3.2 Additive model + intake

```
# create design matrix
Percent_fat = as.numeric(SummarizedExperiment_males$Percent_fat)
Percent_carb = as.numeric(SummarizedExperiment_males$Percent_carb)
intake = as.numeric(SummarizedExperiment_males$Food_intake)

design20a <- model.matrix(~Percent_carb + Percent_fat + intake)

data_sample_names <- colnames(SummarizedExperiment_males)
design_sample_names <- rownames(design20a)

rownames(design20a) <- data_sample_names

fit20a <- lmFit(assay(SummarizedExperiment_males), design20a)
fit20b <- eBayes(fit20a)

table20a <- topTable(fit20b, sort = "none", number = Inf, adjust = "fdr")

count(table20a$adj.P.Val < 0.05)
```

```
## [1] 0
```

```
hist(table20a$P.Value)
```

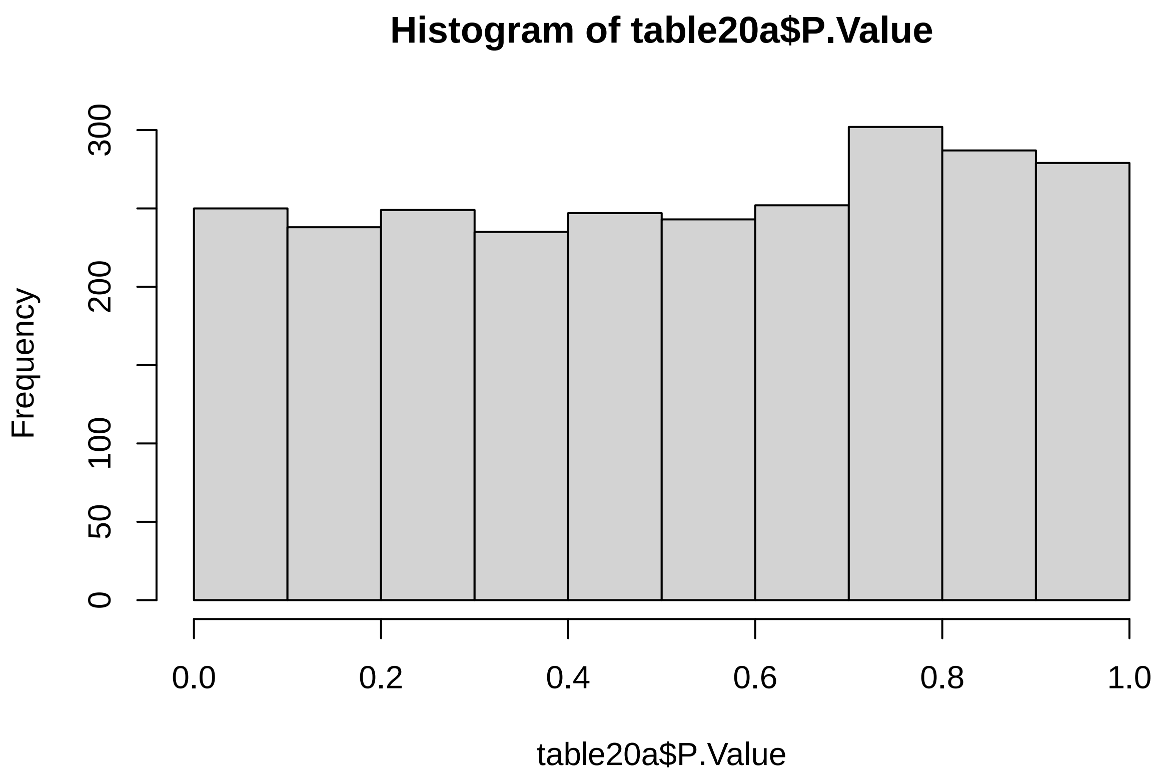

```
# 0 DE
```

##### 6.3.3 Interaction model

```
# create design matrix
Percent_fat = as.numeric(SummarizedExperiment_males$Percent_fat)
Percent_carb = as.numeric(SummarizedExperiment_males$Percent_carb)

design21 <- model.matrix(~Percent_carb + Percent_fat)

data_sample_names <- colnames(SummarizedExperiment_males)
design_sample_names <- rownames(design21)

rownames(design21) <- data_sample_names

fit21 <- lmFit(assay(SummarizedExperiment_males), design21)
fit21a <- eBayes(fit21)

table21 <- topTable(fit21a, sort = "none", number = Inf, adjust = "fdr")
count(table21$adj.P.Val < 0.05)
```

```
## [1] 0
```

```r
hist(table21$P.Value)
```

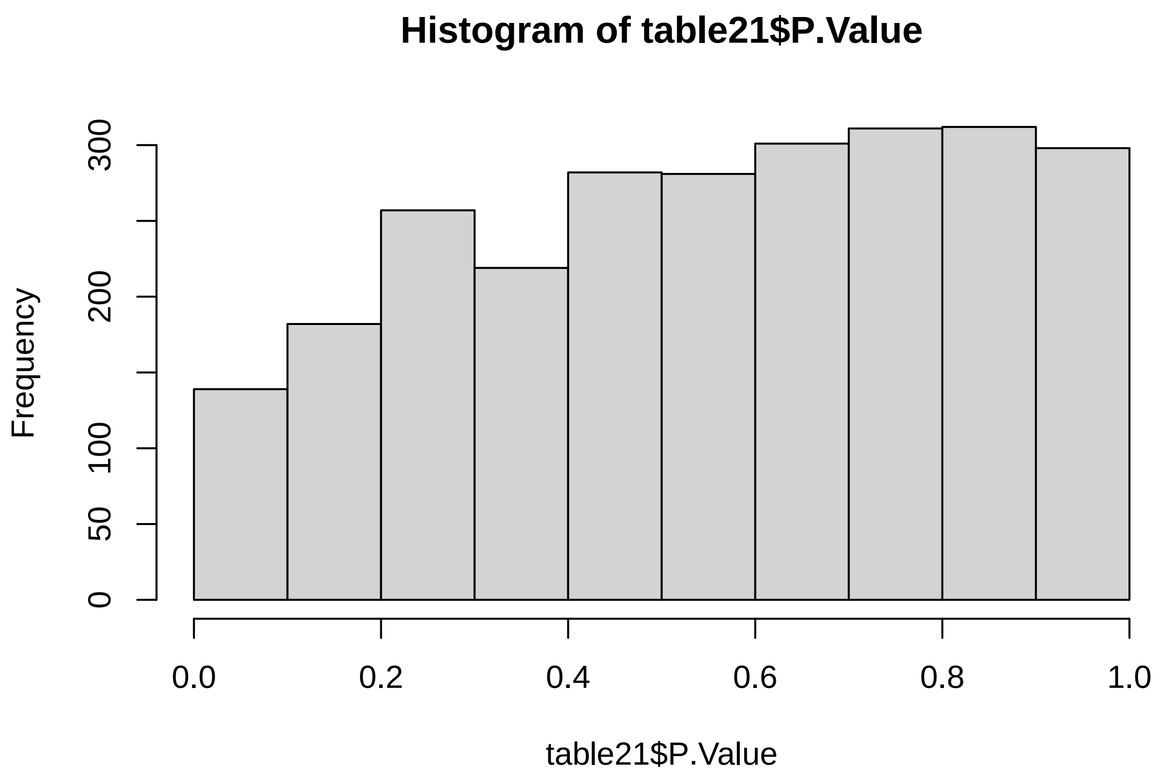

```r
# 0 DE
```

###### 6.3.4 Interaction model + intake

```r
# create design matrix
Percent_fat = as.numeric(SummarizedExperiment_males$Percent_fat)
Percent_carb = as.numeric(SummarizedExperiment_males$Percent_carb)
intake = as.numeric(SummarizedExperiment_males$Food_intake)

design21a <- model.matrix(~Percent_carb * Percent_fat + intake)

data_sample_names <- colnames(SummarizedExperiment_males)
design_sample_names <- rownames(design21a)

rownames(design21a) <- data_sample_names

fit21b <- lmFit(assay(SummarizedExperiment_males), design21a)
fit21b <- eBayes(fit21b)

table21a <- topTable(fit21b, sort = "none", number = Inf, adjust = "fdr")
count(table21a$adj.P.Val < 0.05)
```

```
## [1] 0
```

```r
hist(table21a$P.Value)
```

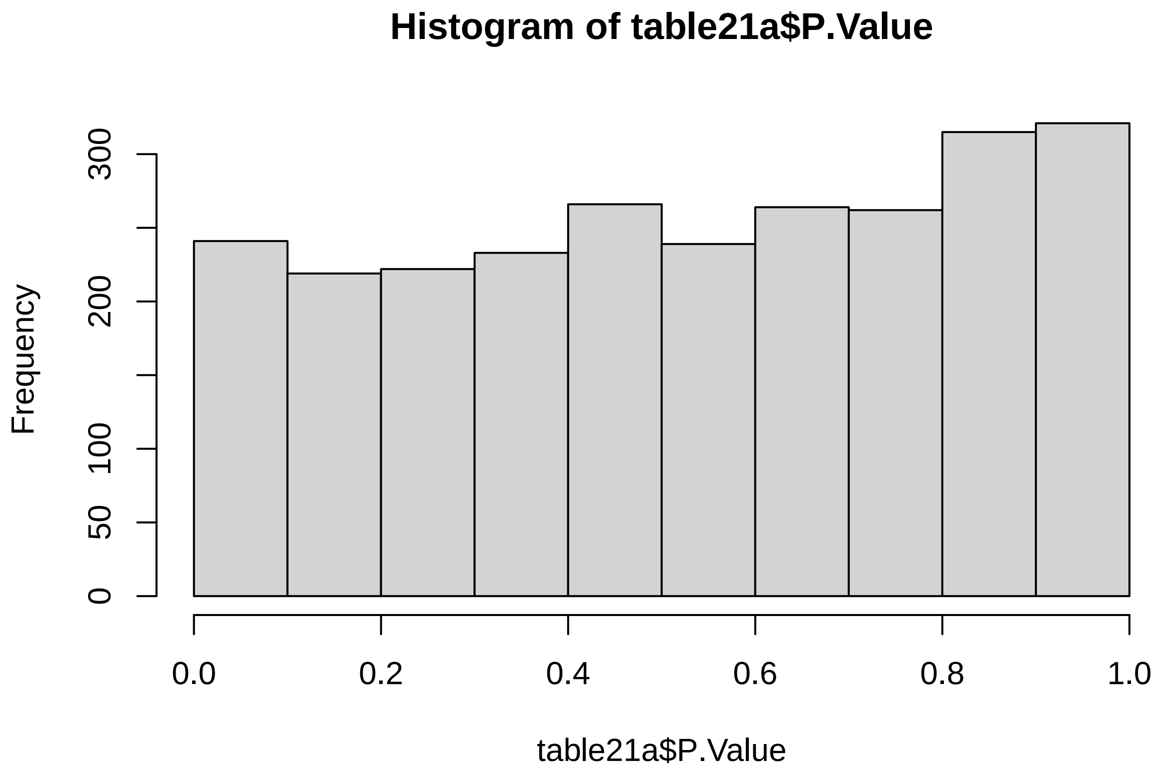

```r
# 0 DE
```

##### 6.4 Protein (%)

```r
# create design matrix
Percent_prot = log2(as.numeric(SummarizedExperiment_males$Percent_prot))

design22 <- model.matrix(~Percent_prot)

data_sample_names <- colnames(SummarizedExperiment_males)
design_sample_names <- rownames(design22)

rownames(design22) <- data_sample_names

fit22 <- lmFit(assay(SummarizedExperiment_males), design22)
fit22a <- eBayes(fit22)

table22 <- topTable(fit22a, sort = "none", number = Inf, adjust = "fdr")

count(table22$adj.P.Val < 0.05)
```

```
## [1] 0
```

```r
hist(table22$P.Value)
```

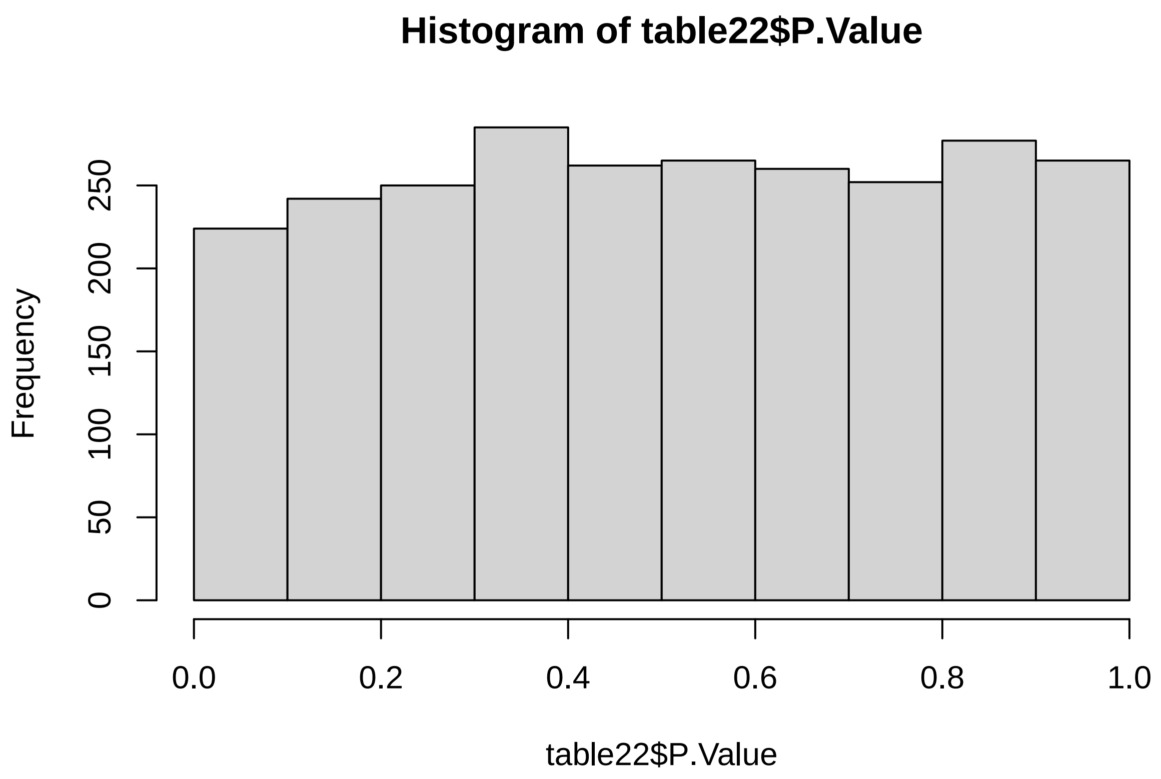

##### 6.4.1 Protein + intake

```r
# create design matrix
Percent_prot = log2(as.numeric(SummarizedExperiment_males$Percent_prot))
intake = log2(as.numeric(SummarizedExperiment_males$Food_intake))

design22a <- model.matrix(~Percent_prot + intake)

data_sample_names <- colnames(SummarizedExperiment_males)
design_sample_names <- rownames(design22a)

rownames(design22a) <- data_sample_names

fit22a <- lmFit(assay(SummarizedExperiment_males), design22a)
fit22b <- eBayes(fit22a)

table22a <- topTable(fit22b, sort = "none", number = Inf, adjust = "fdr")

count(table22a$adj.P.Val < 0.05)
```

```
## [1] 0
```

```
hist(table22a$P.Value)
```

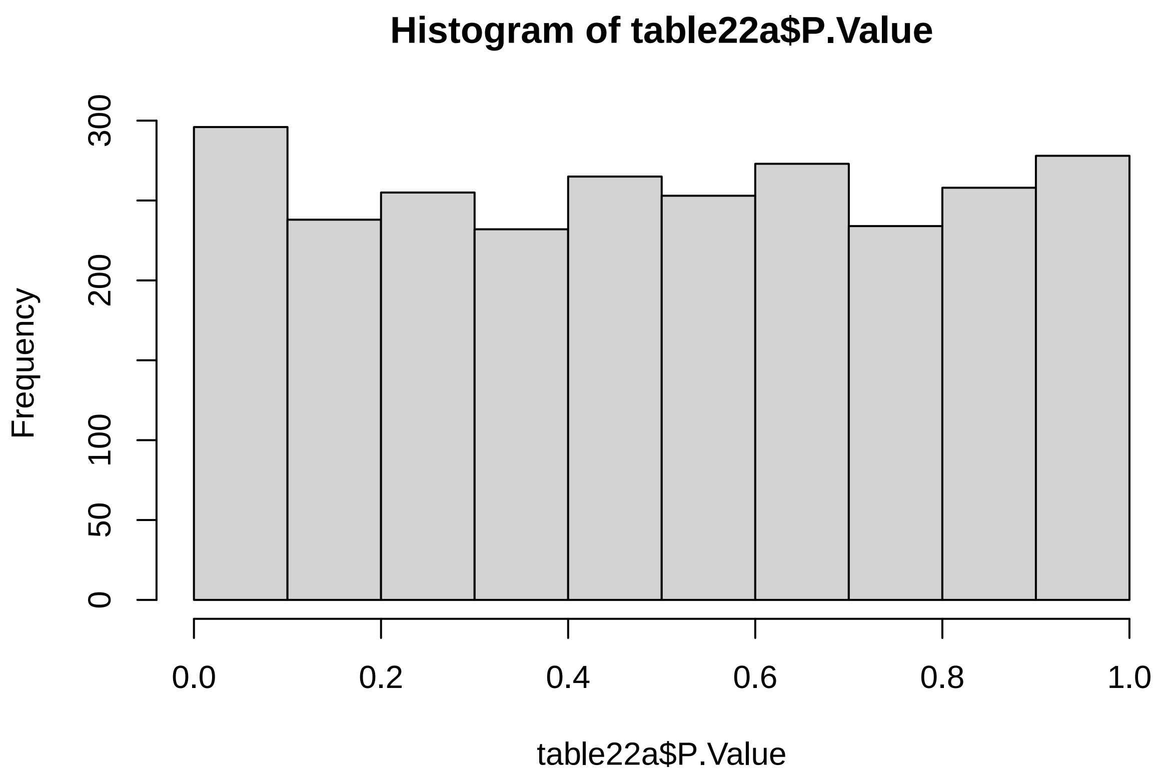

##### 6.5 Carbs (%)

```
# create design matrix
Percent_carb = log2(as.numeric(SummarizedExperiment_males$Percent_carb))

design23 <- model.matrix(~Percent_carb)

data_sample_names <- colnames(SummarizedExperiment_males)
design_sample_names <- rownames(design23)

rownames(design23) <- data_sample_names

fit23 <- lmFit(assay(SummarizedExperiment_males), design23)
fit23a <- eBayes(fit23)

table23 <- topTable(fit23a, sort = "none", number = Inf, adjust = "fdr")

count(table23$adj.P.Val < 0.05)
```

```
## [1] 0
```

```r
hist(table23$P.Value)
```

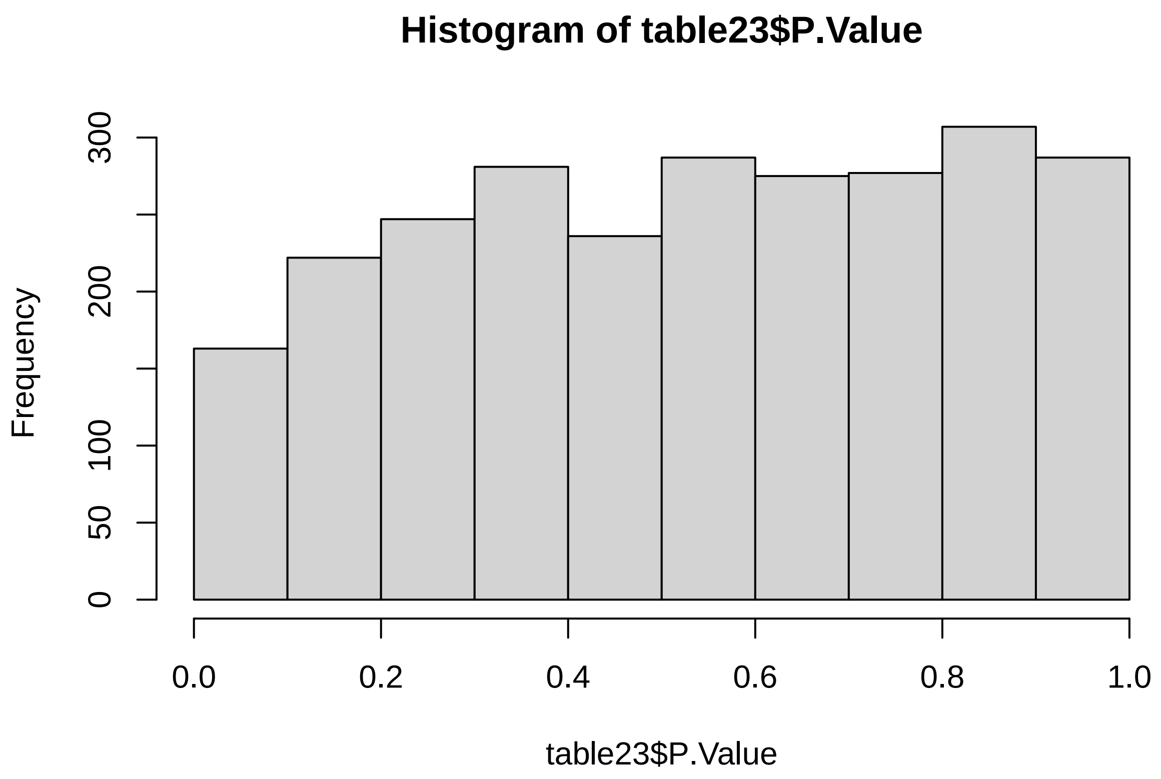

##### 6.5.1 Carb + intake

```r
# create design matrix
Percent_carb = log2(as.numeric(SummarizedExperiment_males$Percent_carb))
intake = log2(as.numeric(SummarizedExperiment_males$Food_intake))

design23a <- model.matrix(~Percent_carb + intake)

data_sample_names <- colnames(SummarizedExperiment_males)
design_sample_names <- rownames(design23a)

rownames(design23a) <- data_sample_names

fit23a <- lmFit(assay(SummarizedExperiment_males), design23a)
fit23b <- eBayes(fit23a)

table23a <- topTable(fit23b, sort = "none", number = Inf, adjust = "fdr")

count(table23a$adj.P.Val < 0.05)
```

```
## [1] 0
```

```r
hist(table23a$P.Value)
```

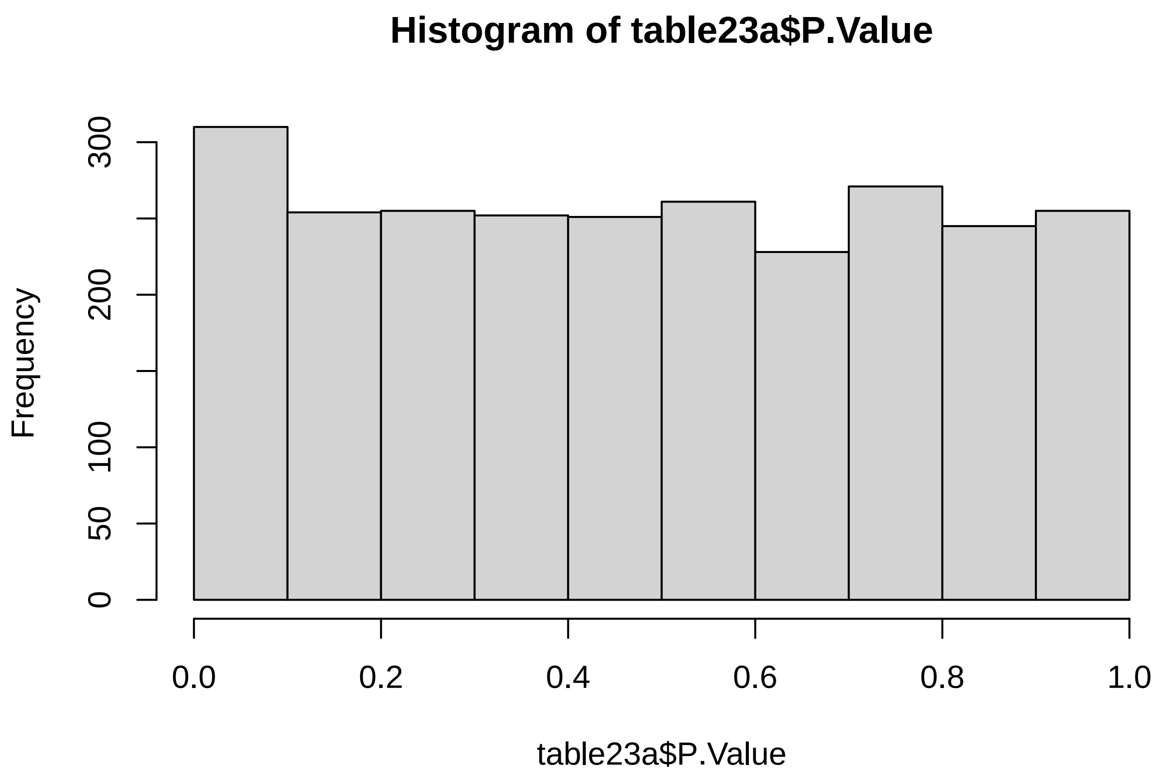

##### 6.6 Fat (%)

```r
# create design matrix
Percent_fat = log2(as.numeric(SummarizedExperiment_males$Percent_fat))

design24 <- model.matrix(~Percent_fat)

data_sample_names <- colnames(SummarizedExperiment_males)
design_sample_names <- rownames(design24)

rownames(design24) <- data_sample_names

fit24 <- lmFit(assay(SummarizedExperiment_males), design24)
fit24a <- eBayes(fit24)

table24 <- topTable(fit24a, sort = "none", number = Inf, adjust = "fdr")

count(table24$adj.P.Val < 0.05)
```

```
## [1] 0
```

```r
hist(table24$P.Value)
```

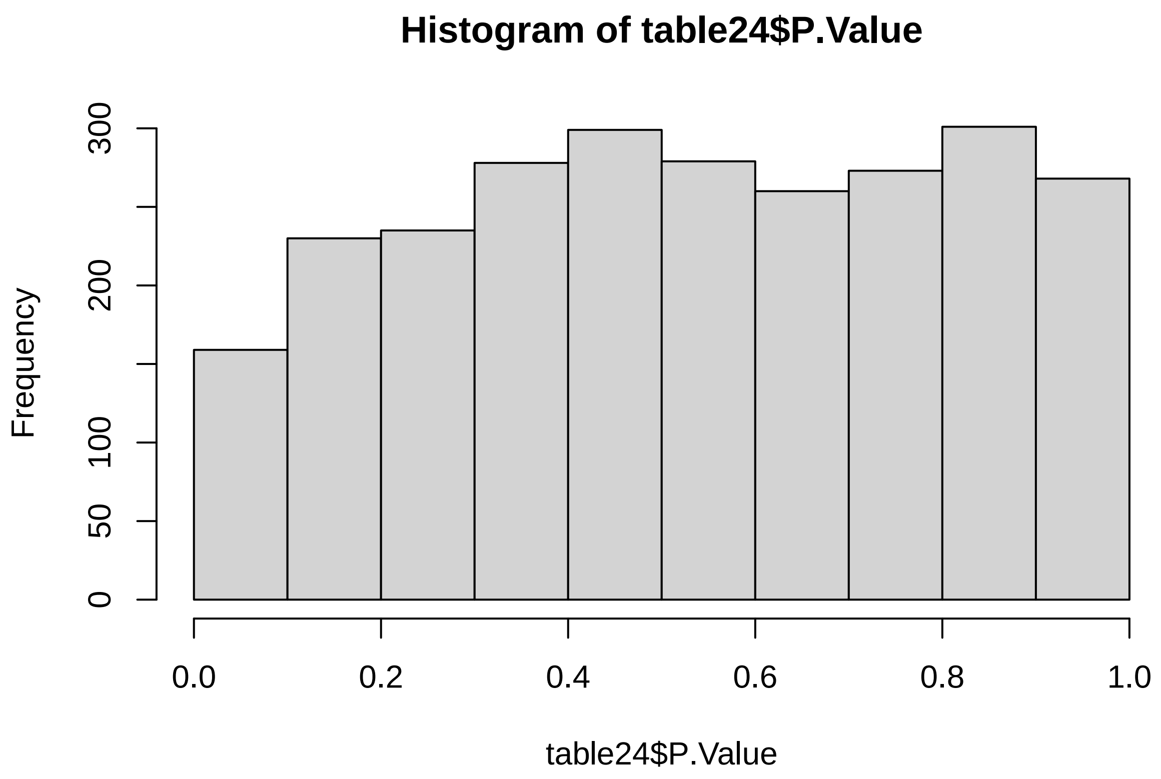

##### 6.6.1 Fat + intake

```r
# create design matrix
Percent_fat = log2(as.numeric(SummarizedExperiment_males$Percent_fat))
intake = log2(as.numeric(SummarizedExperiment_males$Food_intake))

design24a <- model.matrix(~Percent_fat + intake)

data_sample_names <- colnames(SummarizedExperiment_males)
design_sample_names <- rownames(design24a)

rownames(design24a) <- data_sample_names

fit24a <- lmFit(assay(SummarizedExperiment_males), design24a)
fit24b <- eBayes(fit24a)

table24a <- topTable(fit24b, sort = "none", number = Inf, adjust = "fdr")

count(table24a$adj.P.Val < 0.05)
```

```
## [1] 0
```

```r
hist(table24a$P.Value)
```

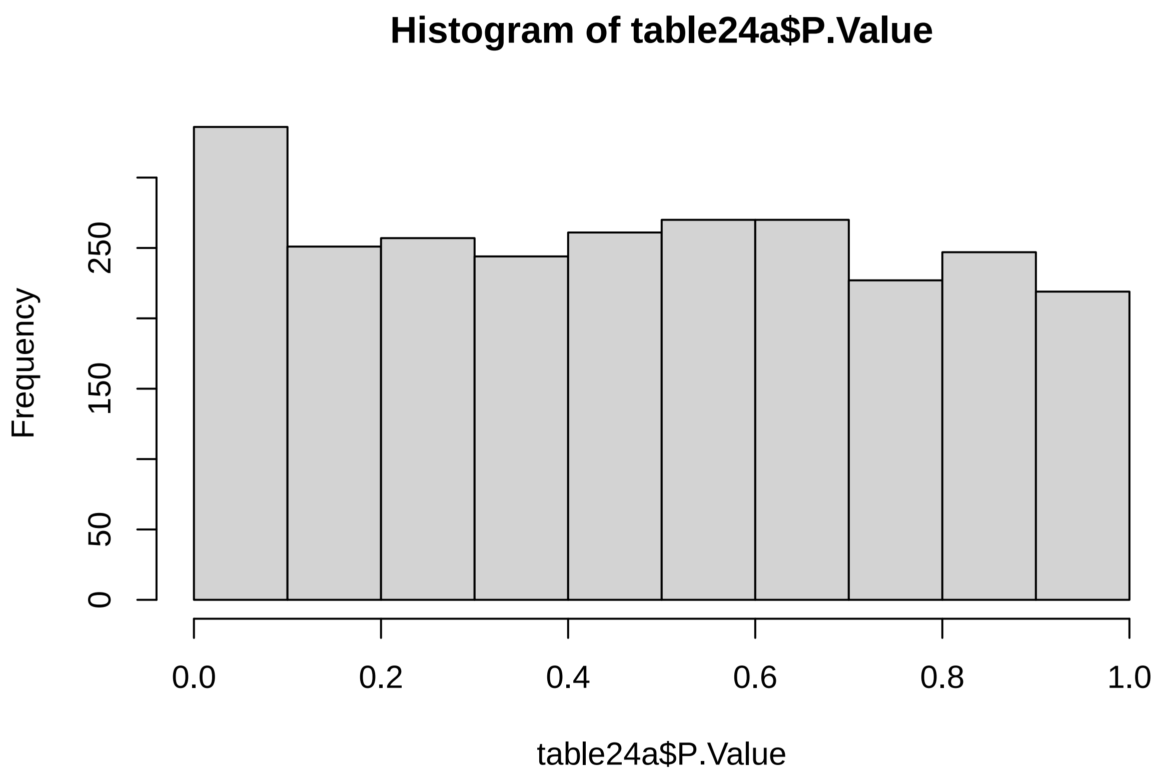
